# Chemotherapy induces myeloid-driven spatial T-cell exhaustion in ovarian cancer

**DOI:** 10.1101/2024.03.19.585657

**Authors:** Inga-Maria Launonen, Erdogan Pekcan Erkan, Iga Niemiec, Ada Junquera, María Hincapié-Otero, Daria Afenteva, Zhihan Liang, Matilda Salko, Angela Szabo, Fernando Perez-Villatoro, Matias M. Falco, Yilin Li, Giulia Micoli, Ashwini Nagaraj, Ulla-Maija Haltia, Essi Kahelin, Jaana Oikkonen, Johanna Hynninen, Anni Virtanen, Ajit J. Nirmal, Tuulia Vallius, Sampsa Hautaniemi, Peter Sorger, Anna Vähärautio, Anniina Färkkilä

**Author notes:** Faculty of Medicine and Health Technology, Tampere University, Tampere, Finland (Erdogan Pekcan Erkan). Equal contribution (Inga-Maria Launonen, Erdogan Pekcan Erkan).

## Abstract

To uncover the intricate, chemotherapy-induced spatiotemporal remodeling of the tumor microenvironment, we conducted integrative spatial and molecular characterization of 97 high-grade serous ovarian cancer (HGSC) samples collected before and after chemotherapy. Using single-cell and spatial analyses, we identify increasingly versatile immune cell states, which form spatiotemporally dynamic microcommunities at the tumor-stroma interface. We demonstrate that chemotherapy triggers spatial redistribution and exhaustion of CD8+ T cells due to prolonged antigen presentation by macrophages, both within interconnected myeloid networks termed “Myelonets” and at the tumor stroma interface. Single-cell and spatial transcriptomics identifies prominent TIGIT-NECTIN2 ligand-receptor interactions induced by chemotherapy. Using a functional patient-derived immuno-oncology platform, we show that CD8+T-cell activity can be boosted by combining immune checkpoint blockade with chemotherapy. Our discovery of chemotherapy-induced myeloid-driven spatial T-cell exhaustion paves the way for novel immunotherapeutic strategies to unleash CD8+ T-cell-mediated anti-tumor immunity in HGSC.

## Introduction

Tumorigenesis and cancer progression are dependent on the ability of malignant cells to escape the recognition and attack by the host’s immune system, involving a complex spatial interplay of cell populations. T cell exhaustion is a process in which T cells become hypofunctional upon chronic antigen stimulation; a subset of these cells can be reinvigorated by immune checkpoint blockade therapies (ICB)^1^. Cancer immunotherapies using ICB have been widely adopted in the clinical setting^2,3^. However, ovarian cancer is one of the diseases where ICB therapies have failed due to a lack of understanding of the molecular and spatial underpinnings of immune evasion^4,5^.

High-grade serous ovarian cancer (HGSC), the most common ovarian cancer subtype, is a highly aggressive disease characterized by ubiquitous deleterious mutations in *TP53^6^*, genomic instability^7^ and high tumor heterogeneity^8^. First-line neoadjuvant chemotherapy (NACT) is commonly administered to patients with unresectable disease or comorbidities^9^. Regardless of the responses to first-line platinum-based chemotherapy, treatment resistance eventually develops in the majority of the patients^10^. The spatiotemporal effects of chemotherapy on the remodeling of the tumor-immune microenvironment remain incompletely understood.

In recent years it has become increasingly evident that anti-tumor immunity plays a critical role in HGSC therapy responses and clinical outcomes^11–14^. We recently identified a phenotypically distinct TME with evidence of increased spatial immunosurveillance and distinct prognostication for the *BRCA1/2* deficient tumors^11^. Consistently, we showed that the combination of ICB with Poly-ADP Ribose Polymerase (PARP) inhibitors is effective in tumors with Homologous recombination (HR) DNA repair deficiency (HRD) and a distinct functional and spatial immune microenvironment^12^. In addition, preclinical studies have highlighted the efficacy of ICB in combination with DNA damaging therapies^15^. Previously, we have shown that NACT enriches a stressed state in cancer cells, associated with immunocompromised macrophages and CD8 T cells^16^. On the other hand, NACT has been found to potentiate the immune fitness of HGSC by inducing oligoclonal expansion of CD8+ T cells and increased infiltration of natural killer cells^17^ as well as CD4+ T cell activation^18^. Additionally, NACT has been found to skew tumor-associated macrophages (TAMs) towards a proinflammatory phenotype, which may potentiate adaptive immune responses^19^.

Macrophages are the most abundant immune and myeloid cell subtype in the tumor microenvironment (TME) of HGSC^11,20^. The heterogeneous and dynamic nature of macrophages gives rise to their multifaceted roles in the TME in maintaining tissue homeostasis^21^. TAMs co-evolve with the cancer ecosystem by initiating inflammatory cascades with the uptake of apoptotic tumor cells, antigen cross presentation, and ultimately driving tissue remodeling and tumor progression^21^. Importantly, it has been hypothesized that for ICB therapies to activate tumor-infiltrating lymphocytes (TILs), TILs must be in close proximity to antigen-presenting myeloid cells so that costimulation can take place^22^. However, how chemotherapy affects these multilayered processes is incompletely understood.

Here, we incorporate multi-omics profiling using genomics, scRNA-seq, high-dimensional whole-section tissue imaging and spatial transcriptomics from 97 patient samples capturing over 6,500,000 single-cells, and 316 spatial domains captured by single-cell and spatial molecular profiling. By combining genomic, single-cell and spatial data from patient-matched HGSC samples collected both before and after platinum-based chemotherapy, we unravel the functional effects of chemotherapy on the single-cell spatial TME. We show how the spatially resolved cell-cell interaction neighborhoods change dynamically during NACT, together with underlying transcriptional programs, resulting in myeloid-derived spatial exhaustion patterns of T cells with distinct gene expression programs. Our findings reveal new opportunities for patient stratification and targets for anti-tumor immune responses in ovarian cancer.

## Results

### Multiomic characterization of single-cell spatial TME in ovarian cancer

We prospectively collected a series of 97 spatiotemporally distinct tumor samples from 55 HGSC patients to characterize the TME dynamics during NACT (**Figure 1a-b**). The tumor samples were subjected to multi-omics profiling with WGS/WES, RNASeq, scRNA-seq, high-plex cyclic immunofluorescence (CycIF), imaging and spatial transcriptomics (**Figure 1b**).

**Figure 1:**
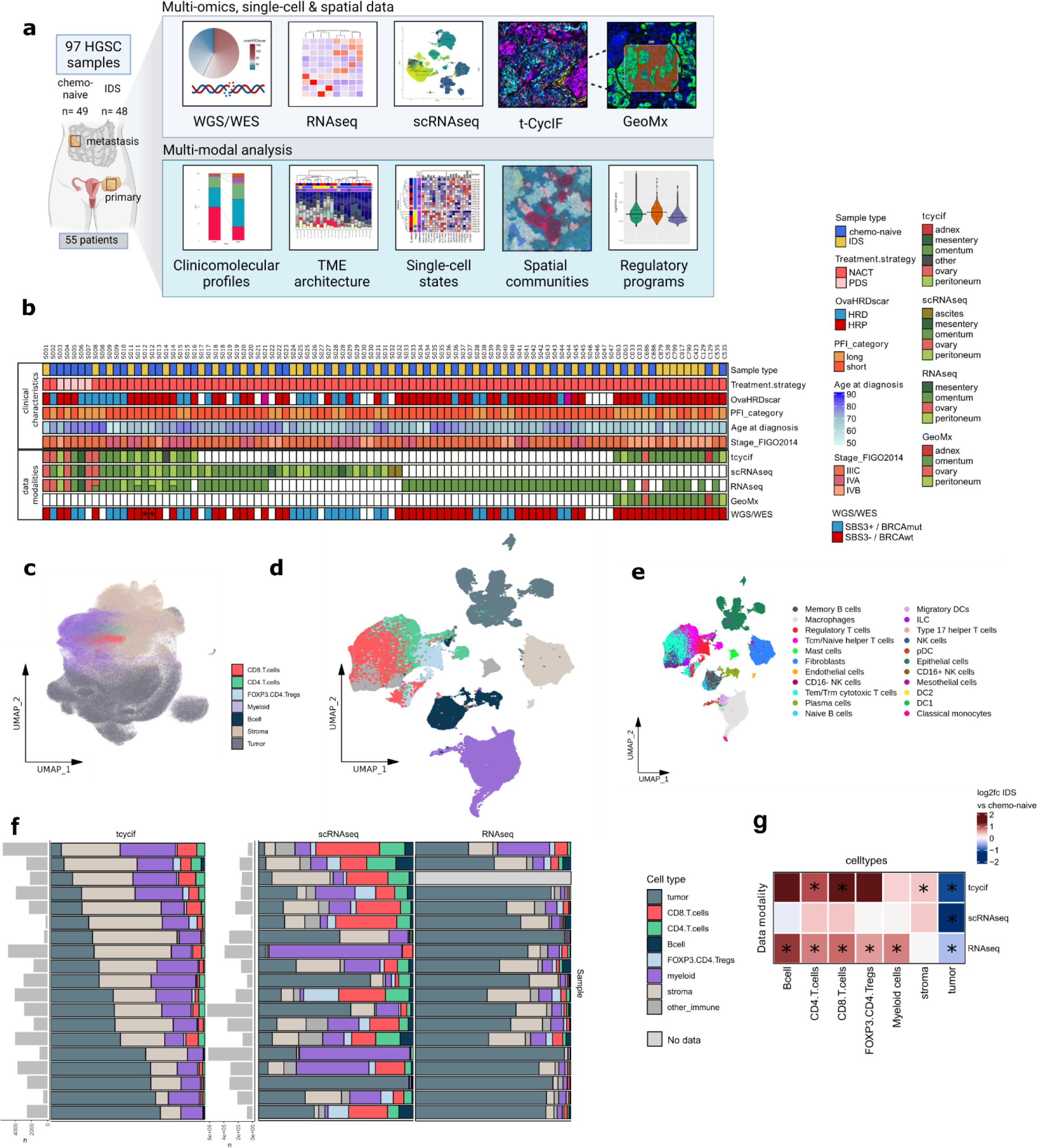
Multiomics characterization of the tumor-immune microenvironment of high-grade serous ovarian cancer. **a** Schematic overview of the data and workflow from clinicomolecular profiles, tumor-immune microenvironment architecture, single-cell states, spatial communities and programs to clinical correlations. **b** Dataset description and clinical features for 97 samples from 55 patients with HGSC. The samples which were SBS3 negative but BRCA mutated are marked with an asterisk. Patient age at diagnosis, tumor BRCA1/2 mutation or HRD status (Sig3, ovaHRDScar) were evenly distributed across the dataset and have been confirmed not to act as confounders of the results. **c** UMAP projection for cell types in t-CycIF data and **d** for scRNA-seq data. The cell type annotation colors for **d** are the same as for **c**. **e** UMAP of fine-grained cell types in scRNA-seq data. **f** Stacked bar plots of cell type proportions per sample for samples with patient- and site matched t-CycIF, scRNA-seq and RNA-seq data. Bar plots are ordered by increasing tumor cell percentage as determined from t-CycIF data. Bar plot annotations show cell type counts per t-CycIF and scRNA-seq cell types. **g** Heatmap showing the log2 fold change for the change in cell type proportions in IDS to chemo-naive samples in the different data modalities. Paired samples Wilcoxon test p-values with a value less than 0.05 for the comparison of IDS and chemo-naive samples per data modality are marked with an asterisk.

Employing whole-slide CyCIF imaging to profile the spatial architecture of the single-cell TME, we successfully identified and analyzed 6,402,172 cells across 22 tumor samples (**see Materials-Methods**). We annotated 2,684,155 tumor cells, 2,060,194 stromal cells, and 1,657,823 immune cells and further grouped them into distinct functional metaclusters (**Figure 1c, S.Fig 1a-c**). For cell type calling we developed TRIBUS, an automatic Flowsom-based clustering algorithm with prior knowledge of cell phenotypes as the input (**see Material-Methods**).

For scRNA-seq, we collected 51 HGSC tumor samples from 29 HGSC patients yielding 136,452 cells, which included 25,091 epithelial/tumor cells, 13,952 stromal cells, and 97,409 immune cells using cluster annotations for canonical markers^16^. To identify divergent immune cell subpopulations, we used the CellTypist algorithm^23^ to reliably infer the cell type labels from an established cross-tissue immune cell atlas. To facilitate the comparison of cell abundances between t-CycIF and scRNA-seq data sets, we aggregated the inferred cell type labels in the scRNA-seq data to match the higher hierarchy cell types in the t-CycIF data (**Figure 1f**). The two-dimensional uniform manifold approximation and projection (UMAP) plots of cells with respect to sample code (**S.Fig 1d**) and treatment stage (**S.Fig 1e**) are shown in the supplementary data. Further, to infer cell populations from bulk RNA-seq data, we performed Kassandra cell type deconvolution **(see Materials-Methods)**. The cell type proportions from patient-and site-matched samples had significant correlations between t-CycIF, scRNA-seq, and bulk RNA-seq data with most of the common cell types **(S.Fig 1f**, **Figure 1f)**. Tumor HRD was assessed using our ovarian cancer-specific optimized ovaHRDScar algorithm from WGS or SNP array data^24^, or single base substitution signature 3 (SBS3) indicating HRD status in altogether 87.6 % of the samples.

### Chemotherapy induces distinct TME cell population dynamics in HGSC

To assess the cell population dynamics during NACT, we integrated the cell populations from our multi-modal omics data. As an expected effect of chemotherapy, the tumor cell fraction decreased significantly after NACT in all data modalities (**Figure 1g**). Within the tumor compartment, the abundance of the proliferating epithelial phenotype decreased after NACT in line with previous results^16^ **(S.Fig 1g-h)**. Consistently^25^, we observed that chemotherapy induced a significant infiltration of CD8+ T cells and CD4+ T cells (**Figure 1g**, **S.Fig 1i**). Additionally, chemotherapy induced an increased signal for B cells, regulatory T cells and myeloid cells in bulk RNA-seq (**Figure 1f**) and in scRNA-seq we observed an increase in mast cells and innate lymphoid cells (ILCs), as well as an increase in natural killer (NK) cells. Collectively, these observations indicate an overall augmented infiltration and activation of the immune responses (**S.Fig 1i-j**).

To identify the potential reconditioning of cell states associated with chemotherapy in an unbiased manner, we used the Milo framework^26^ to uncover the cell state dynamics after NACT in scRNA-seq data. The majority of cell types with significant differences manifested bidirectional changes, meaning that in these cells, certain states were lost and new ones were introduced after chemotherapy (**Figure 2a, S.Fig 2a-b**). Innate lymphoid cells (ILCs) and mast cells were scarce prior to chemotherapy but relatively common in the IDS tumors. Importantly, macrophages, being the most common immune cell population in the HGSC TME, exhibited a substantial emergence of novel cell states in the IDS specimens indicating enhanced versatility of macrophage populations after NACT. The cell states significantly enriched in macrophages after chemotherapy expressed high levels of genes involved in key biological processes regulating immune modulation (Supplementary Table 1), with TNF⍺ and NFκB as the two most significantly enriched pathways (**S. Fig 2c)**. For cytotoxic CD8+ T cells and naïve CD4+ helper T cells, no pre-existing states were lost but new states were introduced during chemotherapy, suggesting that also their phenotypes were more versatile after chemotherapy. On the contrary, the majority of differentially abundant epithelial cell states were identified in the chemo-naïve samples, suggesting that in addition to the number of cells, the variety of cancer cell states diminishes after chemotherapy (**Figure 2a, S.Fig 2a-b**).

**Figure 2:**
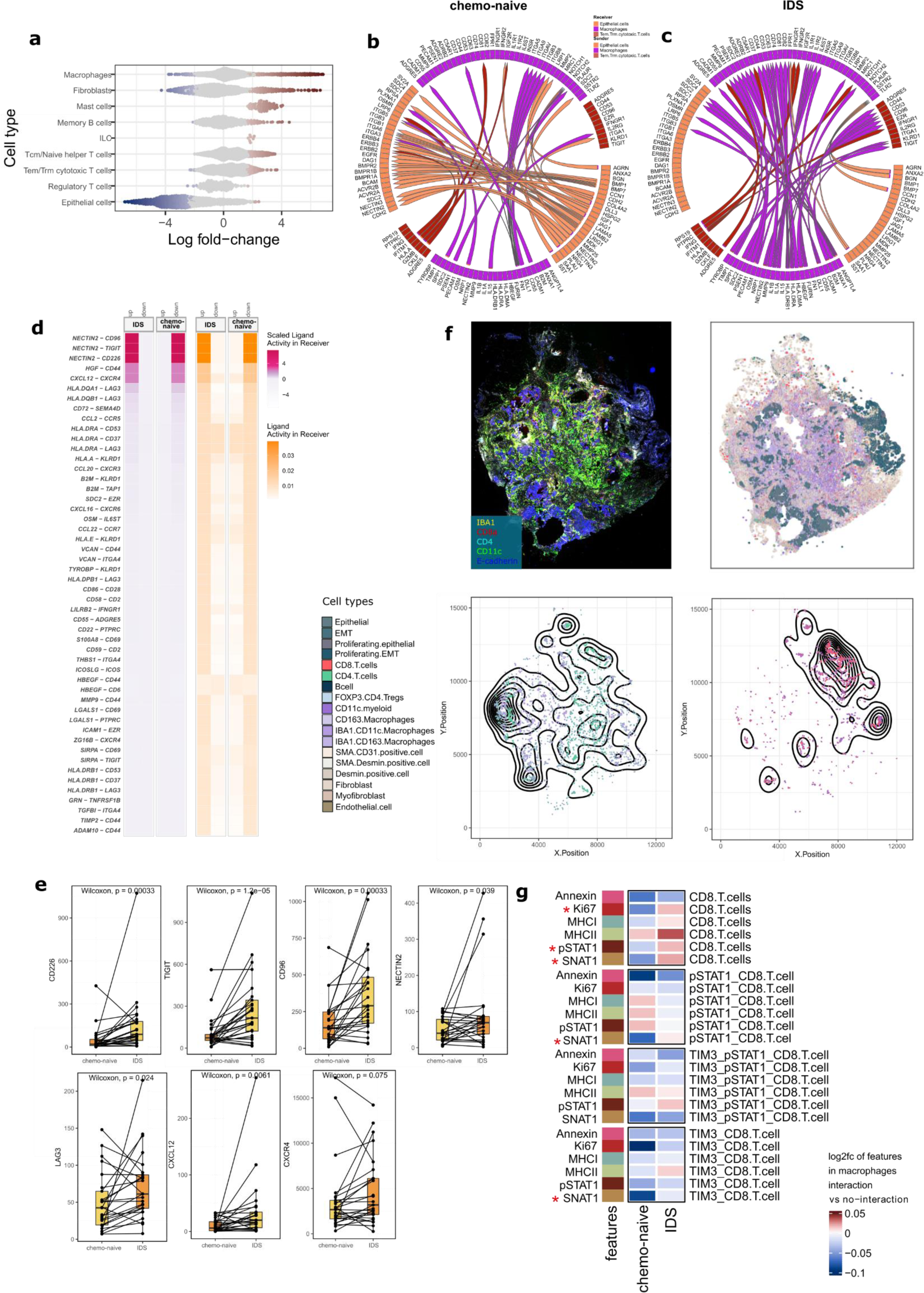
Chemotherapy-associated dynamic changes in cell type proportions and ligand-receptor interactions. **a** Beeswarm plot showing the enrichment of cell state neighborhoods in chemo-naive and IDS samples. Light gray color denotes cell state neighborhoods with an FDR value > 0.05. **b** Circos plot depicting the top 50 ligand receptor interactions in chemo-naive samples and **c** in IDS samples **d** MultiNicheNet ligand-receptor interaction analysis showing the top 50 different ligand receptor interactions between macrophages and CD8+ T cells, their ligand activity, and scaled ligand activity **e** The change in expression of selected ligands and receptors from MultiNicheNet analysis in immune deconvoluted bulkRNAseq data from chemo-naive to IDS samples. In boxplots, the black horizontal lines represent the sample medians, the boxes extend from first to third quartile and the whiskers indicate the values at 1.5 times the interquartile range. Individual dots represent values per sample. **f** Spatial heterogeneity in cell-cell interactions depicted in t-cycif image and associated xy-plot with cell types colored, and contour plots of CD4+T-cell interactions with IBA1+CD11c+ macrophages as well as CD8+ T cell interactions with CD11c+ myeloid cells. **g** Heatmap with log2 fold change of expression values (color of the heatmap) of functional markers in macrophages interacting with CD8+T-cells as compared to without interacting CD8+T-cells in chemo-naive and IDS samples. Paired Wilcoxon test p-values <0.05 when comparing values of IDS and chemo-naive samples are marked with a red asterisk.

### Chemotherapy induces distinct cellular ligand-receptor crosstalk

Given the predominant infiltration of macrophages, increased CD8+T-cell infiltration, and the chemotherapy-induced phenotypic populations, we next focused specifically on these two cell types. To assess how intercellular crosstalk changes after NACT between these cell types and cancer cells, we used the MultiNicheNet framework^27^ to perform ligand-receptor interaction analysis (**see Materials-Methods**), utilizing scRNA-seq data from intra-abdominal metastatic, paired chemo-naive and IDS samples from the same patients. Cancer cell interactions involving cellular adhesion, integrins and Notch signaling dominated the ligand-receptor crosstalk enriched in chemo-naive samples (**Figure 2b**). In the IDS samples, conversely, the dynamics were shifted towards macrophage-dominated signaling (**Figure 2c**). This shift mirrors the observed cell state changes in the differential abundance analysis (Milo). The cancer cells that exhibit decreased cell state versatility following chemotherapy demonstrate a reduced potential for interactions in the IDS samples. Conversely, the emergence of new states in both macrophage and cytotoxic CD8+ T cell populations facilitates a more diverse interaction network across the immune compartment. The strongest induced interaction from macrophages towards cytotoxic CD8+ T cells was NECTIN2 binding to competing receptors: activating receptor CD226^28,29^ and repressive receptors CD96 and TIGIT (**Figure 2d, S.Fig 2d**). While *NECTIN2-CD226* interaction in the IDS samples may contribute to the increase in CD8+ T cell infiltration after NACT, the interactions towards TIGIT and CD96 have the potential to limit the cytotoxic responses by CD8+ T cells, therefore hampering NACT anti-tumor immunity despite T cell infiltration. Furthermore, IDS samples displayed increased MHC II to CD8+ T cell LAG3 interaction that is associated with T cell exhaustion during prolonged antigen stimulation in cancer or chronic viral infection^30,31^. Ligand-receptor analysis further displayed increased signaling potential from macrophages towards the T cell adhesion and homing receptors CXCR4 and CD44 in the IDS specimens, providing further explanation for the increased T cell infiltration in chemotherapy-challenged tumors. Although the ligands HGF and CXCL12 have better defined roles in stromal cells such as cancer-associated fibroblasts^32^, our analysis suggests that macrophages also play an important role in the IDS-induced chemoattraction of T cells.

We further assessed the chemotherapy-induced expression of both the repressive and adhesive receptors identified in the ligand-receptor analysis by using bulk RNA-seq from 25 patients with 50 paired intra-abdominal metastases, after data was immune-deconvoluted with PRISM33 to minimize expression signals from fibroblasts and cancer cells (**Figure 2e**). In line with the ligand-receptor analysis, the immune compartment derived expression of the repressive *TIGIT, CD96* and *LAG3* and adhesive *CD226* were significantly induced by chemotherapy, as well as the expression of the *CXCL12* chemokine. Furthermore, chemotherapy induced the expression of distinct transcription factors in immune-deconvoluted RNA-seq, promoting T-cell exhaustion (*IRF4*)^34^, inflammation and immunity (*SPI1*, *FOXO3*, *CEBPA*) (**S. Fig 2e**).

We next investigated whether macrophages and T-cells were also spatially co-localized, using our t-CycIF single-cell spatial data. Interestingly, the cellular interactions displayed uneven spatial distribution, and the cell-cell interactions between CD8+ or CD4+ T cells with macrophages formed distinct spatial gradients across the tissues (**Figure 2f**). We then probed the functional phenotypes of the macrophages interacting with distinct CD8+ T cell states to assess whether functional immunophenotypes affect the spatial cellular crosstalk. Remarkably, we observed that the functional marker expressions were significantly associated with distinct spatial interactions between the cell types, and that these immunophenotypic interactions dynamically changed during NACT (**Figure 2g**). The heatmap in **Figure 2g** shows the change of marker expression in macrophages according to the presence (or absence) of spatial interactions with distinct CD8+ T cell subpopulations in either chemo-naive or IDS samples (**See Materials-Methods**). MHC II in macrophages exhibited the highest dependency on the CD8+T-cell interaction, with a prominent expression particularly in the IDS samples, indicating potential priming of the pSTAT1 and TIM3 negative CD8+T-cells by macrophages via MHC II. Interestingly, chemotherapy enhanced the spatial interactions of macrophages with an activated phenotype (SNAT1, STAT1, and Ki67 expression) with CD8+ T cells (**Figure 2g**). Furthermore, chemotherapy enhanced the spatial interactions of metabolically active (SNAT1 expression) macrophages and interferon-activated pSTAT1+CD8+ T cells. Interestingly, none of these macrophage states were specific to interactions with terminally exhausted TIM3+CD8+T-cells. These data suggest myeloid-driven spatial regulation of CD8+T-cell states, and highlight their dynamics during chemotherapy.

Similar analysis for tumor-CD8+ T cell interactions showed the highest spatial association of Ki67 expressing tumor cells and effector-like TIM3+pSTAT1+CD8+T-cells, indicating enhanced spatial immunosurveillance towards proliferating tumor cells in HGSC consistent with our previous observations^11^ (**S. Fig 2f**). Additionally, antigen presentation (MHC I-II), glutamine metabolism (TAZ, SNAT1) and interferon activation (pSTAT1) were higher in tumor cells interacting with distinct CD8+ T cell populations (**Figure 2g**). However, the interaction phenotypes between tumor cells and CD8+ T-cells remained unaffected by chemotherapy, indicating a more prominent role for macrophages in shaping the spatial and functional changes of CD8+ T-cells during NACT, as also supported by the ligand-receptor analysis. Taken together, our results point towards distinct spatial crosstalk between macrophages and CD8+ T cells, resulting in enhanced activation and exhaustion of CD8+ T cells during NACT.

### Chemotherapy drives spatial interaction dynamics in the TME

Taking advantage of the spatial information in the t-CycIF data, we annotated 18 distinct spatial microcommunities or recurrent cellular neighborhoods (RCNs) by clustering the cell type neighborhood matrix (**See Materials-Methods**). The resultant RCNs included six tumor cell-dominated neighborhoods (RCN1-6), six stromal cell-dominated neighborhoods (RCN8-13), five immune cell-dominated neighborhoods (RCN14-18) and the tumor-stroma interface (RCN7) consisting of tumor, stromal, and immune cell subtypes (**Figure 3a**). Most of the immune rich RCNs primarily consisted of myeloid cells; however, one immune cell RCN was specifically enriched in T-cells forming hubs with CD8+ and CD4+ T cells (RCN17). After RCN17, the highest T-cell infiltration was occurring in the myeloid-rich neighborhoods containing RCN16, RCN18 and RCN^14^. (**S.Fig 3a**). The RCN with the highest diversity in terms of cell types and marker expressions was the neighborhood consisting of tumor, stromal and immune cells (RCN7), while the RCN with the lowest cellular diversity was the neighborhood primarily consisting of epithelial tumor cells (RCN3) (**Figure 3a**). Clustering of the RCN compositions showed distinct spatial TME neighborhood compositions induced by chemotherapy, while we did not see significant clustering based on the HRD status **(Figure 3b)**. The IDS samples contained significantly decreased proportion of RCN1 and RCN2 corresponding to neighborhoods enriched in proliferating epithelial and epithelial cells, and increased proportions of RCN12 and RCN17 corresponding to myofibroblast and CD8+ T cell and CD4+ T cell enriched neighborhoods, respectively **(S.Fig.3b)**.

**Figure 3.**
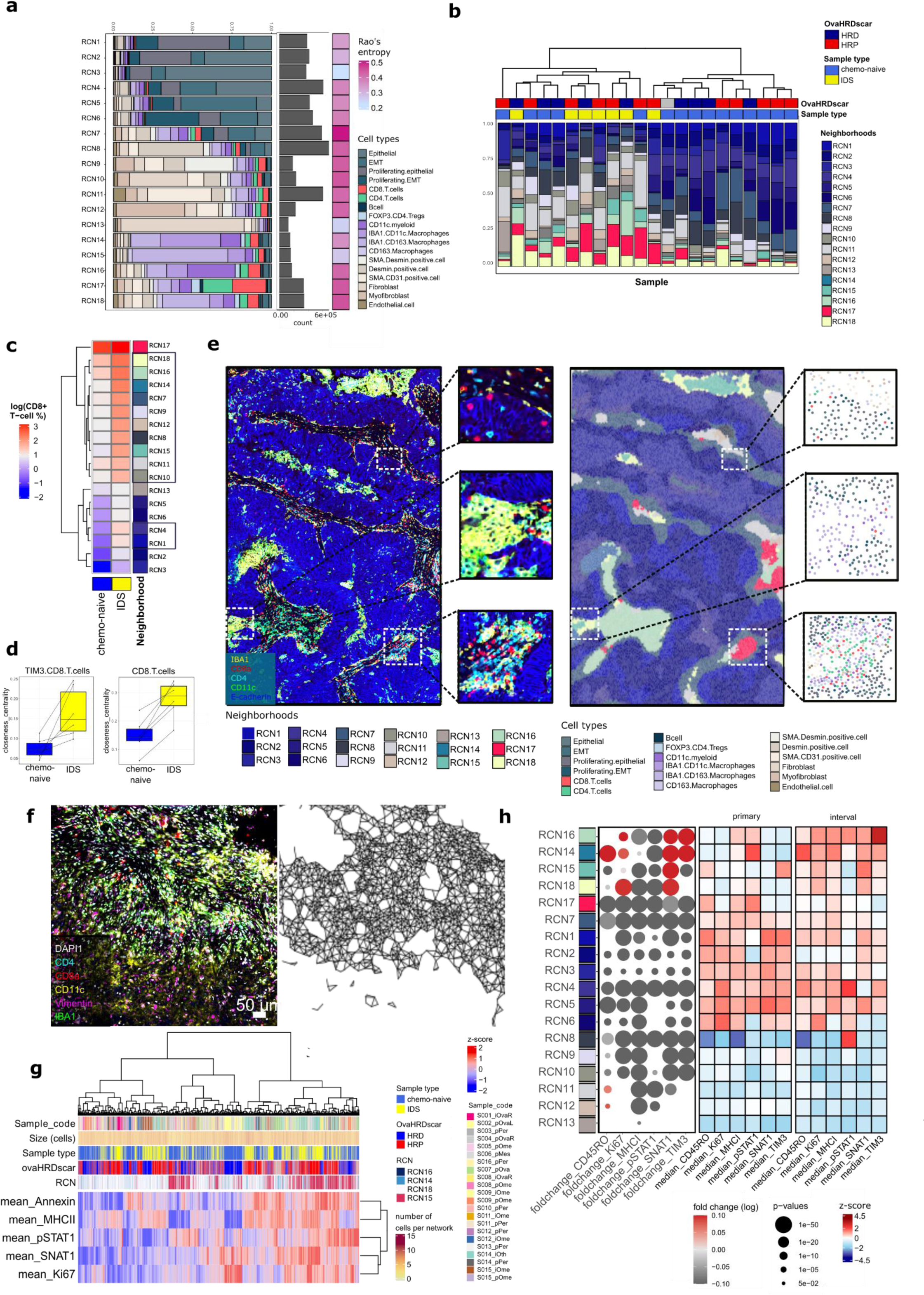
Recurrent cellular neighborhoods uncover heterogeneous cellular architecture. **a** Stacked barplot of the cell type composition of each recurrent cellular neighborhood (RCN). The gray barplot on the right represents the total cell type count in each RCN. The final annotation on the right represents Rao’s entropy of each neighborhood. **b** Bar plot ordered by hierarchical clustering of the RCN proportions per sample clusters them into two groups. Annotations include the HRD status (red=HRP, blue=HRD) and treatment stage (blue=chemo-naive, yellow=IDS). **c** The heatmap of the logarithmic CD8+ T cell proportion per RCN separately in chemo-naive and IDS samples shows a more even CD8+ T cell infiltration pattern in IDS samples. Rows highlighted with black squares have a p-value < 0.05 in the FDR-corrected Wilcoxon paired test between chemo-naive and IDS samples. **d** Box plots showing higher closeness centrality score representing spatial dispersion for TIM3+CD8+T-cells as well as CD8+ T cells in IDS samples as compared to chemo-naive. In boxplots, the black horizontal lines represent the sample medians, the boxes extend from first to third quartile and the whiskers indicate the values at 1.5 times the interquartile range. Individual dots represent values per sample. **e** Representative image showing a t-CycIF image on the left, and corresponding RCN neighborhoods (large panel) and individual cell types (small boxes) on the right. **f** Representative t-CycIF image of myelonets on the left, and cells belonging to myelonets connected using Delaunay triangulation on the bottom (RCN14). **g** Heatmap with hierarchical clustering of the mean functional marker expression per individual myelonet with the patient of origin, HRD status, number of cells per network, and sample type as annotations **h** Dot plot and heatmaps showing differences in CD8+ T cell functional states across the RCN neighborhoods. The dot plot shows the log2 fold change in the mean marker expression in IDS to chemo-naive samples with FDR-corrected Wilcoxon p-values as the size of the dot, while heatmaps on the right show the column-wise z-score of marker expressions in CD8+ T cells in RCNs in chemo-naive and IDS samples, respectively.

Remarkably, chemotherapy elicited a substantial alteration in the spatial distribution of CD8+ T cells within the RCNs **(Figure 3c**). In the chemo-naive samples, CD8+ T cells predominantly occupied the T-cell hubs (RCN17). In contrast, chemotherapy induced a significantly augmented infiltration of CD8+T cells across various RCNs in the IDS samples (**Figure 3c**). The increased infiltration was notably observed towards other immune neighborhoods, stromal neighborhoods (excluding fibroblast-rich RCNs), and neighborhoods abundant in proliferating epithelial cells, as well as those with features of EMT and epithelial cells **(Figure 3c)**. Notably, chemotherapy distinctly induced infiltration of exhausted TIM3+ and not pSTAT1 activated CD8+ T-cells into the myeloid neighborhoods **(Figure 3d, S. Fig 3c**), suggesting that T-cell exhaustion is spatially controlled. A similar trend was observed also for CD4+ T cells and FOXP3+CD4+ Tregs, although these changes did not reach statistical significance (**S.Fig 3d-e**). Altogether, our findings indicate that chemotherapy induces a transition from a spatially constrained immunophenotype to an immune-infiltrated TME with diverse cell-cell immune interaction patterns and spatially regulated exhaustion of CD8+T-cells.

### Myelonets - interconnected functional networks of myeloid subpopulations

Four of the five immune-rich RCNs were especially enriched in myeloid cells (RCN14 to 16, RCN18). We identified individual networks of myeloid cells belonging to these neighborhoods using Delaunay clustering, which identifies cells with direct cell-cell contacts. This analysis revealed distinct interconnected networks of myeloid cells, defined as networks with > 9 cells as “myelonets” **(Figure 3f)**. The size of these networks ranged from small groups of ten cells to large myelonets with thousands of cells. The mean number of cells per individual network was similar in chemo-naive vs IDS samples (**S.Fig 3f-i**), suggesting that the sizes of these networks are not affected by chemotherapy exposure. However, the size varied significantly according to the cell types present in myelonets; the smallest myelonets were identified inside RCN16 (mean 22 cells, SD 20) and RCN18 (mean 78 cells, SD 221) neighborhoods containing also stromal and immune cells and enriched in CD11c+ myeloid cells (RCN16). By contrast, the larger myelonets were predominated by IBA1+CD11c+ macrophages in RCN14 (mean 101 cells, SD 412), and IBA1+CD163+ macrophages in RCN15 (mean 146, SD 379) (**S. Fig 3j**). These findings suggest that the smaller myelonets are more frequently in contact with both tumor, stromal and other immune cells, and the larger myelonets are rather uniform clusters dominated by myeloid-myeloid interactions.

Notably, hierarchical clustering of the functional states of the myeloid cells in the myelonets showed divergent expression profiles within the myelonets for inflammation, antigen presentation, glutamine uptake, proliferation and interferon activation (**Figure 3g**). The myelonets exhibited functional rather than patient-specific clusters, and hierarchical clustering analysis revealed that myelonets in RCN14 neighborhoods, abundant in IBA1+CD11c+ macrophages, demonstrated the most active phenotype indicated by increased expression levels of Annexin A2, MHC II, pSTAT1, SNAT1, and Ki67.

The proportion of T-cells in these myeloid-rich RCNs was higher than in RCNs primarily composed of tumor or stromal cells, except for RCN14 enriched in IBA1+CD11c+ macrophages (**S.Fig 3a**). There were also CD8+T and CD4+T cells residing inside the myelonets, with the highest T-cell proportions in in RCN17 i.e. the T-cell neighborhood, followed by the smaller myelonets in the the tumor-stroma containing RCN16 and RCN18.

The functional marker expressions in the macrophages (MHC II, Ki67, Annexin A2, SNAT1, pSTAT1) and CD4+T and CD8+T cells (TIM3, pSTAT1, CD45RO, SNAT1, Ki67) inside the myelonets did not correlate with the size of myelonets (data not shown), indicating that the cellular states are regulated independently of the myelonet size, and that that also the small myelonets are functionally active.

Interestingly however chemotherapy highly induced the expression of the terminal exhaustion marker TIM3 in CD8+ T-cells residing in the RCN14 and RCN16 neighborhoods enriched specifically in CD11c and CD163 expressing myeloid cells, while the expression decreased in RCN17 enriched in T-cells, RCN15 rich in IBA1+CD163+macrophages as well as the tumor, and stromal-rich RCNs (**Figure 3h**). Moreover, chemotherapy induced the expression of Ki67 and SNAT1 in CD8+T-cells residing in myeloid cell rich RCN14, RCN15, RCN16 and RCN18, suggestive of TCR activation as a result of myeloid-T-cell spatial crosstalk. However, simultaneously pSTAT1 expression was decreased, suggesting compromised effector functionality^35^. Importantly, chemotherapy-induced exhausted phenotypes of tumor-recognizing CD8+T-cells^36^ were distinctively embedded in spatial myeloid neighborhoods with chronic antigen cross presentation. Our results suggest spatially restricted activation and exhaustion patterns in the TME induced by NACT, with exhausted TIM3+CD8+ T cells residing mainly in the tumor islets before NACT, but the balance shifting towards myelonets after NACT (**Figure 3h)**. Similar trends were seen also for CD4+ T-cells and FOXP3+CD4+ Tregs residing in different RCNs (**S. Fig 3k-l**). Interestingly, the expression of CD45RO, a marker of memory function^37^, was lower in the other immune rich neighborhoods after NACT, except in RCN14 enriched in IBA1+CD11c+ macrophages (**Figure 3h**) This finding suggests the presence of spatial immunostimulatory signaling potentiating CD8+T cell plasticity and persistence that takes place distinctively in the networks rich in M1-like macrophages after NACT (**Figure 3h**). A small increase in the CD45RO expression in CD8+T-cells was also seen in two stromal cell rich neighborhoods, RCN11 and RCN12 rich in myofibroblasts. However in general the CD8+T-cells in stromal neighborhoods seemed to harbor a quiescent state (low Ki67, pSTAT1, MHCI, SNAT1, CD45RO, TIM3) both before and after NACT when compared to other RCNs. Jointly, our results underpin the inherently spatial extrinsic cues shaping CD8+T-cell phenotypes in the TME of HGSC.

### Myeloid and T-cell interactions portray chemotherapy responses at the tumor-stromal interface

Chemotherapy exposure associated with the cell type compositions of three RCNs: RCN7 corresponding to the tumor-stromal interface, RCN14 characterized by the presence of IBA1+CD11c+ macrophages, and RCN8 characterized by the presence of Desmin+stromal cells (**Figure 4b, S. Fig 4a-b**). We next focused on the detailed characterization of the RCN7, as it was the most heterogeneous and dramatically chemo-shaped RCN: the cell type proportions in this RCN clustered into two groups which were significantly associated with chemotherapy exposure (Fisher’s exact test, p=0.00021). Moreover, the cell-cell interaction enrichment and depletion patterns differed within the RCN7 between IDS and chemo-naive samples **(Figure 4a,c, S.Fig 4c-d)**. Permutation tests revealed chemotherapy-induced enhanced cell-cell interaction patterns between CD8+T-cells neighboring IBA1+CD163+ macrophages, IBA1+CD11c+ macrophages, CD163+ macrophages, CD11c+ myeloid cells and CD4+ T cells (**Figure 4c**). Moreover, the spatial interactions between CD8+ T cells, CD4+T-cells and FOXP3+CD4+ T cells with neighboring CD31+SMA+ stromal cells, possibly reflecting overlaying pericytes with endothelial cells, were enhanced after chemotherapy. On the contrary, the chemo-naive samples displayed higher spatial interaction patterns between CD8+ T cells and the tumor metaclusters as compared to IDS samples (**Figure 4c**). This finding highlights the immunomodulatory cell-cell interaction patterns of CD8+ T cells occurring at the tumor-stromal neighborhoods induced by chemotherapy. This phenomenon was also validated in an independent HGSC cohort analyzed with t-CycIF for markers for tumor cells, macrophages and CD8+ T cells, with a trend for higher proximity density score for CD8+ T cells and macrophages at the tumor-stromal neighborhoods in IDS samples **(S.Fig 4f)**. Figure **4d** shows examples of enhanced CD8+ T cell macrophage spatial interactions for IDS samples, specifically at the tumor-stromal interface. Importantly, when combining both t-CycIF datasets, we observed a significant association of chemotherapy-induced CD8+T-cell-macrophage interaction in patients with a longer response to chemotherapy (**S. Fig 4g**). This association of enhanced spatial proximity and chemotherapy response was specific to the tumor-stroma interface and did not reach statistical significance when calculated from CD8+ T cell and macrophages irrespective of the neighborhood of origin (**S. Fig 4h**). This finding potentially reflects the tissue remodeling by macrophages and subsequent interactions with recruited CD8+ T cells at sites with increased tumor regression. Consistently, spatial transcriptomics analysis of a publicly available independent cohort of 8 HGSC IDS samples^38^ showed that the deconvolution scores for macrophages in the tumor-stromal interface and stromal areas were linked to good response to chemotherapy **(S. Fig.4i)**. Taken together, our results provide insights to the chemotherapy-associated inflammatory responses seen in HGSC samples39 and point towards increased immunomodulatory interactions of CD8+ T cells and macrophages occurring in response to platinum-based chemotherapy.

**Figure 4.**
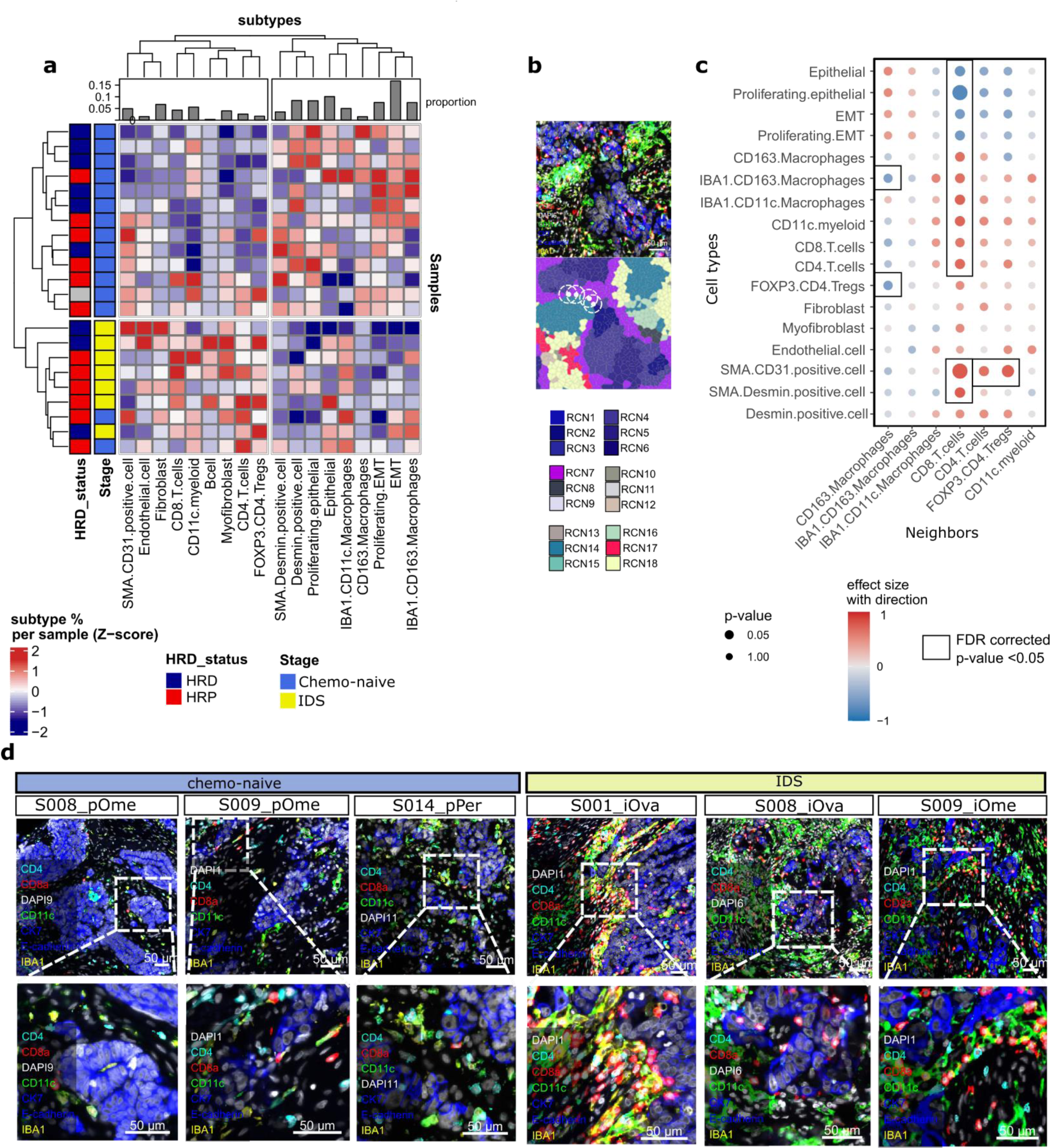
The tumor-stromal interface shows divergent CD8+T-cell and myeloid interaction patterns after chemotherapy. **a** A representative image of a tumor area and neighboring stroma with the corresponding RCN neighborhoods shown below the immunofluorescent image. The neighboring cells for subsequent interaction analyses were performed with a 45 micron radius as depicted by the white dashed circles. **b** Heatmap showing row-wise z-scores for cell type proportions belonging to RCN7 per sample clustered by hierarchical clustering. Annotations on the left include HRD status (red=HRP, blue=HRD) and treatment stage (blue=chemo-naive, yellow=IDS), the bar plot on the top shows the number of individual cell types belonging to the RCN7. **c** Dot plot showing changes in cell-cell interaction in chemo-naive and IDS samples in the RCN7. The y-axis represents the cell types and the x-axis their immune cell neighbors. The color of the dots represents the Wilcoxon test effect size, with values over 0 representing increased interaction probability in IDS samples, and values less than 1 in chemo-naive samples, respectively. The size of the dots represents the Wilcoxon p-value of the comparison. Those dots highlighted by black squares have an FDR<0.05 in the comparison between chemo-naive and IDS samples. **d** Representative images of the tumor-stromal interface and associated immune cell infiltrate in three chemo-naive and three IDS samples.

### Underlying transcriptional programs for peritumoral interacting macrophages and CD8+T-cells

To further study the gene expression programs in the spatial interactions of tumor cells, macrophages and CD8+ T cells, and specifically at the tumor-stromal interface, we performed t-CycIF guided spatial transcriptomics (GeoMx) in the independent Oncocys-OVA cohort of 16 whole-slide chemo-naive and IDS HGSC samples (**Figure 5a**). All patients except for one were genomically annotated as HRP **(Figure 1b)**. To dissect the roles of spatial CD8+T-cell–macrophage interactions, we selected four types of regions of interest (ROIs) using t-CycIF data on an adjacent tumor section, specifically considering the presence or absence of IBA1+ macrophages and CD8+ T cells adjacent to the tumor boundary **(Figure 5b, Supplementary Table 2)**. By using our developed pipeline^40^ to integrate t-CycIF images with GeoMx (**see Materials-Methods**), we segmented the tumor and stromal regions and collected signals from the ROIs separately from these compartments (areas of interest: AOIs), with each AOI consisting of approximately 100 cells. In total, we captured the whole transcriptome signals from 316 AOIs with the average number of 7450 genes detected per AOI. As expected, dimensionality reduction methods revealed patient-specific clustering for tumor, but not for the peritumoral stromal compartments (**S. Fig. 5a**).

**Figure 5.**
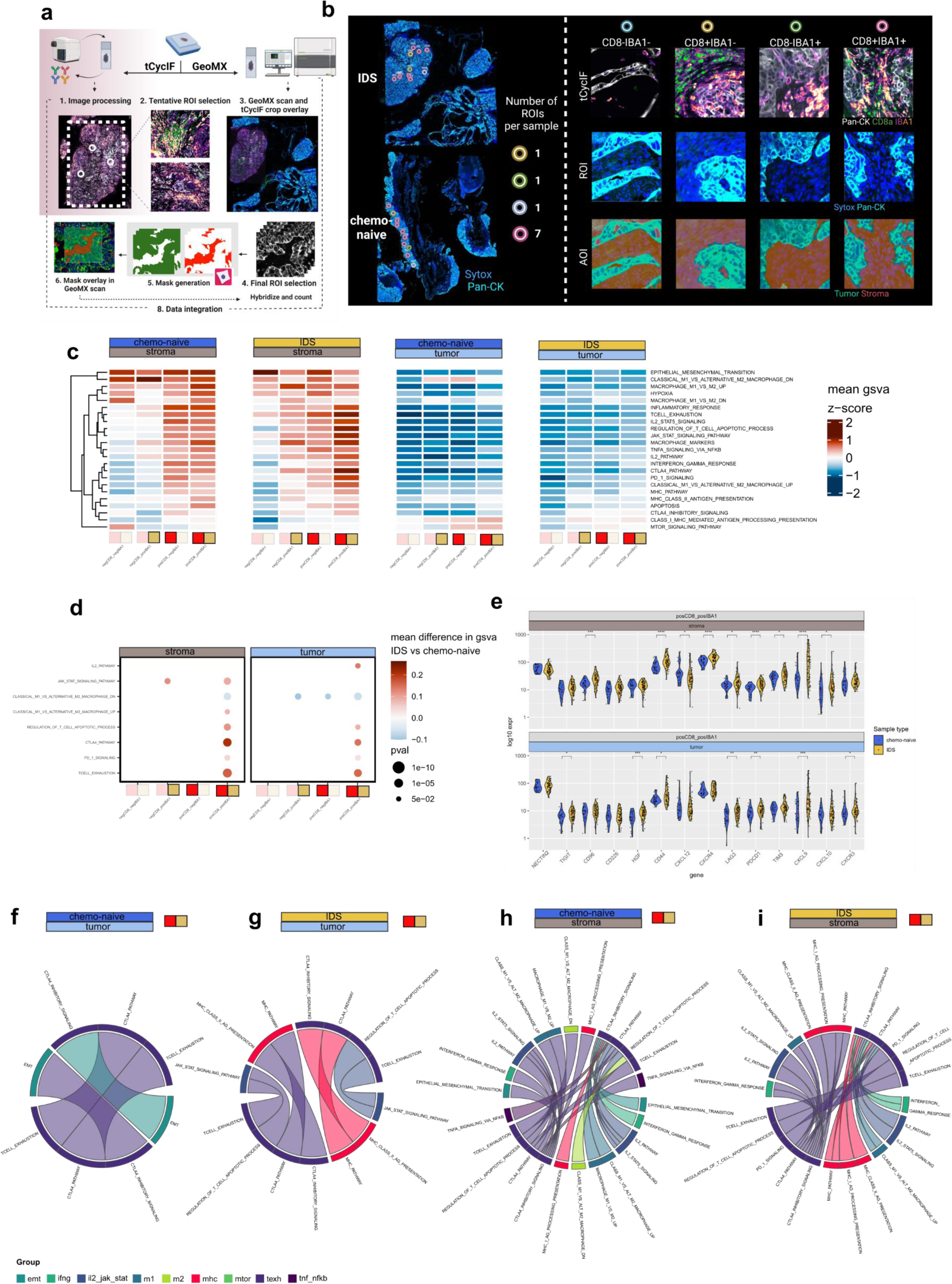
GeoMx spatial transcriptomics reveals differences in pathway activities between regions of different CD8+ T-cells and IBA1+ myeloid cells abundance. **a** Workflow **b** Left panel: A representative image of tissue slide stained for GeoMx spatial transcriptomics sequencing, with selected ROIs based on the relative abundance of CD8+T-cells and IBA1+ macrophages within a slide. Right panel: Representative images of each one of 4 ROI types. Top row displays a corresponding fragment of t-Cycif image of the adjacent tissue slide, used for guided ROI selection. The slide is stained with antibodies against Pan-CK, CD8 and IBA1. Middle row represents selected ROI on GeoMx tissue slide, stained with Sytox for cell nuclei and Pan-CK. Bottom row represents masks created over ROI, representing tumor and stromal areas (AOIs). **c** Clustered heatmaps representing mean z-score values of GSVA scores for 23 pathways. Columns represent each of 4 ROI types, rows are clustered based on values similarities between pathways. **b** represents stromal and tumor AOIs of chemo-naive and IDS samples **c** represents stromal and tumor AOIs of IDS samples from patients with long and short PFS. **d** Dot plots representing statistically significant differences (two-sided Wilcoxon rank-sum test) in GSVA scores between IDS vs chemo-naive samples for each of 4 ROI types and AOI compartments separately. Color represents the difference in mean GSVA score, while size of the dot represents the p-value of Wlcoxon test. Dot plots show the pathways, from Fig. 5 b,c which showed significant differences. **e** Violin plots representing differences in expression of selected genes within CD8+IBA1+ ROI type show differences between chemo-naive and IDS sample. Statistical significance was assessed using a two-sided Wilcoxon rank-sum test. Expression values are represented in log10 scale, with red dot denoting the mean value. **f-i** Circle plots showing significantly correlated (p-value <= 0.05, spearman correlation coefficient > 0.6) groups of pathways in CD8+IBA1+ AOIs from stromal and tumor compartments in chemo-naive and IDS samples separately. Each thin line represents one significantly correlated pathway from a specific pathway group **(Supplementary table 4)**. Thick lines represent multiple pathways. Only pathways correlated with the T-cell exhaustion pathway group (general T-cell exhaustion, CTLA4 and PD1 signaling pathways) are shown. Each of the connections is shown twice - once in the color of the pathway group of interest (various colors) and once in the color of T-cell exhaustion group (light green), to highlight the bilateral nature of interactions.

We calculated gene set variation analysis (GSVA) scores for 23 selected pathways connected to immune response (**Supplementary Table 3**), to validate our previous findings linked to the macrophage–CD8+ T cell crosstalk, separately in each AOIs (**see Materials-Methods**). Overall, we observed higher activity of most of the pathways in stromal AOIs in comparison to tumor, reflecting higher overall immune activity in this spatial peritumoral compartment in the tumor tissue **(Figure 5c).** This observation is in line with our findings on T- cell infiltration occurring in the stromal rather than in the tumor neighborhoods **(Figure 3c)**.

Detailed pathway analyses using gene set variation analysis (GSVA), revealed that chemotherapy significantly induced the activity of several pathways connected with T-cell exhaustion (signatures for canonical T-cell exhaustion, CTLA-4, PDL-1 signaling and T-cell apoptosis regulation). Interestingly, these pathways were significantly enriched exclusively in AOIs with colocalizing CD8+ T-cells and IBA1+ macrophages, both in the tumor and stromal compartments (**Figure 5d**). Notably, we did not observe significant induction of the T-cell exhaustion signature^41^ in CD8+T-cells during chemotherapy within the non-spatial scRNA-seq data, in either paired- or grouped analyses **(S. Fig 5b-c**), although single exhaustion markers were induced by chemotherapy in deconvoluted bulkRNA-seq data **(Figure 2e**). These findings support the notion that chemotherapy-induced T-cell exhaustion patterns are inherently spatially restricted, and mediated by direct cell-cell interactions with macrophages **(Figure 5d**, **Figure 3h).** Additionally, IL-2 activity was induced by chemotherapy in the tumor compartment exclusively in AOIs with CD8+T-cells and IBA1+ macrophage interactions **(Figure 5d).** Since IL-2 is known to promote CD8 T-cells activation as well as T-reg maturation^42^, this observation supports the hypothesis of chemotherapy-induced excessive T-cell activation by macrophages. Chemotherapy also induced pathways that suggest a shift from M2 towards M1 macrophages in the stromal compartment only in AOIs with CD8+T-cells and IBA1+ macrophages. For the tumor compartment however, the same shift towards M1 macrophages was observed for both double-positive CD8+T-cell—IBA1+macrophage AOIs, as well as single-positive AOIs containing either CD8+T-cells or IBA1+macrophages. That suggests a prominent role of CD8+T-cells in chemotherapy-induced M1 polarization in intratumoral stroma, but less so within tumor areas **(Figure 5d)**. Furthermore, JAK-STAT activity was elevated after chemotherapy in stromal areas with IBA1+ macrophages regardless of CD8+T-cell presence, indicating a role for JAK-STAT pathway in the chemo-induced macrophage states **(Figure 5d).** Next, we observed that chemotherapy induced the expression of key T-cell exhaustion related genes and ligand-receptor pairs, previously found to be connected with CD8+T-cell-macrophage interactions, exclusively within CD8+IBA1+ AOIs **(Figure 5e, S. Fig. 5f)**. Particularly, CD96, CD44, HGF, CXCL12, CXCR4, LAG3, CXCL9, CXCL10, PDCD1 and TIM3 were induced by chemotherapy in the stromal compartment, while HGF, CD44, CXCL9, CXCR3, PDCD1, LAG3 and TIGIT were induced in the tumor compartment (**Figure 5e).**

Additionally, we inferred 14 cancer-related pathway activities using PROGENy scores^43^ (**see Materials-Methods**). We found that the expression of WNT and TGFβ pathways, which are involved in maintenance of immune homeostasis^44,45^, were significantly decreased after chemotherapy at the stromal areas with colocalizing CD8+T-cells and macrophages adjacent to the tumor border (**S. Fig 5d-e**). TGFβ secreted by cancer-associated fibroblasts (CAFs) has been found to restrict CD8+T-cell infiltration into tumor, dampen the inflammatory functions of macrophages and maintain immune homeostasis as well as restrict responses to ICB^44^. Moreover, Wnt signaling promotes T-cell exclusion and resistance to ICB^46,47^, while also inducing the production of CD8+ memory stem cells^48^. scRNA-seq data showed TGFβ pathway activity primarily in the fibroblasts and endothelial cells, while the WNT pathway was also active in the majority of immune cells **(S.Fig 5g)**. Taken together, these pathways might promote the T-cell exclusion in chemo-naive samples.

Finally, to elucidate the molecular foundations of CD8+ T cell-IBA1+ macrophage crosstalk, we computed correlation coefficient scores between all GSVA scores of selected pathways and analyzed the results jointly for pathways with shared underlying biological processes (e.g., T cell exhaustion, CTLA-4 inhibitory signaling, PD1 signaling, regulation of T-cell apoptotic process, IL2-JAK-STAT axis, EMT) (**Supplementary Table 4)**. Predominantly positive correlations were observed, and the analysis of significant correlations (p-value <= 0.05, Spearman correlation coefficient > 0.6) revealed distinct biological processes contributing to T-cell exhaustion across distinct clinical samples and spatial TME compartments (**Figure 5f-j, S. Fig 5h-k**). In general, the tumor compartment exhibited fewer significant correlations with the T-cell exhaustion signature, indicating an active role for the non-epithelial compartment in regulating anti-tumor immunity. Chemo-naive tumor areas displayed correlations only with the EMT signature, while chemo-exposed tumor areas exhibited correlations with IL2-JAK-STAT and MHC II pathways **(Figure 5f-g).** Additionally, in the stromal compartment at baseline, T-cell exhaustion was associated with TNF⍺-NFκβ as well as M1 macrophage pathway activities (**Figure 5h**). Following chemotherapy, IFN-ɣ pathways displayed enhanced association with T-cell exhaustion, indicating chemo-induced inflammatory signaling crosstalk **(Figure 5i)**. Connections were also identified between T-cell exhaustion and IL2-JAK-STAT and M1 macrophages, although they were not affected by chemotherapy treatment. Notably, within the CD8+ IBA1+ spatial neighborhoods in both stromal and tumor compartments, chemotherapy induced enhanced correlations between MHC class II signaling and T-cell exhaustion (**Figure 5f-i**). This finding suggests that the chemotherapy-induced T-cell exhaustion processes are likely driven by prolonged antigen presentation on MHC molecules by myeloid cells. Additionally, our results reveal links to other critical immune stimulatory pathways such as IL2-JAK-STAT induced by chemotherapy, potentially acting through the induction of the MHC presentation machinery^49^. Our findings shed light on chemotherapy-induced spatiotemporal changes in the interactions between CD8+ T-cells and macrophages, emphasizing distinct molecular signaling circuits affecting T cell phenotypes localized in the peritumoral compartment.

### Combining anti-TIGIT antibody with chemotherapy enhances CD8+ T-cell activation in HGSC

To functionally validate the role of chemotherapy on the identified immunosuppressive signaling pathways, we used our established functional immuno-oncology platform consisting of immune-competent patient-derived cultures50 (iPDCs) from three chemo-naïve and one chemotherapy-exposed HGSC patients (**Figure 6a-b**). Using flow cytometry, we tested the effect of 3-day treatments in the different cultures and confirmed that the main effector immune cell types were present in the iPCDs throughout the duration of the experiment. (**Figure 6c**). Then, we assessed the impact of single- and combination ICB treatments with pembrolizumab (anti-PD1 antibody) and tiragolumab (anti-TIGIT antibody). Notably, we observed that the single agent and combination ICBs resulted in a general increase in granzyme B expression - a marker of cytolytic activity51 - in CD8+ T-cells in the iPDCs in comparison to the corresponding non-treated control (**Figure 6d, Supplementary Table 5**). Moreover, in the chemo-naïve samples, combining chemotherapy to ICB also resulted in a marked increase of granzyme B positivity of CD8+T-cells in comparison to tiragolumab-based immunotherapy alone (**Figure 6e**). Deviations from this trend in two conditions are likely attributable to patient-specific responses to the selected doses (see **Materials-Methods**). These observations imply a dynamic interplay between chemotherapy and immunotherapy, where the cytotoxic activity of chemotherapy exposed CD8+ T cells may be unleashed by anti-TIGIT and anti-PD1 immunotherapies. Further investigation into the underlying mechanisms governing these interactions and especially into the precise “window” period for introducing immunotherapy is warranted to optimize treatment strategies for HGSC.

**Figure 6:**
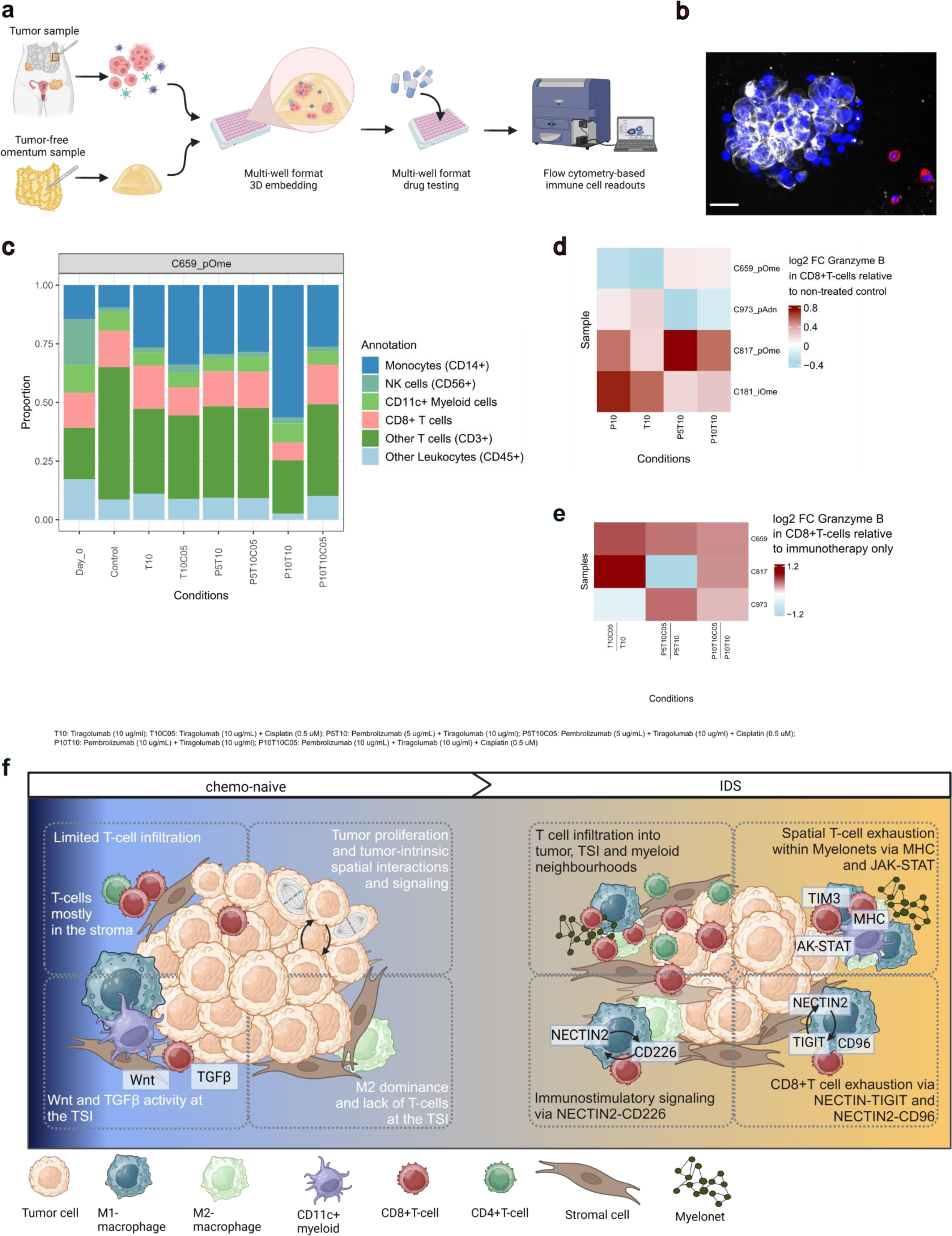
Immunocompetent spheroids show T-cell activation following treatment with anti-TIGIT, anti-PD1 and cisplatin. **a** General workflow illustrating the culturing process of patient-derived immuno-competent cultures (iPDCs) and subsequent flow cytometry analysis. **b** Representative immunofluorescence image of the cancer cell spheroids alongside infiltrating immune cells within the iPDCs. Blue: Hoechst, Red: CD45, Gray: CK7, Yellow: Death cell marker. Scale bar = 50 um **c** Barplots presenting the proportions of the various immune cell types within one iPDC. **d** Heatmap visualizing the Log2 Fold Change (FC) of Granzyme B expression in CD8+ T-cells of each tiragolumab-based immune checkpoint blockade (ICB) treatment condition compared to non-treated control. **e** Heatmap visualizing the Log2 Fold Change (FC) of Granzyme B expression in CD8+ T-cells of each tiragolumab-based ICB treatment condition combined with cisplatin compared to the corresponding ICB treatment condition alone. **e** Summary figure of findings regarding the differences of chemo-naive and IDS samples.

## Discussion

Our multiomics study, incorporating data from two large patient cohorts at an unprecedented single-cell and spatial scale, illuminates the dynamic shifts within the TME, encompassing alterations in cell subtypes, states, spatial arrangements, and the intricate molecular and cellular interconnections activated in response to platinum-based chemotherapy **(Figure 6f)**. In the chemo-naive HGSC TME, tumor-intrinsic interactions predominate, accompanied by spatially restricted T-cell infiltration and an immune-suppressive tumor-stroma interface. However, chemotherapy augments the versatility of macrophage- and T cell states, while concurrently reducing that of cancer cells fostering a more intricate interaction potential within the immune compartment of HGSCs. The chemotherapy-driven orchestration of T cell infiltration and recruitment is governed by strategic ligand-receptor interactions via *CD226* and *NECTIN*2, as well as signaling through the adhesive CXCR4 and CD44 receptors. Simultaneously, active immune modulation by macrophages occurs within the myelonets and at the tumor-stromal interface through the specific, competing TIGIT- and CD96-NECTIN2 mediated molecular and cellular crosstalk. These findings illuminate a promising avenue for therapies targeting TIGIT and T-cell exhaustion, holding the potential to unleash CD8+ T-cell-mediated immune surveillance from the myeloid niches. We thus demonstrate that the impact of chemotherapy extends beyond its conventional role, revealing a nuanced orchestration of spatially confined T-cell exhaustion through direct cell-cell interactions with macrophages. These revelations not only unveil potential opportunities for immunotherapeutic interventions but also provide a compelling rationale for patient stratification.

In accordance with our findings, prior studies have documented an escalation in both stromal and intraepithelial CD8+ T-cell infiltration following NACT^25,52–54^. While the surge in CD8+ T-cells theoretically aligns with an anticipated response to PD1/PDL1 checkpoint blockade^55^, such correlation has not been consistently observed in earlier studies, potentially attributed to concurrent immunosuppressive factors^52^. Furthermore, diverse outcomes have been reported in terms of overall and immunosuppressive macrophage populations^54,52,53,19^. The use of varied technologies, and the also herein observed macrophage heterogeneity and plasticity likely contribute to these disparate findings. From our t-CycIF data, we discerned no statistically significant variations in macrophage infiltration patterns following NACT. Nevertheless, our characterization of the functional states of myelonets - networks of interconnected myeloid cells - revealed spatially enhanced infiltration and restricted exhaustion of CD8+ T-cells and within these neighborhoods in response to chemotherapy. Myelonets, ranging from small networks comprising ten cells to those spanning thousands of cells, thus unveil previously undiscovered yet biologically and spatially pivotal roles in regulating TME function and controlling anti-tumor immunity.

Moreover, a recurrent, cellularly versatile neighborhood comprising tumor, stromal and immune cells, and localized to the tumor-stromal interface, exhibited distinct cell type compositions, as well as patterns of cell-cell interaction enrichment and depletion influenced by chemotherapy. Chemotherapy induced enhanced interactions between CD8+ T cells and myeloid cells, potentially linked with a more favorable chemotherapy response. Notably, the increased spatial proximity of CD8+ T cells and macrophages at the tumor-stromal interface following chemotherapy suggests sensitivity to ICB^12,22^, with the potential to lead to improved long-term outcomes in patients responding to platinum-based chemotherapy. The close proximity of myeloid cells to CD8+ T cells and the existing T-cell immunity in these patients appears to be crucial for eliciting an effective immunotherapeutic response. Across the data modalities studied, we saw enhanced CD8+ T cell to myeloid cell interactions in samples collected at IDS as compared to chemo-naive samples, and further in those samples with a better chemotherapy response, shedding light to the fibroinflammatory reaction occurring in response to NACT.

Our results from spatially resolved transcriptomics underpin the chemotherapy-induced enhanced signaling crosstalk between T cell exhaustion and antigen presentation in areas with colocalizing CD8+ T cells and macrophages. Antigen presentation and long-duration synapses by macrophages have been shown to prime T cell exhaustion resulting in immunosuppression^56^ as opposed to antigen presentation by dendritic cells which has been shown to sustain antitumor immune responses with the production of terminally differentiated effector population^57^. In chemo-exposed samples, the highly inflamed tumor-immune microenvironment and tumor-cell derived neoantigens harvested and presented by macrophages possibly cause T-cell hypofunction. This is concordant with a previous report from breast cancer, where macrophages harbored more inflamed phenotypes in environments with exhausted T-cells^58^.

Chemotherapy distinctly induced enhanced interactions of *TIGIT* and *CD96* in CD8+ T cells with *NECTIN2* in macrophages. This together with the chemotherapy-induced expression of *TIGIT* highlights the potential of targeting TIGIT in conjunction with chemotherapy in ovarian cancer. TIGIT has been found to be highly expressed in exhausted CD8+ T cells in HGSC, and targeting TIGIT reduced tumor growth in HGSC mouse models^59^. Further, the TIGIT-NECTIN2 axis has been found to be important in different stages of ovarian cancer^60^. Importantly, the dual blockage of TIGIT and the PD1-PDL1 axis is required for efficient antitumor responses^61^. Currently, no anti-TIGIT therapies are approved for clinical use^62^. Phase III clinical trials targeting TIGIT along with the programmed death pathway have been active in non-small cell lung cancer^63^ and melanoma^64^. Recently, the TIGIT-NECTIN2 axis has emerged as a crucial immune checkpoint also in neuroblastoma^65^ and hepatocellular carcinoma^66^. Importantly, our results highlight the possible utility of targeting TIGIT and PD1 in conjunction with chemotherapy treatment in patients with HGSC to unleash T-cells from the myeloid niches. We observed an increased CD8+ T-cell activation when the iPDCs were treated with tiragolumab-based immunotherapy and when the cells were simultaneously (PDS samples), or previously (IDS sample), exposed to chemotherapy. While our iPDC system did not allow sequential treatment administration to directly assess T-cell exhaustion followed by immunotherapy rescue, our results suggest that the CD8T+-cell activity can be unleashed via ICB in HGSC.

We acknowledge that our study with detailed analysis of several multiomic datasets is not without limitations. Regarding the identification of cell phenotypes, the inherent properties of the t-CycIF marker panel left a subset of cells (14%) unassigned to a clear cell subpopulation. Moreover, tissue sampling from the same anatomical site was not always feasible, especially after chemotherapy, as the abundance of tumor cells was markedly affected in response to treatment. Acknowledging the existence of potentially unique TMEs in different metastatic sites, recording of more anatomical sites during chemotherapy would strengthen the findings. However, most of our samples were collected patient- and site-matched from intra-abdominal metastases, and the main findings, i.e., spatial phenotypes were consistent across metastatic sites, and confirmed in an independent dataset.

In conclusion, our findings underscore the intricate co-evolution of the tumor-immune-stromal milieu during the course of chemotherapy, exerting profound effects on anti-tumor immunity revealing potential therapeutic targets. We demonstrate that interconnected networks of myeloid cells, or myelonets drive spatial exhaustion of CD8+ T cells, especially in patients responding to chemotherapy. As new combination treatment strategies involving immunotherapies are clinically explored, it becomes imperative to consider the dynamic shifts within the tumor-immune microenvironment during standard-of-care chemotherapy. This awareness is crucial for optimizing strategies that effectively unleash CD8+ T-cell antitumor immunity, liberating it from myeloid cell niches and facilitating robust tumor cell eradication in patients with ovarian cancer.

## Materials

### Cohort description

Tumor samples and clinical data were collected in two independent prospective clinical trials; DECIDER NCT04846933, and Oncosys-OVA NCT06117384. Samples for WGS, bulk RNA-seq, scRNA-seq and t-CycIF were collected from the DECIDER study of patients with HGSC treated at the Turku University Hospital, Finland. Tumor specimens (scRNA-seq: n=51, t-CycIF n=22, bulk RNA-seq n=60, WGS n=60) were collected from HGSC patients (n=44) at the time of laparoscopy or primary debulking surgery (chemo-naive) and interval debulking surgery (IDS) (**Supplementary Table 2**). In addition to newly produced scRNA-seq data from 25 tumors, the scRNA-seq cohort contains data from three clinical specimens originally published in ^33^, 21 specimens originally published in ^16^, and two specimens originally published in ^67^. Genomic HRD annotations were previously published in^24,68,69^, and bulk RNAseq in^70^. The DECIDER study was approved by the Ethics Committee of the Hospital District of Southwest Finland (ETMK 145/1801/2015). Tumor samples for t-CycIF, GeoMx, bulk RNA-seq and WES (n=16) were collected from patients (n=11) participating in the prospective ONCOCYS-OVA study in the Helsinki University Hospital, Finland (HUS334/2021). Additionally, spatial transcriptomics data for additional patients with HGSC (n=8) were publicly available (GSE211956)^38^.

In accordance with the ethical standards from the 1975 Declaration of Helsinki, every patient from DECIDER and ONCOSYS-Ova trials provided an informed written consent to the collection, storage, and analysis of the samples and subsequent data.

## Methods

### Sample preparation for scRNA-seq

Tumor specimens were incubated overnight in a mixture of collagenase and hyaluronidase (Department of Pathology, University of Turku) to obtain single-cell suspensions. Chromium Single-Cell 3′ Reagent Kit v.2.0 and v.3.0 (10x Genomics) were used to prepare scRNA-seq libraries. Libraries were sequenced on Illumina HiSeq 4000 (Jussi Taipale Lab, Karolinska Institute, Sweden), HiSeq 2500, and NovaSeq 6000 instruments (Sequencing Unit of the Institute for Molecular Medicine Finland, Finland).

### Preprocessing scRNA-seq data

The published data (**Supplementary table 6**) were jointly processed with the new data produced for this study as follows. The fastq files were processed with the Cell Ranger v.6.0.1 pipeline (10x Genomics) for demultiplexing, alignment, barcode processing, and unique molecular identifier (UMI) quantification. Gene annotation was done using GENCODE v25 and GRCh38.d1.vd1 was used as reference genome. The count matrices were loaded into Seurat (v3.2.2) and dimensionality reduction and initial clustering using k-means were performed using the UMAP algorithm. Based on marker expression, cells were assigned to three major cell types: epithelial tumor cells (*WFDC2*, *PAX8*, *MUC16*), stromal cells (*COL1A2*, *FGFR1*, *DCN*), and immune cells (*CD79A*, *FCER1G*, *PTPRC*). miQC^67^ was used separately per each cell type to filter out cells with high proportion of mitochondrial reads or low number of genes without assigning a hard threshold. The cells expressing a combination of *DCN*, *PAX8* or *PTPRC* were also filtered to exclude possible doublets. After initial filtering, dimensionality reduction, clustering and cell-type assignment were performed again. Moreover, to ensure high-quality cells for downstream analysis, stromal cells with a number of UMI (in log scale) <11, immune cells with < 10.5 and tumor cells with <12 were removed through further filtering.

### Cell type annotation in scRNA-seq data

The celltypist23 package (v.0.1.9) was used for automated annotation of cell types. The predicted cell type labels were acquired by running the function *celltypist.annotate*(model = ‘Immune_All_Low.pkl’, majority_voting = True).

### Differential cell abundance analysis in scRNA-seq data

We used the speckle R package^71^ to calculate the changes in cell proportions after NACT. After performing logit transformation of the cell proportions, we ran the *propeller* function to perform linear modeling to calculate the changes in cell proportions between IDS and chemo-naive samples in a pairwise fashion.

As the secondary approach, we used a robust strategy for differential abundance testing for scRNA-seq data^26,72^. The Milo framework models cell states as neighborhoods on a nearest neighbor graph and uses a negative binomial generalized linear model for hypothesis testing to identify differentially abundant neighborhoods between conditions of interest between (chemo-naive vs IDS samples).

We created a Milo object from the Seurat object, and built a k-nearest neighbors (KNN) graph with the function *buildGraph*(k = 200, d = 10), and created neighborhoods with the function *makeNhoods*(prop = 0.05, k = 200, d = 10, refined = TRUE). We counted the cells in the neighborhoods with the *countCells* function and calculated the distances between the neighborhoods with the *calcNhoodDistance* function. We created a design matrix using the metadata and ran the *testNhoods* function with the design (∼ 0 + class) to find the differentially abundant cell state neighborhoods between chemo-naive and IDS samples.

To further analyze gene expression in the Milo identified IDS enriched macrophage neighborhoods, we performed differential expression analysis (DEA) between pure macrophage neighborhoods that showed statistically significant enrichment in the IDS samples (cell_type == “Macrophages” & cell_type_fraction == 1 & logFC > 0 & SpatialFDR < 0.05) and those that did not show statistically significant changes (cell_type == “Macrophages” & cell_type_fraction == 1 & SpatialFDR >= 0.05). Seurat’s *FindMarkers()* function was used to identify differentially expressed genes above the specified threshold (logfc.threshold = 0.25) in at least 10% of the cells (min.pct = 0.10), using the Wilcoxon test (test.use = “wilcox”).

The clusterProfiler R package^73,74^ was used to perform overrepresentation analysis (ORA). The *enrichGO()* function was used to identify enriched gene ontology (GO) biological processes (BP) related to genes with higher expression in macrophages enriched in the IDS samples (pAdjustMethod = “BH”, pvalueCutoff = 0.01, qvalueCutoff = 0.05), and a dot plot was used to visualize the top 10 terms.

### Ligand-receptor interaction analysis

We used the MultiNicheNet framework^27^ to infer the changes in cell-cell interactions after NACT. We subsetted the Seurat object to retain cell types of interest (“Macrophages”, “Epithelial cells”, “Tem/Trm cytotoxic T cells”) in paired chemo-naive and IDS samples from matched patients. The *multi_nichenet_analysis* function was run with the following parameters: sample_id = “sample”, group_id = “class”, celltype_id = “cell_type”, covariates = “patient”, batches = NA, min_cells = 10, contrasts_oi = c(“’interval-primary’,’primary-interval’”), logFC_threshold = 0.75, p_val_threshold = 0.05, fraction_cutoff = 0.10, p_val_adj = TRUE, empirical_p_value = FALSE, top_n_target = 250. We used the *get_top_n_lr_pairs* function to filter the top 50 ligand-receptor interactions across all contrasts and used the *make_circos_group_comparison* function to create the circos plots to visualize the predicted interactions.

For ligand-receptor interactions specific to macrophages and CD8+ T cells, we selected the top 50 predicted interactions, ranked them based on scaled ligand activity in receiver cells (i.e., CD8+ T cells), and used the *make_sample_lr_prod_activity_plots* function to visualize the ligand-receptor pseudobulk product expression across samples and ligand activity in the same plot.

### Pathway activity inference in scRNA-seq data

We used the decoupleR R package^75^ to infer the activities of 14 signaling pathways in cell types present in the chemo-naive and IDS samples. The *get_progeny* function was used to access the gene weights for the top 500 genes stored in the PROGENy R package^43^ and the *run_ulm* function was used for univariate linear modeling on the normalized count matrix extracted from the Seurat object. The ComplexHeatmap R package^76,77^ was used to visualize the scaled pathway activity scores across cell types. The same approach was used to compare pathway activities in macrophages identified in the **Differential expression analysis (DEA) and overrepresentation analysis (ORA)** section.

### Quantification of T-cell exhaustion in scRNA-seq data

To quantify exhaustion in CD8+ T cells, we used the UCell algorithm^78^ by running the AddModuleScore_UCell function with default parameters to calculate the module score for a gene set linked to T-cell exhaustion^41^. The base::mean() function was used to calculate the average module scores for each sample. Paired samples Wilcoxon test (ggpubr::stat_compare_means(method = “wilcox.test”, paired = TRUE) was used to compare average T-cell exhaustion scores between paired chemo-naive and IDS samples. The complete list of genes is shown in **Supplementary Table 7**.

### Visium spatial transcriptomics analysis

Visium spatial transcriptomics analyses of the abundance of cell types in the proximity of tumor were made using publicly available dataset of 8 IDS HGSC samples with different response for chemotherapy (GSE211956)^38^. Based on H&E images, each spot was manually annotated by a pathologist as ‘tumor’, ‘mixed’ (partial tumor and stromal morphology), ‘stroma’ and ‘other’. For gene expression data, first the QC and preprocessing of input feature-barcode matrices in h5 format were made using the Seurat R package^79^. Spots with less than 500 features detected, more than 25% of mitochondrial genes or more than 20% of hemoglobin genes, were removed from the analysis. Next, feature-barcode matrices were normalized using the SCTransform function and merged with histological annotations. For stromal spots, 3 zones of tumor-stromal interface (TSI) were defined based on proximity to the closest tumor/mixed spot. To assess the abundance of certain cell types within the different spot types (tumor, mixed, tsi1, tsi2, tsi3, distant stroma), we used FindTransferAnchors function from Seurat R package. This deconvolution method produced for a given spot, a normalized score resembling the closeness to each cell type present in the labeled reference scRNAseq dataset (a downsampled scRNAseq dataset described in the previous section).

### t-CycIF imaging and image processing

The samples were stained consecutively with the validated antibodies **(Supplementary Table 8**) followed by scanning with the RareCyte CyteFinder scanner with the t-CycIF protocol^80^. The image files were corrected, stitched, and registered using the BaSiC tool and ASHLAR algorithm^81^. The raw ome-tiff images were segmented into cells based on probability maps created by UNET^82^ with a modified version of the S3Segmenter algorithm (https://github.com/HMS-IDAC/S3segmenter). Cytoplasm segmentation masks were created by expanding the nuclei region with a 2-pixel ring dilation. Each cell’s mean fluorescence intensity was computed by Python’s scikit-image library using in-house scripts. In total over the 22 whole slide images 8,912,899 cells were identified, from which first 1,448,531 cells were removed by cycifsuite’s detect_lost_cells function (https://github.com/yunguan-wang/cycif_analysis_suite/tree/MCF10A/cycifsuite) due to tissue movement or detachment. Further quality control was carried out in the Napari image viewer, and artifacts (hairs, folds, bubbles) were removed by manual area selection, affecting 43,470 cells.

### t-CycIF cell type calling

Log transformation was applied to the quantified single-cell signals. TRIBUS^83^ recently developed in our lab was first used to assign cells semi-automatically and iteratively into cell types using SOM-based clustering in R with single-cell feature quantification table and cell phenotype marker positivity and negativity as inputs. The markers used for cell type characterization are shown in **Supplementary Table 9.** First, tumor cells were separated from non-tumor cells, after which non-tumor cells were categorized into stromal, myeloid, and lymphoid cells, and these further into subtypes. The tumor cells were also separated to metaclusters using the expressions of CK7, E-cadherin, Ki67 and vimentin, and stromal cells were subcategorized into metaclusters using the expressions of SMA, Vimentin, CD31, CK7 and Desmin. Cells that were not assigned to any of the mentioned categories were labeled as “Other” (mean 14% of the cells per sample), these mostly resided at the stromal areas. Quality control for the cell type calls was performed using the *pl.image_viewer* function in Scimap-toolkit in python by overlaying the cell type calls on top of the images. Cells that had been mislabeled, mostly B-cells as “other” cells, were reclassified using the *pl.gate_finder(), pl.addROI_image()* and *hl.classify()* functions separately for each image for selected areas.

### t-CycIF spatial analysis

Cell-cell interactions with CD8+ and CD4+T-cells and their relationships to functional marker expression by tumor cells and myeloid cells were computed using the Giotto package in R^84^. The *FindInteractionChangedFeats* function with Delaunay network was used separately for each image to find the fold changes for markers which changed based on the spatial interaction with another celltype. Delaunay networks for the analysis of recurrent cellular neighborhoods were calculated using the spatstat package^85^ in R, with the maximum length between edges 45 pixels.

Recurrent cellular neighborhoods were computed using the Scimap package^86^ in python, by first calculating the fractions of neighboring cell types for each centering cell using *tl.spatial_count* with a radius of 100 pixels, and the resulting neighborhood matrix was then clustered using *tl.spatial_cluster* using kmeans clustering with k=18. 100 pixel radius was chosen to find large scale cellular structures from the single-cell spatial data, the radius encompasses around 3-4 cells in length. The identity of each cluster was defined by the proportions of cell types composing it, and by overlaying these on top of the images using the *pl.image_viewer* function in scimap. The Rao’s quadratic entropy per RCN was computed using the *rao.diversity* function in the SYNCSA R-package, with cell type proportions and functional marker expressions as inputs. Furthermore, the cell-cell interaction and avoidance patterns compared to a random background were computed using the *tl.spatial_interaction* function from the scimap-package with a radius of 45 pixels.

Centrality scores were calculated using the Squidpy^87^ package in python separately for each image for each cell type with the *gr.centrality_scores* function.

### Low-plex t-CycIF staining and image processing

The samples were stained with seven antibodies (**Supplementary Table 10**) following the t-CycIF protocol^80^. Images were obtained within the Pathology Department at the University of Helsinki, employing a high-throughput scanner equipped with an ECLIPSE Ni-E body, an ORCA-Flash4.0 LT PLUS camera by Hamamatsu, and a Märzhäuser Slide Express slide loader. Tile stitching and registration across different cycles were performed similarly as for highplex t-CycIF images. Nuclei segmentation was carried out by STARDIST^88,89^. Single-cell information was extracted similarly as for highplex t-CycIF images for 8,829,233 cells.

### Low-plex t-CycIF cell type calling

Cell type calling for low-plex t-CycIF was performed using the *pl.gate_finder*, *pp.rescale* and *hl.classify* functions for PanCK, IBA1 and CD8 for each image in scimap-toolkit in Python. The cell type calls were confirmed by overlaying them on top of the images using the *pl.image_viewer* function in scimap. Recurrent cellular neighborhoods were computed using the *spatial_count* function with r=100px and *spatial_cluster* function, with k-means clustering using k=10 for the combined dataset with the high-plex t-CycIF samples included and cell type classes (tumor, IBA1+macrophages, CD8+T-cells, other cells) aligned.

### GeoMx

The GeoMx spatial transcriptomics procedure was conducted at the Single Cell Analytics unit within the Institute for Molecular Medicine Finland (FIMM), adhering to the prescribed GeoMx guidelines for slide staining and scanning. The human Solid Tumor TME morphology marker kit was employed for this purpose. The tissue slide chosen to perform GeoMx sequencing was adjacent to the one on which lowplex t-CycIF have been performed, to enable guided ROI selection.

### GeoMx ROI selection with lowplex t-CycIF crop overlay

Lowplex t-CycIF stainings were employed to guide the selection of regions of interest (ROIs, n=160) in the GeoMx experiment. The chosen regions contained tumor-stroma interface features, indicated by the presence of PanCK+ and Vimentin+ cells, and a variety of CD8 (CD8+ T-cells) and IBA1 (macrophages) staining markers for optimal ROI identification within the t-CycIF crop (CD8+IBA1+, CD8+IBA1-, CD8-IBA1+, CD8-IBA1-) **(Supplementary Table 2)**. The ROIs were selected based on the relative abundance of CD8+ and IBA1+ cells observed within a given slide. The chosen t-CycIF crop was imported into the GeoMx software and manually superimposed onto the GeoMx scan using landmarks present in both images. Consistency in choosing similar types of areas, avoiding tissue folds, necrotic regions, or adipose tissue, while maintaining similarities in ROI size, tumor-stroma ratio and cell count across all samples was aimed. An external segmentation pipeline generated in CellProfiler^90^ was employed to establish distinct areas of Illumination (AOIs, n=320) separately for the tumor and stromal compartments.

### GeoMx and t-CycIF data pairing

Whole slide image alignment was achieved in QuPath^91^ through the Warpy plugin, which performs affine registration, incorporating the moving image to the coordinate system of the base image. The resulting files were exported as pyramidal ome.tiff files to be analyzed in other platforms.

GeoMx data quality control and preprocessing.

DCC files obtained from the sequence provider were handled with the usage of GeomxTools R package^92^, provided by Nanostring. Segment (AOI) and probe level QC were performed as described in the package vignette. Overall, four ROIs were removed due to insufficient quality, and remaining 316 were further processed. Only one probe was removed from the dataset due to insufficient quality. Later, counts for the probes targeting the same gene were aggregated. Additional QC was performed, based on LOQ (limit of quantification), as described in the GeomxTools vignette. Since we expected a high level of variance in our dataset, Gene Detection Rate parameter was set up to 0.01 (gene was removed if expression > LOQ in more than 1% of AOIs). Almost 2000 genes and none of the AOIs were removed during this procedure, leaving a total number of 16,727 genes detected in the whole dataset with sufficient quality.

Next, data was normalized using Q3 quartile normalization method from GeomxTools package and dimensionality reduction was performed using t-SNE and UMAP methods using Rtsne^93^ and umap^94^ R packages.

### GeoMx pathway activity analysis and correlation analysis

Based on the previous observations, we selected 23 pathways from the MSigDB database^95^, from Hallmark and Canonical Pathways collection, to compare their activities within different types of AOI and clinical features of the patients. Selected pathways were connected with the processes such as regulation of T-cells, macrophages and cancer cell biology, as well as immune response. T-cell exhaustion related gene set was retrieved from Zhang et al16. The full list of selected pathways can be found in the **Supplementary Table 3.** Q3 normalized gene expression matrix was used for pathway analysis. Selected pathway activities were inferred using the GSVA method with Poisson kernel from the GSVA^96^ R package. Additionally PROGENy scores, representing the core features of cancer biology, were calculated using the progeny^43^ R package with 10x permutation.

Statistical significance of the differences between pathways activities in different AOI types was assessed using a two-sided Wilcoxon rank-sum test.

We used Spearman correlation for analyzing the correlations between pathways in CD8+IBA1+ AOIs. We calculated correlation coefficients and p-values for all pathway combinations. Next, we grouped all pathways into groups resembling important biological signaling axis: T-cell exhaustion, IL2-JAK-STAT, TNF⍺-NFκβ, IFNγ, mTOR, MAPK, TGFβ-WNT, M1 and M2 macrophages, MHC presentation and EMT (**Supplementary Table 4)**. Next, we calculated a number of significant correlations (p-value <= 0.05, spearman correlation coefficient > 0.6). We analyzed only positive correlations, since no significant negative correlations were found within our data. At the end we compared the number of significant correlations between groups of pathways, separately for tumor and stromal compartments and chemo-naive and IDS samples.

### WGS and SBS3

WGS was performed as previously described8. Samples with a tumor purity <10% were excluded. COSMIC single base substitution reference signatures v3.3.1^97^ were used, adjusted for trinucleotide frequency as reported in^68^. HRD samples were defined as those with a positive SBS3 (Sig3) status and computed as described in^68^.

### Genomic Homologous recombination deficiency scoring

OvaHRDscars were calculated using the package ovaHRDscar (https://github.com/farkkilab/ovaHRDscar) as described in our previous publication using either WGS or SNP-array data^24^. ovaHRDscar package quantifies three types of allelic imbalances: loss of heterozygosity, telomeric allelic imbalances and large scale transitions of certain characteristics. For WGS data, calculation was based on copy-number profiles estimated with PURPLE^98^ using structural variants from GRIDSS2^99^. A cutoff value of ovaHRDscar >= 54 was used to define HRD positivity. Samples with a tumor purity <10% were excluded from calculations.

### Bulk RNA-seq data analysis

Fifty omental samples from 25 patients collected before NACT and at IDS were subjected to RNA sequencing. The preprocessing of bulk RNA sequencing reads was conducted using the SePIA pipeline^100^ within the comprehensive Anduril framework^101^ as previously described^33^. For the batch correction, the POIBM method was used with default parameters^102^. We employed the statistical framework PRISM^33^ to decompose the gene-level effective read counts into cancer, immune, and stromal-specific signals. To assess the proportion of tumor, stromal, and immune cell subtypes, we employed the Kassandra deconvolution algorithm^103^ using transcripts-per-million (TPM) expression values as input. To characterize the tumor proliferation rate, we acquired the expression of the genes related to the tumor proliferation rate functional signature, as was previously described^104^. To evaluate transcription factor activity, we used the DecoupleR framework v1.4.3^75^ in Python and accessed the CollecTRI gene regulatory network^105^ using decoupler.get_collectri(). We inferred TF enrichment scores using the univariate linear model approach with decoupler.run_ulm(), using gene-level statistics from DESeq2^106^ output (stat value) as input. For comparing the expression of selected genes, we utilized the trimmed mean of M-values (TMM) of the genes.

### Immuno-competent patient-derived cultures

The tissue for this experiment was obtained from three chemo-naive HGSC patients and from one chemo-exposed sample from the ONCOSYS-Ova prospective cohort. iPDCs were generated from the harvested tissue following the protocol described by Nagaraj *et al^50^*. Briefly, a fresh ∼1 mm3 omental tumor piece was finely minced and mildly digested with 1 U/mL Dispase II (Sigma-Aldrich, D4693) and 5 U/mL RQ1 RNAse-free DNase I (Promega, M6101) in AdF+++ medium (DMEM/F-12 (Gibco) supplemented with 1X Glutamax (Gibco, 35050038), 10 mM HEPES (Sigma-Aldrich, H0887), and 1% v/v Pen/Strep (Gibco, 15140122)). The resulting cell suspension was washed and resuspended in RBC lysis buffer (Roche, 11814389001). Subsequently, cells were washed again, and cell count and viability were assessed using a LUNATM *fl* Dual Fluorescence Cell Counter (Logos Biosystems). Then, cells were resuspended in complete medium (Advanced DMEM-F12,Gibco) supplemented with cytokines and growth factors as reported by Nagaraj *et al.50* and seeded in ultra-low attachment 48-well suspension culture plates (Greiner Bio-One, 677102), embedded in an omentum gel-based matrix at a cell density of 1x103 cells/µL. On the next day, iPDCs were treated with cisplatin (Selleck Chemicals, S1166) pembrolizumab (Selleck Chemicals, A2005), and tiragolumab (Selleck Chemicals, A2028) as single agents and in different combinations. The doses for cisplatin and pembrolizumab were selected based on previous work of th-e effective doses in the same model system^50^, and tiragolumab concentrations based on the literature^107^. After three days, cells were harvested and CD8+ T cell activation was estimated by measuring the expression levels of granzyme B with multiparameter flow cytometry. Samples were run in a BD LSRFortessaTM cell analyzer (BD Biosciences), and the details about the employed antibodies are listed in **Supplementary Table 11**. Resulting flow cytometry data was analyzed using FlowJO, cell proportions and heatmaps were plotted using R scripts.

### Statistical analyses

Statistical analyses were performed using R version 4.0.3. The paired sample tests were performed using the two-tailed Wilcoxon signed-rank test, and unpaired tests using two-tailed Wilcoxon rank-sum test using the *wilcox.test* from stats package in R. The paired changes in cell type abundance in scRNA-seq data were assessed using linear modeling with *propeller* function from the speckle R-package^71^. P-values were corrected using the Benjamini-Hochberg procedure when appropriate (in Figures 2a, 3c, S.Fig 3c-d, 4b). Spearman correlation coefficients and p-values were used when calculating correlations. The threshold for significance was set up as less or equal to 0.05 in all analyses.

## Acknowledgements

This work was supported by the European Union’s Horizon 2020 research and innovation programme under grant agreement no. 667403 for HERCULES and no. 965193 for DECIDER. This study was co-funded by the European Union (ERC, SPACE 101076096). Views and opinions expressed are however those of the author(s) only and do not necessarily reflect those of the European Union or the European Research Council. Neither the European Union nor the granting authority can be held responsible for them.

This study was additionally funded by the Sigrid Jusélius Foundation, Research council of Finland (grant numbers 1339805, 350396), Cancer Foundation Finland, Finnish Medical foundation, Ida Montin foundation, Emil Aaltonen foundation, Biomedicum foundation, K. Albin Johanssons foundation, Instrumentarium Foundation, and HORIZON-MSCA-2021-PF project 101067835 (M.F.). R00CA256497 (A.J.N.) from NCI. The authors are grateful to the CSC-IT Center for Science (Finland) for computational resources, and to the FIMM-HCA and Genomics units for their services.

Figures 1a, 5a, 6a and 6e were created with Biorender.com.

## Data availability

The raw images and quantified signals for t-CycIF data can be found from Synapse XXX upon publication. Unpublished raw scRNA-seq data will be deposited in the European Genome-phenome Archive (EGA), and processed scRNA-seq data will be deposited in Gene Expression Omnibus (GEO) upon publication.

## Code availability

The scripts used for scRNA-seq, t-CycIF and GeoMx analyses can be found from www.github.com/XXXXX upon publication.

## Contributions

I-M.L processed and analyzed the t-CycIF data and wrote the manuscript. E.P.E generated, processed, and analyzed the scRNA-seq data and wrote the manuscript. I.N generated and analyzed the GeoMx data and wrote the manuscript. M.H and A.N performed the iPDC experiments, analyzed the data and wrote the manuscript. A.S preprocessed the t-CycIF data. D.A analyzed the RNAseq data and wrote the manuscript. A.J performed the GeoMx experiments, Z.L. assisted with GeoMX data analyses. M.F preprocessed the scRNA-seq data. F.P and J.O provided the HRD calculations. M.S, U-M.H, E.K, A.V and J.H provided samples and clinical data. T.V and A.N.J supported the data analysis. S.H and P.S supported the data analysis and provided resources. A.V and A.F conceived and supervised the study and wrote the manuscript.

---

**Supplementary Figure 1.**
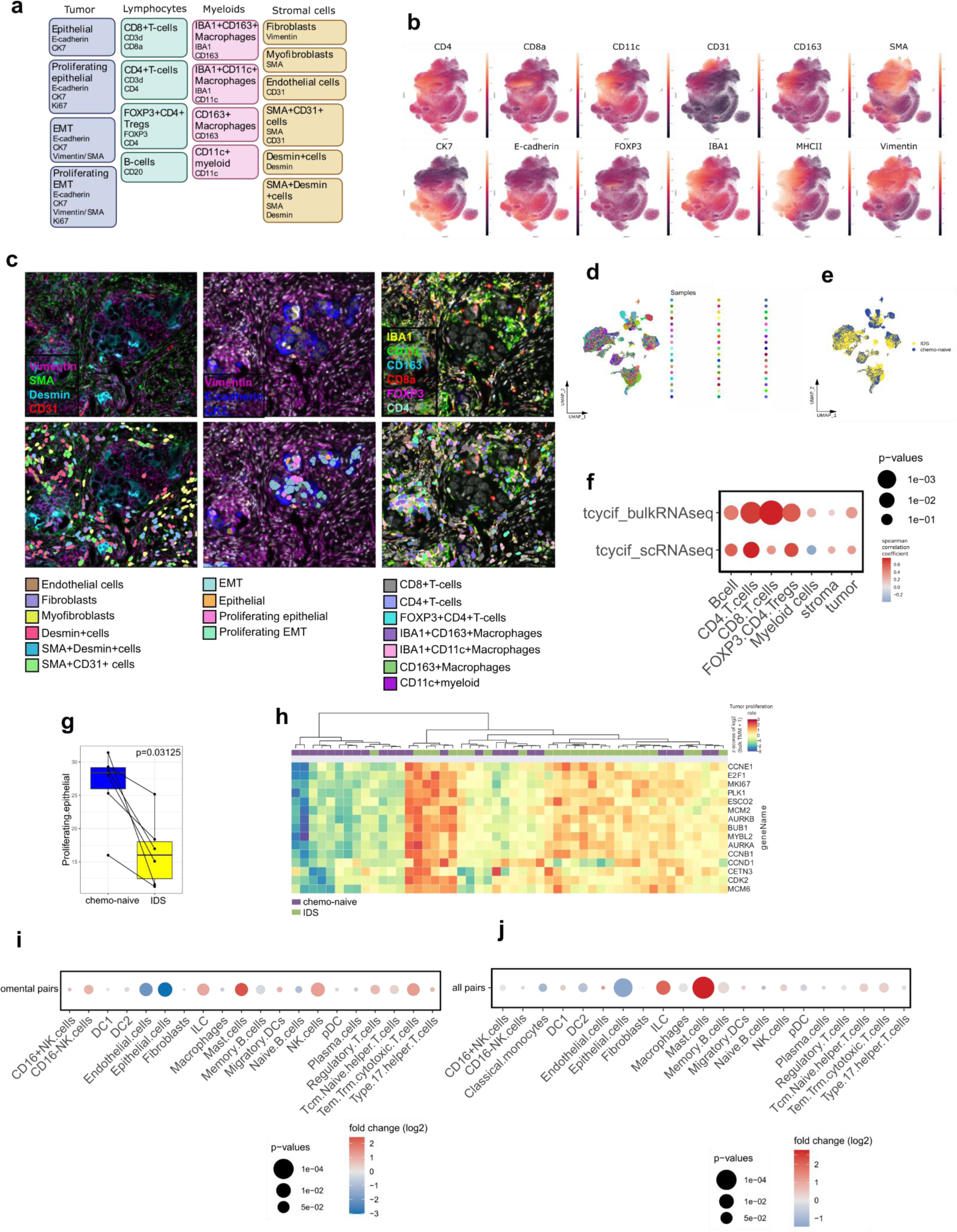
Single-cell phenotyping of HGSC samples. **a** Markers defining the cell types in t-CycIF single-cell data **b** UMAPs colored by t-CycIF marker expressions of cell type markers for the cell types shown in Figure 1c. **c** Representative examples of cell type calls with panels displaying different markers on the same tissue area. d-e UMAPs of scRNA-seq cells colored by sample type and sample of origin, respectively. **f** Dotplot showing the sample-wise spearman correlations between cell type proportions in t-CycIF and scRNA-seq data as well as in t-CycIF and bulk RNAseq data. **g** Change in proliferating epithelial metacluster proportions out of all tumor cells in the t-CycIF data in chemo-naive and IDS samples. **h** Heatmap with hierarchical clustering of genes for tumor proliferation in RNAseq data. The annotations include sample type and HRD status. *i-j* Dot plots showing the log2 fold change and p-values for fine grained cell types in paired omental samples, and all paired samples, respectively. In boxplots, the black horizontal lines represent the sample medians, the boxes extend from the first to the third quartile, and the whiskers indicate the values at 1.5 times the interquartile range. Individual dots represent values per sample.

**Supplementary figure 2:**
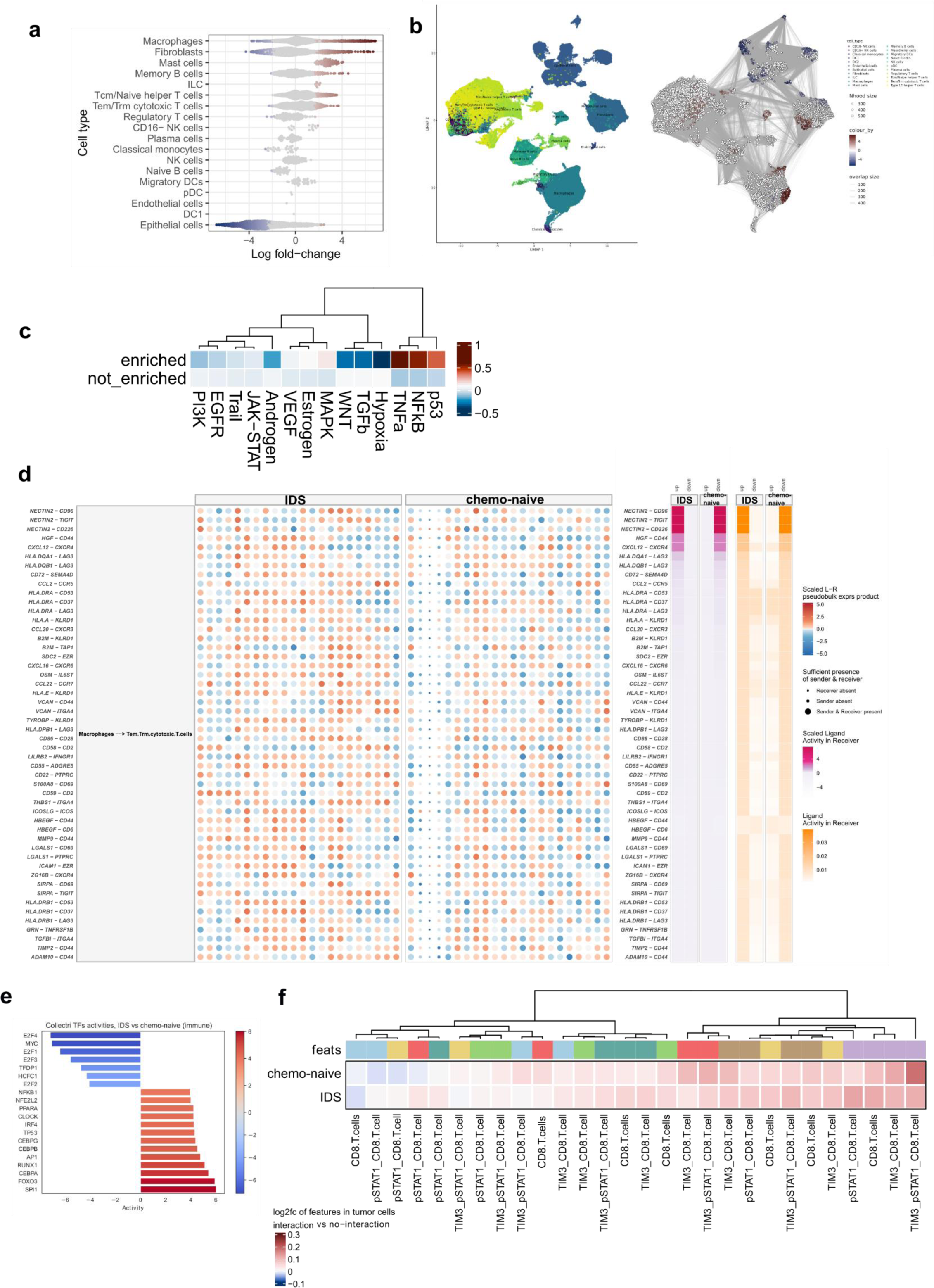
Tumor phenotype changes from chemo-naive to IDS samples. **a** Beeswarm plot showing the enrichment of cell state neighborhoods with all cell type categories and **b** Milo neighborhoods projected onto the UMAP projection. **c** Heatmap of the scaled macrophage pathway PROGENY scores enriched after chemotherapy. **d** MultiNichenet plot corresponding to Figure 2d, showing the scaled ligand-receptor pseudobulk expression product per sample. **e** Heatmap with log2 fold change expression values (color of the heatmap) of functional markers in tumor cells (features in top annotation) interacting with CD8+T-cells as compared to without interacting CD8+T-cells in chemo-naive and IDS samples. **f** Top 20 significantly different transcription factor activities in immune deconvoluted RNAseq data between IDS and chemo-naive samples. Positive values indicate higher activity in IDS samples.

**Supplementary Figure 3.**
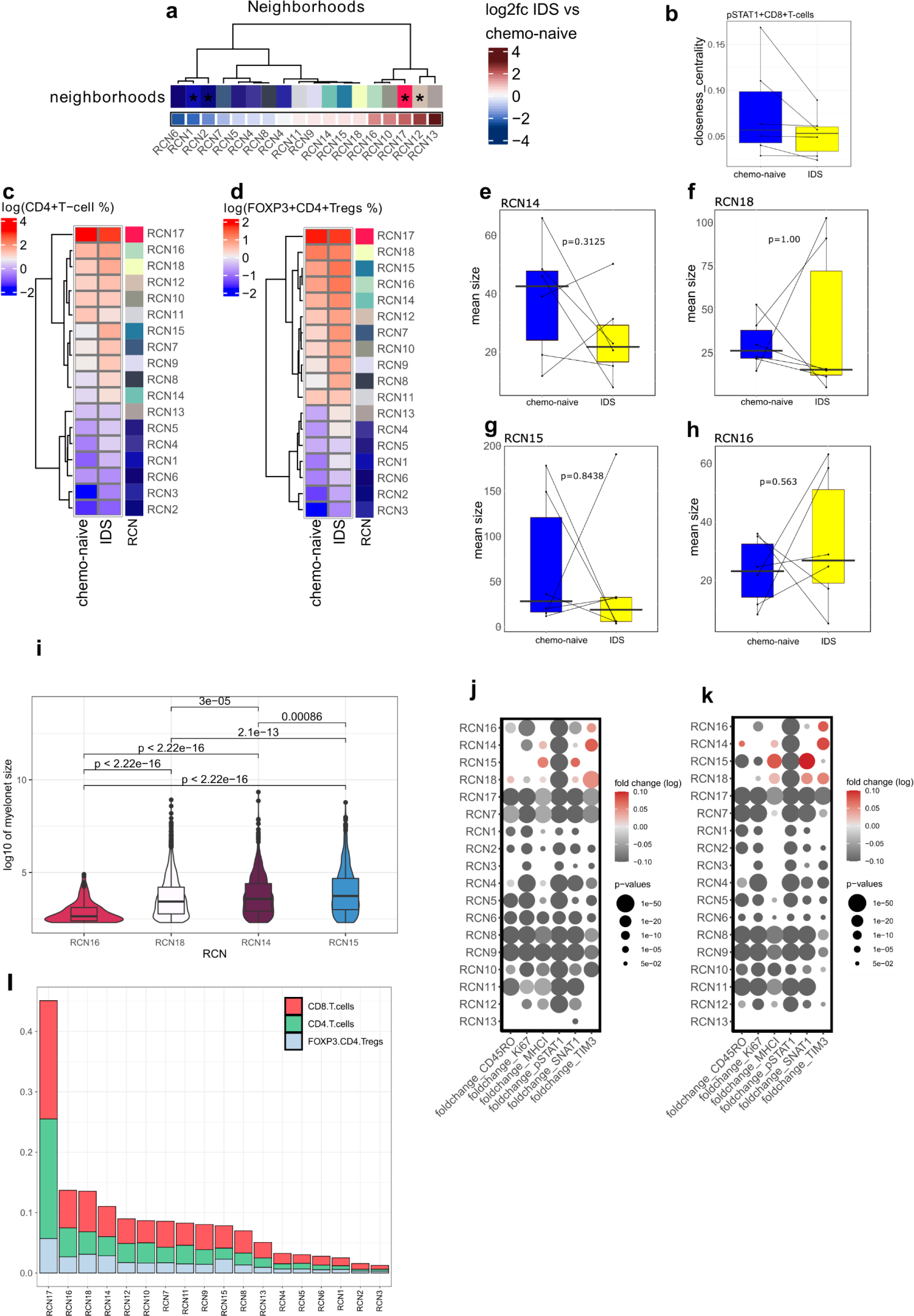
Characterization of recurrent cellular neighborhoods. **a** Barplot showing the proportions of CD8+T-cells, CD4+T-cells and FOXP3+CD4+Tregs in each RCN. **b** Heatmap of the change in RCN proportions between chemo-naive and IDS samples. The color represents the log2 fold-change and those RCNs marked with an asterisk have a p-value <0.05 in paired Wilcoxon test. **c** The change in closeness centrality score of pSTAT1+CD8+ T cells from chemo-naive to IDS samples. **d** log2 CD4+ T cell and **e** FOXP3+CD4+Treg proportions per RCN in chemo-naive and IDS samples. **f-i** Box plots with paired Wilcoxon p-values for the change RCN size in RCN14, RCN18, RCN15, and RCN16 between chemo-naive and IDS samples **j** Violin plots with box plots showing cell numbers per individual myelonet in RCN15, RCN18, RCN14 and RCN15 **k** Dot plot for the log2 fold-change in mean functional marker expression between IDS and chemo-naive samples for CD4+T-cells and **l** for FOXP3+CD4+ T-cells. The size of the dots represents the FDR-corrected Wilcoxon p-values and the color of the dot the fold-changes. In boxplots, the black horizontal lines represent the sample medians, the boxes extend from the first to the third quartile, and the whiskers indicate the values at 1.5 times the interquartile range. Individual dots represent values per tumor.

**Supplementary Figure 4.**
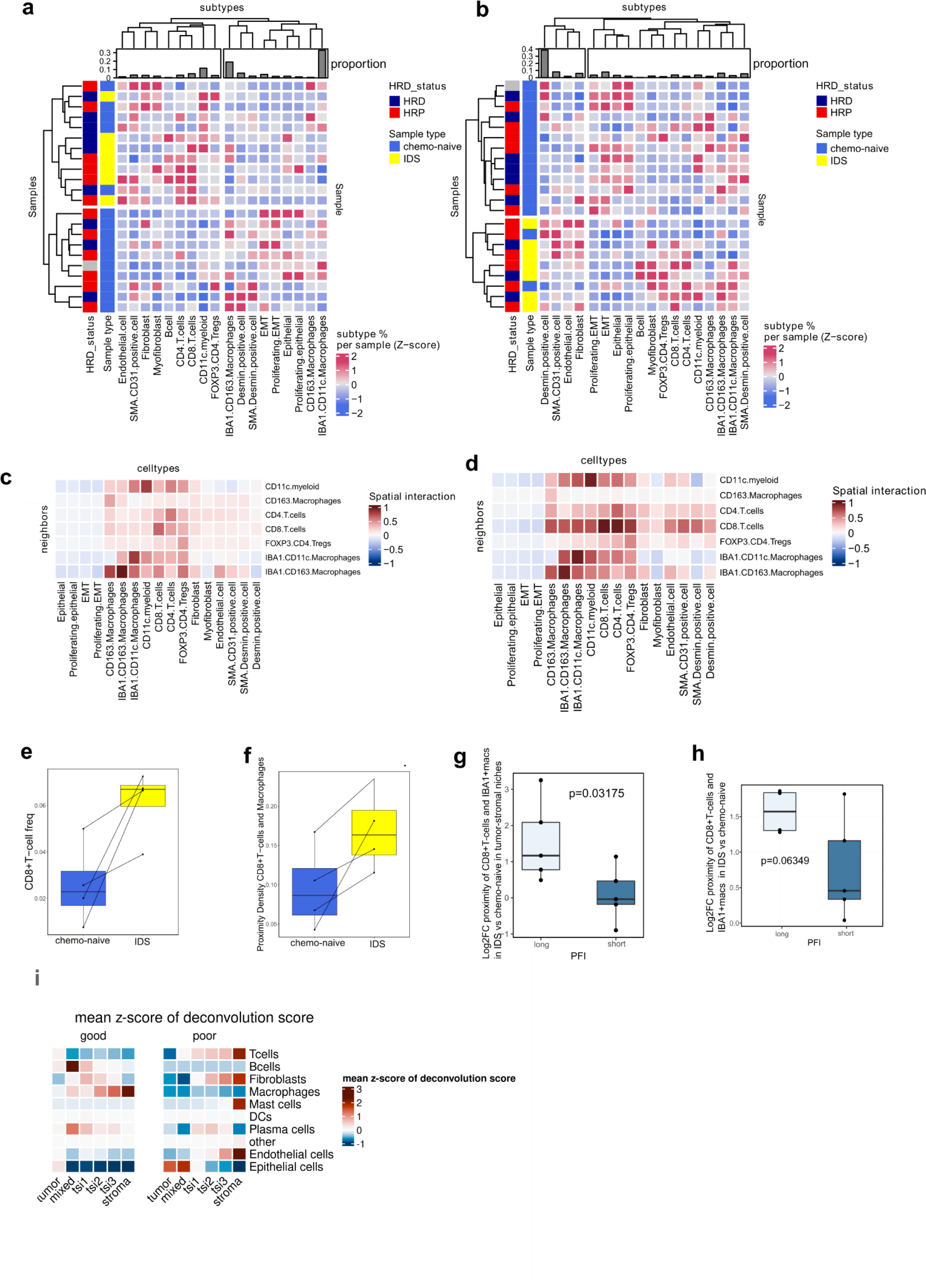
Characterization of RCN7. **a** Heatmap with hierarchical clustering of the z-scored cell type proportions in RCN14 and **b** in RCN8. The annotations include sample type (blue=chemo-naive, yellow=IDS), HRD-status (red=HRP and blue=HRD) and the bar plots indicate the count of the specific cell type in that RCN. **c** and **d** Heatmaps for the values for cell-cell interactions in RCN7 corresponding to the dotplot in Figure 4c are shown separately for chemo-naive and IDS samples, respectively. **e** Box plot showing a trend towards increased CD8+T-cell proportion out of all cells in IDS as compared to chemo-naive samples in the lowplex-cycif dataset. **f** Increased proximity density of CD8+T-cells and IBA1+macrophages at the RCN corresponding to tumor-stromal interface in the lowplex t-CycIF dataset. **g** Box plot showing the log2 fold change in the proximity density of CD8+T-cells and IBA1+ macrophages in IDS as compared to chemo-naive samples separately for patients with a long or short PFI (cutoff 12 months) in the RCN corresponding to tumor-stromal interface. **h** Box plot showing the log2 fold change in the proximity density of CD8+T-cells and IBA1+ macrophages in IDS as compared to chemo-naive samples separately for patients with a long or short PFI (cutoff 12 months) in the entire tissue areas. The black horizontal lines represent the sample medians, the boxes extend from the first to the third quartile, and the whiskers indicate the values at 1.5 times the interquartile range. Individual dots represent values per tumor. **i** Heatmap showing the mean z-score of cell type deconvolution score in different regions from Visium IDS samples with good (n=3) and poor (n=5) chemotherapy response.

**Supplementary Figure 5.**
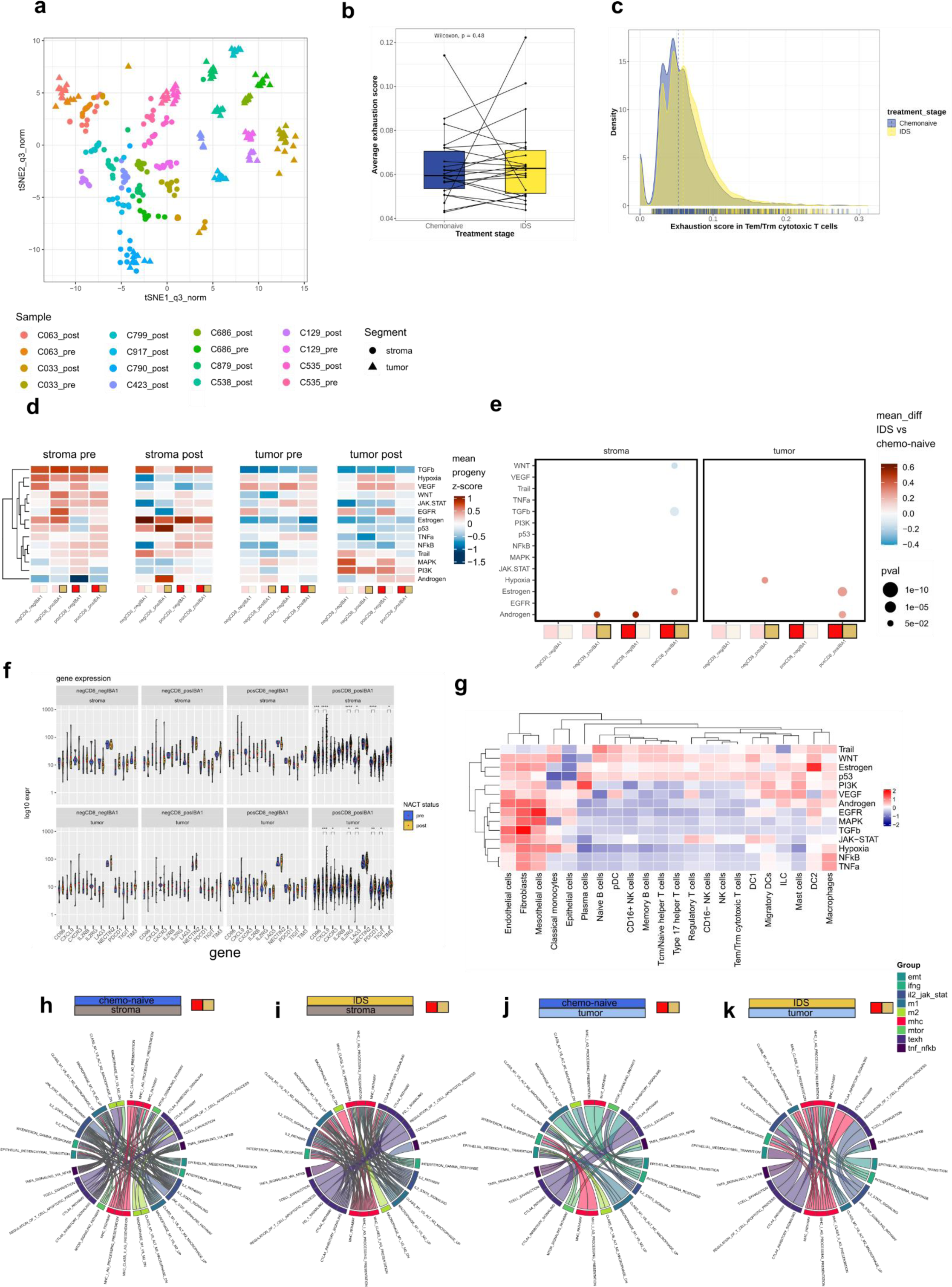
GeoMx spatial transcriptomics dimensionality reduction, PROGENy scores and expression of selected genes. **a** Scatterplot showing t-SNE dimensionality reduction for the 316 AOIs. Colors represent samples, while shape the tumor or stromal type of the AOI. **b-c** Box plot and density plot show no and minimal change in T-cell exhaustion score from the scRNA-seq in CD8+T-cells between chemo-naive and IDS samples. **d** Clustered heatmap representing mean z-score values of PROGENy scores for 14 PROGENy pathways representing different features of cancer biology. Columns represent each of 4 ROI types (CD8-IBA1-, CD8+IBA1-, CD8-IBA1+ and CD8+IBA1+), and rows are clustered based on values similarities between PROGENy pathways. The heatmap represents stromal and tumor AOIs of chemo-naive and IDS samples.**e** Dot plots representing statistically significant differences (two-sided Wilcoxon rank-sum test) in PROGENy scores between IDS vs chemo-naive samples. Color of the dot represents the difference in mean progeny score, while size of the dot represents the p-value of the Wilcoxon test for each of 4 ROI types (CD8-IBA1-, CD8+IBA1-, CD8-IBA1+ and CD8+IBA1+) and AOI compartments separately. **f** Violin plots representing differences in expression of selected genes within all 4 ROI types (CD8-IBA1-, CD8+IBA1-, CD8-IBA1+ and CD8+IBA1+) show differences between chemo-naive and IDS samples. Statistical significance was assessed using a two-sided Wilcoxon rank-sum test. Expression values are represented in log10 scale, with red dot denoting the mean value. **g**. PROGENy scores for cell types from chemo-naive scRNA-seq data. **h-k** Circle plots showing significantly correlated (p-value <= 0.05, spearman correlation coefficient > 0.6) groups of pathways in CD8+IBA1+ AOIs from stromal and tumor compartments in chemo-naive and IDS samples separately. Each thin line represents one significantly correlated pathway from a specific pathway group **(Supplementary table 4)**. Each of the connections is shown twice, in the colors representing both correlated pathway groups to highlight the bilateral nature of interactions.

