## Supplementary tables for "Chemotherapy induces myeloid-driven spatial T-cell exhaustion in ovarian cancer"

Supplementary table 1: Differential expression analysis for macrophages Milo

|  | p_val | avg_log2FC | pct.1 | pct.2 | p_val_adj |
| --- | --- | --- | --- | --- | --- |
| GCLM | 0 | 0.566735436710083 | 0.567 | 0.23 | 0 |
| CSF1 | 0 | 0.567700076177834 | 0.341 | 0.063 | 0 |
| S100A11 | 0 | 0.62712804011826 | 1 | 0.989 | 0 |
| RSAD2 | 0 | 2.07670332151983 | 0.669 | 0.255 | 0 |
| CYP1B1 | 0 | 1.21551067662969 | 0.872 | 0.445 | 0 |
| DPP4 | 0 | 0.38530523948425 | 0.275 | 0.039 | 0 |
| SERPINE2 | 0 | 0.595451370341463 | 0.25 | 0.045 | 0 |
| CCL20 | 0 | 1.09263287310927 | 0.79 | 0.39 | 0 |
| TM4SF1 | 0 | 1.40389585040716 | 0.537 | 0.19 | 0 |
| CXCL8 | 0 | 1.17990160295531 | 0.999 | 0.869 | 0 |
| CXCL1 | 0 | 0.996166453057606 | 0.775 | 0.322 | 0 |
| CXCL3 | 0 | 0.802875158923621 | 0.936 | 0.59 | 0 |
| CXCL2 | 0 | 1.09894637760906 | 0.909 | 0.659 | 0 |
| SLC7A11 | 0 | 0.828089632015532 | 0.744 | 0.301 | 0 |
| SQSTM1 | 0 | 0.640554012113668 | 0.912 | 0.688 | 0 |
| SNX10 | 0 | 1.03193451175548 | 0.828 | 0.596 | 0 |
| TXN | 0 | 1.28706883020231 | 0.979 | 0.922 | 0 |
| MALAT1 | 0 | -0.855692435562527 | 1 | 0.997 | 0 |
| MMP1 | 0 | 1.35406258621309 | 0.324 | 0.064 | 0 |
| MGST1 | 0 | 0.682014031006476 | 0.635 | 0.25 | 0 |
| TXNRD1 | 0 | 1.13011213718164 | 0.778 | 0.377 | 0 |
| C12orf49 | 0 | 0.640101257011789 | 0.545 | 0.226 | 0 |
| TDRD3 | 0 | 0.377827790330181 | 0.333 | 0.071 | 0 |
| C15orf48 | 0 | 0.700585343112518 | 0.978 | 0.749 | 0 |
| AQP9 | 0 | 0.582923402567407 | 0.676 | 0.319 | 0 |
| TPSB2 | 0 | 0.666957023021003 | 0.294 | 0.065 | 0 |
| TPSAB1 | 0 | 1.07187946689079 | 0.338 | 0.074 | 0 |
| HS3ST2 | 0 | 0.521434679476138 | 0.345 | 0.07 | 0 |
| CCL18 | 0 | 1.27709063473559 | 0.512 | 0.189 | 0 |
| SERPINB2 | 0 | 1.09766160664841 | 0.818 | 0.308 | 0 |
| FTLP3 | 0 | 0.736165900581501 | 0.598 | 0.229 | 0 |
| MMP9 | 0 | 1.22762335978466 | 0.738 | 0.451 | 0 |
| MT-RNR1 | 0 | -1.43750117405049 | 0.867 | 0.976 | 0 |
| MT-RNR2 | 0 | -1.07441186385444 | 0.971 | 0.995 | 0 |
| SDCBP | 1.93057289850555e-306 | 0.560468364868034 | 0.942 | 0.776 | 7.68812045371865e-302 |
| C4orf32 | 1.23728077176784e-301 | 0.361611148226898 | 0.296 | 0.075 | 4.92722321741106e-297 |
| ATP6V0D2 | 4.0386753639611e-298 | 0.256163227228629 | 0.203 | 0.034 | 1.60832169019023e-293 |
| LRP12 | 6.16504008269675e-291 | 0.450880753602899 | 0.481 | 0.189 | 2.45510391213233e-286 |
| GM2A | 5.06174203076841e-290 | 0.740087785395331 | 0.673 | 0.409 | 2.0157375289129e-285 |
| SLC41A2 | 9.56996770270391e-287 | 0.357167647231462 | 0.357 | 0.111 | 3.81104823824778e-282 |
| NAA50 | 5.54041555183237e-286 | 0.518274990419435 | 0.618 | 0.309 | 2.20635968520621e-281 |
| RACK1 | 5.58456761468711e-286 | -0.669328488212766 | 0.84 | 0.933 | 2.22394236119685e-281 |
| BCL2A1 | 6.3591259453308e-283 | 0.699293814179406 | 0.869 | 0.606 | 2.53239472520908e-278 |
| RP11-1143G9.4 | 7.49932282109156e-272 | 0.394995528912726 | 0.441 | 0.164 | 2.98645532704329e-267 |
| GLRX | 7.61824628711543e-267 | 0.72739825570578 | 0.749 | 0.479 | 3.03381421891798e-262 |
| RPS11 | 2.14546607809114e-261 | -0.639435623201682 | 0.91 | 0.945 | 8.54388956278234e-257 |
| RPL3 | 5.32442348134751e-261 | -0.734563519031282 | 0.906 | 0.95 | 2.12034516297702e-256 |
| PTMA | 7.81500909893739e-261 | -0.711989083150597 | 0.934 | 0.976 | 3.11217107346984e-256 |
| PLPP3 | 3.26145267608732e-260 | 0.337145965963117 | 0.33 | 0.101 | 1.29880829919825e-255 |
| RPS18 | 1.16013793390001e-256 | -0.742923860431246 | 0.981 | 0.982 | 4.62001729417e-252 |
| ATP2B1 | 5.21839628703044e-254 | 0.584734341905562 | 0.817 | 0.565 | 2.07812195338413e-249 |
| RAN | 9.74221326321659e-254 | 0.639327422922472 | 0.785 | 0.57 | 3.87964158781074e-249 |
| CCL3 | 3.59062417800298e-253 | 1.14313161740627 | 0.933 | 0.842 | 1.42989426640613e-248 |
| PALLD | 2.2857983136001e-252 | 0.414474183912481 | 0.295 | 0.086 | 9.10273462424968e-248 |
| CTSD | 2.10476130410263e-251 | 0.722257525289518 | 0.96 | 0.878 | 8.38179094132792e-247 |
| RPL7A | 4.64400087737681e-251 | -0.725559431761502 | 0.867 | 0.928 | 1.84938046939777e-246 |
| RPL10 | 3.39192536297535e-249 | -0.60137375362679 | 0.998 | 0.998 | 1.35076643729767e-244 |
| RPS16 | 5.54191933606268e-246 | -0.664290153806335 | 0.985 | 0.988 | 2.20695853720024e-241 |
| RPS4X | 4.40393169976275e-244 | -0.702077423894308 | 0.965 | 0.974 | 1.75377772079652e-239 |
| ARRDC4 | 1.96420844894061e-243 | 0.322343726726225 | 0.307 | 0.094 | 7.82206730621618e-239 |
| NOTCH2 | 2.58502831687886e-240 | 0.433503964461133 | 0.519 | 0.245 | 1.02943582663067e-235 |
| RP11-20G13.3 | 3.00630704217226e-239 | 0.348285088195866 | 0.177 | 0.032 | 1.19720165340426e-234 |
| CMPK2 | 3.49527851423642e-238 | 0.401350252771775 | 0.362 | 0.129 | 1.39192476272437e-233 |
| S100A6 | 1.36246967734837e-235 | 0.623858482731514 | 0.997 | 0.955 | 5.42576299610442e-231 |
| AMPD3 | 2.24929824538924e-235 | 0.476543981725811 | 0.562 | 0.291 | 8.95738040261356e-231 |
| RPL19 | 5.28870323906055e-234 | -0.61117052248428 | 0.979 | 0.986 | 2.10612029089108e-229 |
| HMOX1 | 4.66922707944688e-229 | 0.92065334591299 | 0.789 | 0.57 | 1.85942629984813e-224 |
| UBA52 | 2.13503493868484e-228 | -0.501927551901107 | 0.976 | 0.985 | 8.50234963632464e-224 |
| TDP2 | 1.57426659385808e-226 | 0.428187218354631 | 0.549 | 0.269 | 6.26920185672104e-222 |
| MIR155HG | 1.45971685082168e-224 | 0.61499568968347 | 0.54 | 0.275 | 5.81303041502716e-220 |
| ACTG1 | 1.56016853508962e-224 | -0.779659838854095 | 0.546 | 0.758 | 6.2130591572874e-220 |
| RABGEF1 | 5.65117324847772e-224 | 0.471604080362391 | 0.504 | 0.243 | 2.25046672274128e-219 |
| FTL | 4.91212214248208e-221 | 0.576079594489105 | 1 | 1 | 1.95615440080064e-216 |
| DDX60L | 8.16192583991667e-220 | 0.334388876487128 | 0.412 | 0.165 | 3.25032372723002e-215 |
| WTAP | 2.18946185978627e-218 | 0.592897352866917 | 0.873 | 0.709 | 8.71909396422686e-214 |
| AKR1B1 | 9.28980585007571e-218 | 0.587138013203782 | 0.665 | 0.392 | 3.69947938367565e-213 |
| SH3BP5 | 4.77663563319025e-215 | 0.477945548524746 | 0.592 | 0.333 | 1.90219960820535e-210 |
| RAB18 | 2.7732760160884e-214 | 0.409256061730442 | 0.522 | 0.263 | 1.10440170788689e-209 |
| RPS10 | 3.62230403474469e-211 | -0.584272776404022 | 0.473 | 0.717 | 1.44251013575638e-206 |
| TGFBI | 2.80145382461843e-210 | -0.742505190946529 | 0.145 | 0.428 | 1.1156229565778e-205 |
| PRNP | 4.39454638772631e-210 | 0.533771331005843 | 0.642 | 0.398 | 1.75004020798425e-205 |
| CD59 | 1.18192379930686e-209 | 0.512111356185888 | 0.728 | 0.481 | 4.70677514597969e-205 |
| GAPDH | 1.57505071924913e-209 | -0.794455133958192 | 0.991 | 0.997 | 6.27232447926581e-205 |
| TM4SF19 | 3.01196877109373e-209 | 0.255047920464149 | 0.216 | 0.055 | 1.19945632371266e-204 |
| CSTB | 1.96830529569335e-208 | 0.463404050867609 | 0.956 | 0.9 | 7.83838217903962e-204 |
| EREG | 1.00231365944747e-207 | 0.580924844161469 | 0.865 | 0.59 | 3.99151368601766e-203 |
| SOD2 | 6.0357234411163e-204 | 0.460853387402509 | 0.998 | 0.962 | 2.40360614595574e-199 |
| DPYSL3 | 1.90941247776341e-203 | 0.326952309472626 | 0.214 | 0.055 | 7.60385331019724e-199 |
| AKR1C2 | 6.84343898446701e-201 | 0.353962157546546 | 0.161 | 0.032 | 2.7252627067843e-196 |
| GADD45B | 8.0305240378018e-200 | -0.879434689250864 | 0.38 | 0.63 | 3.19799558757381e-195 |
| TSLP | 1.09922320364176e-199 | 0.415834554865141 | 0.147 | 0.026 | 4.37743656386258e-195 |
| BLOC1S2 | 3.7677025923779e-199 | 0.478579967659088 | 0.619 | 0.369 | 1.50041220336265e-194 |
| HLA-DRB6 | 2.19329168178674e-197 | -0.834389211735177 | 0.274 | 0.521 | 8.73434546437935e-193 |
| ARL8B | 2.11608367169563e-196 | 0.446691995012175 | 0.719 | 0.483 | 8.42688000579351e-192 |
| MMP10 | 4.56703285337249e-196 | 0.349517195579783 | 0.141 | 0.025 | 1.81872949319853e-191 |
| RPL23A | 9.83349403041018e-196 | -0.581962381817809 | 0.923 | 0.944 | 3.91599232773025e-191 |
| ASPH | 3.40309423418054e-194 | 0.405083398583024 | 0.504 | 0.249 | 1.35521421687772e-189 |
| RASGRP3 | 5.18118136905233e-194 | 0.604488476267419 | 0.586 | 0.356 | 2.06330185659771e-189 |
| JUNB | 1.92880238083351e-193 | -0.85336216676368 | 0.331 | 0.598 | 7.6810697211933e-189 |
| AKR1C1 | 3.46259877899948e-192 | 0.499181396155106 | 0.24 | 0.071 | 1.37891071176096e-187 |
| TNFAIP6 | 6.70346936233566e-190 | 0.48471145016618 | 0.565 | 0.281 | 2.66952260416293e-185 |
| DFNA5 | 2.7877890389345e-188 | 0.341563566696214 | 0.367 | 0.152 | 1.11018122897489e-183 |
| MT2A | 1.82082154772639e-187 | -1.47414669238247 | 0.846 | 0.895 | 7.25105764951082e-183 |
| BASP1 | 2.14204273883624e-186 | 0.480125830110538 | 0.907 | 0.703 | 8.53025679886758e-182 |
| MT-ATP6 | 8.12899361826989e-186 | -0.670409735602458 | 0.856 | 0.961 | 3.23720912860362e-181 |
| MT-ND1 | 1.11157302782425e-185 | -0.592671858817864 | 0.893 | 0.972 | 4.42661726870451e-181 |
| CFD | 1.616403265571e-183 | 0.698196000433698 | 0.691 | 0.471 | 6.4370027244834e-179 |
| OSBPL8 | 1.9776803190576e-183 | 0.572082200264332 | 0.657 | 0.421 | 7.87571633458306e-179 |
| TSC22D3 | 2.51377706507904e-183 | -0.699328151109391 | 0.178 | 0.445 | 1.00106144062643e-178 |
| PEA15 | 7.22280628391924e-183 | 0.463125556801881 | 0.612 | 0.377 | 2.87633814644516e-178 |
| ATP5G2 | 1.01323041938091e-182 | -0.525982589089552 | 0.341 | 0.599 | 4.03498749910061e-178 |
| HIVEP2 | 1.2980298155668e-182 | 0.286717900237922 | 0.326 | 0.124 | 5.16914413453167e-178 |
| NBPF19 | 1.38523728154546e-182 | 0.376057096018848 | 0.588 | 0.322 | 5.5164304262985e-178 |
| IL6 | 2.10889104380588e-182 | 0.695760953067568 | 0.431 | 0.198 | 8.39823680374814e-178 |
| DRAM1 | 2.37517372215398e-182 | 0.414101939688775 | 0.65 | 0.391 | 9.45865431373379e-178 |
| ARPC3 | 9.00141445802568e-182 | -0.495366907971607 | 0.644 | 0.818 | 3.58463327961957e-177 |
| PTGS2 | 7.20284125255982e-181 | 0.717245616044528 | 0.52 | 0.273 | 2.8683874720069e-176 |
| RNF13 | 2.05334544280248e-179 | 0.423080329805487 | 0.617 | 0.373 | 8.17703755687233e-175 |
| MT-ND5 | 3.87547003703878e-177 | -0.657822798210819 | 0.824 | 0.923 | 1.54332843284995e-172 |
| ATP13A3 | 1.20926660342715e-175 | 0.401335319589971 | 0.682 | 0.43 | 4.81566239482793e-171 |
| RPS24 | 4.48473428106298e-174 | -0.55881935771886 | 0.97 | 0.974 | 1.78595573274771e-169 |
| EIF4G2 | 1.503966245125e-173 | 0.453960622548003 | 0.823 | 0.659 | 5.98924477796127e-169 |
| SDS | 1.58077773228036e-173 | -1.16872892332669 | 0.104 | 0.34 | 6.29513116326009e-169 |
| RGS10 | 7.72172146644501e-173 | -0.591658107945626 | 0.181 | 0.426 | 3.07502113958239e-168 |
| GPNMB | 1.16425058542134e-172 | 0.765406430775301 | 0.73 | 0.618 | 4.63639510632339e-168 |
| SSTR2 | 7.01400634958732e-172 | 0.336176396980491 | 0.323 | 0.126 | 2.79318774859616e-167 |
| ANXA5 | 8.09372385575836e-172 | 0.332293152218451 | 0.972 | 0.854 | 3.22316365107865e-167 |
| UBE2H | 8.61465414067491e-172 | 0.34980767930819 | 0.451 | 0.222 | 3.43061371844097e-167 |
| RPL29 | 1.66682192445583e-170 | -0.524751849780928 | 0.958 | 0.964 | 6.63778494976045e-166 |
| POLR2L | 3.55730792846188e-170 | 0.415439275951129 | 0.772 | 0.549 | 1.41662673635137e-165 |
| RPS9 | 2.81055752026357e-169 | -0.441255460939071 | 0.991 | 0.988 | 1.11924832129456e-164 |
| LAPTM5 | 1.01837719860543e-168 | -0.535537065292785 | 0.939 | 0.94 | 4.0554835180064e-164 |
| LYZ | 2.91907979853386e-166 | 0.337308205973902 | 0.888 | 0.745 | 1.16246514817014e-161 |
| RHOF | 3.37786779685811e-166 | 0.396783529280885 | 0.449 | 0.225 | 1.3451682927428e-161 |
| NEDD4L | 4.33433894669153e-165 | 0.353122446213711 | 0.443 | 0.218 | 1.72606379874097e-160 |
| RPL6 | 1.95594250127203e-163 | -0.497477041263716 | 0.875 | 0.93 | 7.7891498228156e-159 |
| B2M | 4.09057042491365e-163 | 0.293343642449785 | 1 | 1 | 1.62898786031336e-158 |
| RPL5 | 6.0548695854002e-163 | -0.593994974259407 | 0.745 | 0.841 | 2.41123071499392e-158 |
| PPIA | 1.11631912142045e-162 | -0.51238311677824 | 0.717 | 0.826 | 4.44551763723264e-158 |
| RPL28 | 4.57151558377336e-162 | -0.477416181945594 | 0.997 | 0.995 | 1.82051465092607e-157 |
| EID3 | 4.41876651008391e-161 | 0.28464422608739 | 0.285 | 0.108 | 1.75968538731072e-156 |
| MIR4435-2HG | 1.89993076819301e-160 | 0.434564625112414 | 0.424 | 0.209 | 7.56609429817502e-156 |
| APOC1 | 4.69959611195208e-158 | 1.08651603090152 | 0.733 | 0.722 | 1.87152015966268e-153 |
| CALR | 1.08881254963881e-156 | -0.578678190963639 | 0.514 | 0.701 | 4.33597821642664e-152 |
| APOE | 2.28692209854133e-156 | 0.943375475467201 | 0.767 | 0.842 | 9.10720987302115e-152 |
| TBCA | 3.03141564848237e-156 | 0.400424064463923 | 0.685 | 0.467 | 1.20720065369513e-151 |
| PFDN2 | 8.13846391015794e-156 | 0.398494592847848 | 0.587 | 0.361 | 3.24098048294219e-151 |
| RPL4 | 7.51468703717288e-154 | -0.550588619748774 | 0.626 | 0.766 | 2.99257381881336e-149 |
| AC013461.1 | 7.60633615367058e-154 | 0.321186788837369 | 0.392 | 0.185 | 3.02907124647623e-149 |
| SDC2 | 3.04835161069078e-152 | 0.377380177628041 | 0.63 | 0.364 | 1.21394506192539e-147 |
| RPS3A | 5.43633422648748e-152 | -0.570782965097525 | 0.959 | 0.963 | 2.16491137901411e-147 |
| RGS2 | 5.60721653192534e-152 | -0.659745209245707 | 0.094 | 0.319 | 2.23296183950863e-147 |
| JUN | 2.46107391652973e-151 | 0.287379248356487 | 0.856 | 0.666 | 9.80073465779633e-147 |
| FNDC3B | 1.41808390879174e-150 | 0.376522892654716 | 0.636 | 0.409 | 5.64723554998136e-146 |
| NEAT1 | 3.11467231236582e-150 | -0.655836201150222 | 0.952 | 0.973 | 1.24035595495344e-145 |
| GADD45A | 7.45602814022355e-150 | 0.614092512656143 | 0.497 | 0.288 | 2.96921408628122e-145 |
| SMIM4 | 4.78181328014486e-148 | 0.328739183336217 | 0.417 | 0.211 | 1.90426150255209e-143 |
| AK4 | 7.14877316785396e-148 | 0.252250614508029 | 0.283 | 0.11 | 2.84685593863448e-143 |
| GSN | 8.66702565221241e-148 | -0.48989711550823 | 0.126 | 0.353 | 3.45146962548055e-143 |
| SLC25A6 | 1.70251599916071e-147 | -0.528873118358717 | 0.554 | 0.717 | 6.77992946345771e-143 |
| RPS3 | 1.99170276871753e-147 | -0.527062059496059 | 0.962 | 0.969 | 7.93155793586382e-143 |
| ZFP36 | 2.50741537824719e-147 | -0.694853470114431 | 0.167 | 0.402 | 9.98528026079378e-143 |
| SETD5 | 2.61528371478465e-146 | 0.355928697995884 | 0.332 | 0.149 | 1.04148443373869e-141 |
| ABCA1 | 1.39972617431419e-145 | 0.403327544691931 | 0.759 | 0.562 | 5.57412954397141e-141 |
| MT-CO3 | 1.70761601497548e-145 | -0.473528427220564 | 0.912 | 0.979 | 6.80023925643684e-141 |
| TNFSF14 | 2.68709462389822e-145 | 0.282564806049763 | 0.227 | 0.078 | 1.07008169207499e-140 |
| RPL10A | 3.51267965756318e-145 | -0.599604896248106 | 0.836 | 0.871 | 1.39885442003139e-140 |
| CCL7 | 1.59575132021735e-144 | 0.514488565198826 | 0.248 | 0.091 | 6.35476048250153e-140 |
| ID2 | 1.02778024654431e-143 | -0.935832844394626 | 0.428 | 0.617 | 4.09292927581339e-139 |
| EEF1A1 | 7.50265329829088e-143 | -0.408505069162731 | 1 | 0.999 | 2.98778162297838e-138 |
| ACTB | 1.0583011448681e-142 | -0.998260307249122 | 0.947 | 0.964 | 4.21447264920823e-138 |
| KLF4 | 7.3494560201038e-141 | -0.628849087673257 | 0.159 | 0.382 | 2.92677387088594e-136 |
| POMP | 1.48843803841407e-140 | 0.341368038426596 | 0.887 | 0.756 | 5.92740680037634e-136 |
| AHR | 1.41083727139139e-139 | 0.368748593533813 | 0.563 | 0.349 | 5.61837726586191e-135 |
| GOLGA7 | 1.64185358882326e-139 | 0.293162151516676 | 0.391 | 0.194 | 6.53835354677087e-135 |
| SLC38A2 | 3.02675523414576e-139 | 0.451391009945396 | 0.671 | 0.464 | 1.20534473689387e-134 |
| YWHAG | 3.98284851957627e-139 | 0.283846351455845 | 0.384 | 0.188 | 1.58608976595086e-134 |
| COTL1 | 4.32411673488619e-138 | -0.586632229629309 | 0.374 | 0.578 | 1.72199300733373e-133 |
| RPL15 | 4.78331774768949e-138 | -0.440534171476559 | 0.979 | 0.98 | 1.90486062666238e-133 |
| HK2 | 5.24503055227519e-138 | 0.300074112340322 | 0.438 | 0.23 | 2.08872851683255e-133 |
| ASAP1 | 6.64649710690178e-138 | 0.322815935840719 | 0.593 | 0.361 | 2.6468345428815e-133 |
| MMP12 | 9.92902232741898e-138 | 0.593717194161943 | 0.131 | 0.03 | 3.95403456144806e-133 |
| NACA | 4.16391132522362e-137 | -0.414988932311558 | 0.888 | 0.937 | 1.6581944070438e-132 |
| CHIT1 | 7.16433461200052e-137 | 0.32918651286635 | 0.149 | 0.038 | 2.85305297253697e-132 |
| TNIP1 | 1.0957573067564e-136 | 0.364577564062334 | 0.684 | 0.46 | 4.363634322696e-132 |
| SRP14 | 4.9815556657511e-136 | 0.336945445017855 | 0.959 | 0.908 | 1.98380491277206e-131 |
| S100A9 | 5.31543461984921e-136 | 0.335782001965616 | 0.791 | 0.52 | 2.11676552866255e-131 |
| UFM1 | 1.24100938142825e-135 | 0.336169944145273 | 0.58 | 0.359 | 4.94207165966171e-131 |
| ST3GAL1 | 5.45614916313725e-135 | 0.336445335803818 | 0.549 | 0.33 | 2.17280228123615e-130 |
| RAB7A | 4.16815013549503e-134 | 0.354483147739366 | 0.762 | 0.582 | 1.65988242845819e-129 |
| RPL13 | 5.73763642898059e-134 | -0.454772618400258 | 0.995 | 0.994 | 2.28489895511294e-129 |
| MT-CYB | 1.85871714656885e-133 | -0.481565017755451 | 0.875 | 0.963 | 7.40196929278114e-129 |
| MT-ND4 | 3.06095086915062e-133 | -0.484300646543733 | 0.934 | 0.98 | 1.21896246462185e-128 |
| FAM26F | 6.16505668390086e-133 | -0.410841996359497 | 0.066 | 0.263 | 2.45511052322984e-128 |
| GPX1 | 1.85348974045708e-132 | -0.50675893114826 | 0.86 | 0.92 | 7.38115219342221e-128 |
| AC002456.2 | 1.57857753392664e-131 | 0.273897991339423 | 0.196 | 0.066 | 6.28636931335606e-127 |
| GPR183 | 5.67810519579083e-131 | -0.709296292493175 | 0.459 | 0.638 | 2.26119183211978e-126 |
| MT-CO2 | 9.10613368141058e-131 | -0.512794299779354 | 0.935 | 0.98 | 3.62633561594813e-126 |
| MT-ND2 | 2.43609311601472e-130 | -0.471245224808266 | 0.938 | 0.979 | 9.70125361590542e-126 |
| MESDC1 | 3.54698326642787e-130 | 0.323288997662169 | 0.506 | 0.292 | 1.41251514618957e-125 |
| FAM107B | 1.06879050572627e-129 | 0.338165744510832 | 0.611 | 0.394 | 4.25624443095372e-125 |
| NRIP3 | 5.1670293569313e-129 | 0.273361671830156 | 0.413 | 0.214 | 2.05766610081075e-124 |
| CEBPD | 1.38884602946822e-128 | -0.594194164864152 | 0.2 | 0.415 | 5.53080154315129e-124 |
| PDE4DIP | 9.03524000807358e-127 | 0.378675517238241 | 0.746 | 0.577 | 3.59810362841514e-122 |
| PHLDA2 | 1.3410645581933e-126 | 0.366934719795714 | 0.496 | 0.286 | 5.34052139009318e-122 |
| CYLD | 9.94993988843593e-125 | 0.34209189894496 | 0.535 | 0.334 | 3.96236456177184e-120 |
| YWHAZ | 1.44684419742479e-124 | 0.356521503304021 | 0.818 | 0.666 | 5.76176764740476e-120 |
| PLAUR | 1.46774510155597e-124 | 0.36942449000218 | 0.964 | 0.859 | 5.84500131792632e-120 |
| NLRP3 | 3.62568174265961e-124 | -0.386996108068561 | 0.045 | 0.226 | 1.44385524037934e-119 |
| FGL2 | 4.24506750028028e-124 | -0.501121960642664 | 0.103 | 0.301 | 1.69051323063662e-119 |
| CSF3 | 4.26123939888769e-122 | 0.289977114444704 | 0.242 | 0.094 | 1.69695336581904e-117 |
| HLA-DPB1 | 1.2414496063986e-121 | -0.913160993888549 | 0.734 | 0.822 | 4.94382476756116e-117 |
| KMO | 2.98085555604397e-121 | 0.251117475117852 | 0.316 | 0.146 | 1.18706610808339e-116 |
| APLP2 | 6.72867515099767e-121 | 0.29242017568728 | 0.873 | 0.721 | 2.6795603053818e-116 |
| GNAI2 | 7.4876462112105e-121 | -0.392622894488313 | 0.247 | 0.465 | 2.98180535069036e-116 |
| CD74 | 8.6564379808583e-120 | -0.847221768771458 | 0.868 | 0.936 | 3.4472532971172e-115 |
| RPL9 | 1.86302993912507e-118 | -0.409352459482998 | 0.978 | 0.982 | 7.41914412657778e-114 |
| EEF1B2 | 2.17370574605202e-118 | -0.487014708894858 | 0.579 | 0.715 | 8.65634839250295e-114 |
| YWHAE | 2.46387988901454e-117 | 0.350688586729567 | 0.706 | 0.53 | 9.8119088820226e-113 |
| CYCS | 7.27509142134752e-117 | 0.38540017432408 | 0.719 | 0.548 | 2.89715965672322e-112 |
| CAPN2 | 9.76298196837625e-117 | 0.255105869376191 | 0.363 | 0.184 | 3.88791230926648e-112 |
| CKB | 1.38269548331522e-116 | 0.367713462655932 | 0.399 | 0.21 | 5.5063082232062e-112 |
| ACTR3 | 2.89972043869136e-115 | 0.296636112945434 | 0.612 | 0.403 | 1.15475567030006e-110 |
| RPL18 | 1.25832575769072e-114 | -0.4131591468478 | 0.953 | 0.964 | 5.01103066485175e-110 |
| MFSD2A | 1.01046728905017e-113 | 0.307816668234977 | 0.43 | 0.242 | 4.02398388518451e-109 |
| SLAMF7 | 1.18136103480272e-113 | 0.286246579624338 | 0.372 | 0.188 | 4.70453404889488e-109 |
| ARHGDIB | 1.35772251446034e-113 | -0.437231964167075 | 0.257 | 0.472 | 5.4068583693354e-109 |
| RAB10 | 3.45135278458482e-112 | 0.296748107897657 | 0.511 | 0.314 | 1.37443221940521e-107 |
| GLUL | 3.74598825419966e-112 | -0.792950864215328 | 0.511 | 0.656 | 1.49176490246993e-107 |
| RPL30 | 5.30717054795158e-112 | -0.425634065034877 | 0.984 | 0.977 | 2.11347452731076e-107 |
| PER1 | 1.32607932045852e-110 | -0.285204517383305 | 0.048 | 0.217 | 5.28084567786195e-106 |
| RPS2 | 3.7512148448003e-110 | -0.486918965236864 | 0.986 | 0.986 | 1.49384628764482e-105 |
| PLD3 | 5.49494232239672e-110 | 0.438718557494827 | 0.725 | 0.594 | 2.18825088104805e-105 |
| FOSB | 9.69786957199595e-110 | -0.504693649504603 | 0.151 | 0.347 | 3.86198259965595e-105 |
| RPL26 | 1.68786747822203e-109 | -0.443009631972685 | 0.96 | 0.961 | 6.72159465852358e-105 |
| LGALS9 | 1.72783193882795e-109 | -0.37153087468095 | 0.105 | 0.286 | 6.88074512999456e-105 |
| EIF3J | 1.96403153900567e-109 | 0.301447098685105 | 0.403 | 0.225 | 7.82136279778229e-105 |
| RPL36A | 3.44414194146468e-109 | -0.475051703994514 | 0.307 | 0.503 | 1.37156064534948e-104 |
| FNIP2 | 8.48871402149737e-109 | 0.336556275663435 | 0.814 | 0.652 | 3.3804605847809e-104 |
| DYNLL1 | 1.05715391841977e-108 | 0.432423336719838 | 0.695 | 0.489 | 4.20990404932306e-104 |
| RPL27A | 5.66736448863538e-108 | -0.434723867368328 | 0.98 | 0.977 | 2.25691456030927e-103 |
| USP12 | 7.39696205440806e-108 | 0.383614451679955 | 0.699 | 0.536 | 2.94569219892692e-103 |
| PIK3CB | 8.48310414689508e-108 | 0.250312445512842 | 0.392 | 0.213 | 3.37822656441803e-103 |
| RIT1 | 3.01678032004567e-107 | 0.322126632548131 | 0.58 | 0.387 | 1.20137242685179e-102 |
| UBC | 3.35582707452646e-107 | -0.48723582803903 | 0.969 | 0.979 | 1.33639101588867e-102 |
| KLF2 | 5.09264345971343e-107 | -0.560268601133702 | 0.048 | 0.212 | 2.02804340496168e-102 |
| MRPS6 | 5.56026521562382e-107 | 0.314046315032898 | 0.473 | 0.273 | 2.21426441681787e-102 |
| MCL1 | 6.84760842646713e-107 | -0.418531128882847 | 0.388 | 0.575 | 2.726923103672e-102 |
| CD82 | 7.78303567888511e-106 | 0.26720280917741 | 0.472 | 0.277 | 3.09943829840242e-101 |
| TSC22D1 | 1.76300594284832e-105 | 0.492533754697441 | 0.595 | 0.412 | 7.02081856620485e-101 |
| TUBB | 5.12803598875911e-105 | -0.472479064474336 | 0.319 | 0.515 | 2.04213777180354e-100 |
| RPL13A | 5.45629609805234e-105 | -0.42862858223198 | 0.995 | 0.988 | 2.17286079512738e-100 |
| CASP1 | 8.12867325727193e-105 | 0.291941429264826 | 0.405 | 0.226 | 3.2370815512434e-100 |
| RPSA | 1.14056461007475e-104 | -0.554880401265263 | 0.709 | 0.784 | 4.54207044670068e-100 |
| PPIF | 1.21219093221495e-104 | 0.332250578434876 | 0.644 | 0.451 | 4.82730794935959e-100 |
| CSF2 | 1.40814467719633e-104 | 0.285967895634176 | 0.131 | 0.038 | 5.60765454799896e-100 |
| RPL31 | 1.51359684769546e-104 | -0.42391020258224 | 0.957 | 0.957 | 6.02759672657761e-100 |
| PTPN12 | 2.55746280523875e-104 | 0.277732336725435 | 0.524 | 0.329 | 1.01845841293023e-99 |
| LUCAT1 | 7.3086338651421e-103 | 0.263852636787996 | 0.36 | 0.188 | 2.91051726411554e-98 |
| HMGN2 | 1.01487483943596e-102 | -0.48502003867952 | 0.268 | 0.458 | 4.04153607308583e-98 |
| PNPLA8 | 1.76220605477395e-102 | 0.310792230410094 | 0.561 | 0.374 | 7.01763317192631e-98 |
| HERPUD1 | 3.18075828426478e-102 | -0.510579839831059 | 0.238 | 0.429 | 1.26667337154277e-97 |
| NBN | 4.90631793290138e-102 | 0.349224327606441 | 0.44 | 0.261 | 1.95384299041932e-97 |
| EIF5B | 1.63675366288303e-101 | 0.277622872954684 | 0.431 | 0.252 | 6.51804411169909e-97 |
| CGGBP1 | 2.19108470517185e-101 | 0.256624882720821 | 0.406 | 0.231 | 8.72555662140587e-97 |
| IER2 | 4.75138835660335e-101 | -0.622746291313693 | 0.271 | 0.462 | 1.89214538525015e-96 |
| HIF1A-AS2 | 4.89989552417955e-101 | 0.379961403061386 | 0.283 | 0.135 | 1.95128539459402e-96 |
| CSGALNACT2 | 6.21923251602976e-101 | 0.283290810526862 | 0.532 | 0.344 | 2.47668496485853e-96 |
| RPLP2 | 1.75032110183022e-100 | -0.366320614163356 | 0.992 | 0.989 | 6.9703037238185e-96 |
| PTP4A1 | 1.87956569055388e-100 | 0.340249510046263 | 0.554 | 0.375 | 7.48499444949271e-96 |
| IL1RN | 3.58918174538601e-99 | 0.63053289015697 | 0.613 | 0.432 | 1.42931984646507e-94 |
| TCEB1 | 5.00378527156349e-99 | 0.316699865732177 | 0.724 | 0.553 | 1.99265740869473e-94 |
| RHEB | 1.37652675353979e-98 | 0.311516560902075 | 0.786 | 0.625 | 5.48174249062151e-94 |
| RPL14 | 2.117844040082e-97 | -0.40428954290524 | 0.907 | 0.933 | 8.43389032081853e-93 |
| CCL2 | 2.7664207945162e-97 | 1.24558277096022 | 0.337 | 0.181 | 1.10167175300019e-92 |
| ZFP36L2 | 2.8325798448422e-97 | -0.546072323857167 | 0.171 | 0.346 | 1.12801827161151e-92 |
| CCNL1 | 6.34949705853877e-97 | -0.459095066255077 | 0.461 | 0.616 | 2.5285602136219e-92 |
| IFI6 | 7.99654686654295e-97 | 0.532932815519513 | 0.723 | 0.586 | 3.1844648586634e-92 |
| SGK1 | 1.21706883416452e-96 | -0.734788509601889 | 0.328 | 0.498 | 4.84673321829336e-92 |
| HMGA1 | 8.35890249067765e-96 | 0.309457859640153 | 0.565 | 0.377 | 3.32876573886256e-91 |
| FGR | 9.8900293944938e-96 | 0.26768760481395 | 0.521 | 0.335 | 3.93850640576927e-91 |
| SEC61B | 2.40907036495765e-95 | 0.307504267546369 | 0.899 | 0.789 | 9.59364091437085e-91 |
| RPL24 | 3.60461334821643e-95 | -0.36467876487978 | 0.921 | 0.938 | 1.43546517366023e-90 |
| CD164 | 3.94417944249651e-95 | 0.287333125456997 | 0.574 | 0.393 | 1.57069057938538e-90 |
| RHOB | 7.29190925907361e-95 | -0.787226736724335 | 0.121 | 0.286 | 2.90385702424088e-90 |
| RNASE1 | 1.52412731260101e-94 | -0.928263202071068 | 0.54 | 0.634 | 6.06953219697099e-90 |
| RHOC | 1.64485673270309e-94 | 0.328403676545209 | 0.622 | 0.447 | 6.5503129666435e-90 |
| CAPZB | 1.73061389579893e-94 | -0.35418721766767 | 0.398 | 0.57 | 6.89182371724008e-90 |
| PHACTR1 | 4.75399302040107e-94 | 0.282519543410908 | 0.508 | 0.32 | 1.89318264051432e-89 |
| LDHA | 3.91618108733799e-93 | -0.522478203393361 | 0.609 | 0.721 | 1.55954079441061e-88 |
| HLA-C | 6.00776967098745e-93 | 0.349085232783862 | 0.989 | 0.981 | 2.39247411607733e-88 |
| HLA-DPA1 | 9.65146591654126e-93 | -0.900253701483255 | 0.735 | 0.811 | 3.84350327194422e-88 |
| PELI1 | 1.02176412886535e-92 | -0.542142334890892 | 0.292 | 0.448 | 4.0689712903805e-88 |
| RCSD1 | 8.75419061741894e-92 | 0.302823954288514 | 0.302 | 0.157 | 3.48618132957474e-87 |
| RPS14 | 3.64812116790784e-91 | -0.367597264517283 | 0.992 | 0.991 | 1.45279129269594e-86 |
| PTPRC | 6.93962106567213e-91 | -0.315583173764034 | 0.118 | 0.289 | 2.76356529698261e-86 |
| SPINK1 | 1.99581569552776e-90 | 0.270363738904671 | 0.157 | 0.056 | 7.94793684430021e-86 |
| IL18 | 2.45011809944257e-90 | -0.264185113194678 | 0.068 | 0.222 | 9.75710530741016e-86 |
| HNRNPAB | 2.18632623567115e-89 | 0.258141426609312 | 0.39 | 0.227 | 8.70660696831321e-85 |
| JUND | 3.08083782402802e-89 | -0.463944718517276 | 0.343 | 0.511 | 1.22688204666268e-84 |
| SEC62 | 1.92047314281646e-88 | 0.337516688677832 | 0.721 | 0.542 | 7.64790019663797e-84 |
| RPS26 | 4.93737244346088e-88 | -0.348648762770102 | 0.736 | 0.834 | 1.96620982815943e-83 |
| SAMD9 | 1.19957845369691e-87 | 0.26600571133072 | 0.317 | 0.169 | 4.77708127615722e-83 |
| SMS | 1.33212723079053e-87 | 0.307540548471767 | 0.644 | 0.475 | 5.30493027117711e-83 |
| PMAIP1 | 1.38103517152326e-87 | -0.815814988361331 | 0.268 | 0.424 | 5.49969636355707e-83 |
| ARPC1B | 1.39023916354992e-87 | -0.357752393882029 | 0.461 | 0.626 | 5.53634942100483e-83 |
| GLTSCR2 | 1.52886595209543e-87 | -0.345097596992474 | 0.338 | 0.508 | 6.08840288102963e-83 |
| SNRPG | 2.63231007531426e-87 | 0.26145626543849 | 0.599 | 0.412 | 1.0482648412924e-82 |
| RPL32 | 3.30372393708487e-87 | -0.376635131093111 | 0.989 | 0.987 | 1.31564198346531e-82 |
| SPI1 | 4.46172908928201e-87 | -0.330160733900684 | 0.394 | 0.587 | 1.77679437522477e-82 |
| DSE | 5.27493360228141e-87 | 0.287502325046715 | 0.58 | 0.405 | 2.10063680843653e-82 |
| NENF | 2.2188755903156e-86 | 0.290471033630154 | 0.565 | 0.402 | 8.83622826331383e-82 |
| SERPINF1 | 1.42099770361266e-85 | -0.30091452309213 | 0.044 | 0.182 | 5.6588391550967e-81 |
| PSMA7 | 1.44913425721302e-85 | 0.286044516226272 | 0.864 | 0.753 | 5.77088735249941e-81 |
| LAMTOR5 | 3.26212969945422e-85 | 0.259267151984513 | 0.594 | 0.414 | 1.29907791021365e-80 |
| F3 | 5.60842973081137e-85 | 0.287903054419602 | 0.254 | 0.12 | 2.23344497170101e-80 |
| SUB1 | 8.65702092745744e-85 | 0.30095262030379 | 0.811 | 0.672 | 3.44748544394137e-80 |
| PPIB | 8.85147804881962e-85 | -0.35678538064189 | 0.488 | 0.642 | 3.52492410338144e-80 |
| DSTN | 1.6793843109133e-84 | 0.292316647017311 | 0.498 | 0.328 | 6.68781214135002e-80 |
| RPL27 | 2.68949341572363e-84 | -0.363296283960207 | 0.943 | 0.943 | 1.07103696294362e-79 |
| HSPB1 | 3.09722677179901e-84 | -1.13750262096767 | 0.232 | 0.394 | 1.23340861733352e-79 |
| FAU | 4.51630107169683e-84 | -0.267219206941506 | 0.992 | 0.991 | 1.79852657578183e-79 |
| INHBA | 7.92081129505729e-84 | 0.40118095808928 | 0.323 | 0.176 | 3.15430468203066e-79 |
| CD52 | 2.1065479673833e-83 | 0.286264165506295 | 0.366 | 0.206 | 8.38890597051053e-79 |
| THUMPD3-AS1 | 3.66990290467192e-83 | 0.2532857119574 | 0.37 | 0.214 | 1.4614654337275e-78 |
| ATOX1 | 4.10551684192585e-83 | 0.293008258817962 | 0.749 | 0.599 | 1.63493997196013e-78 |
| S100A8 | 4.8903089510422e-83 | 0.754600147014454 | 0.465 | 0.302 | 1.94746773357354e-78 |
| CSF1R | 6.72590475938048e-83 | -0.29710140907344 | 0.05 | 0.187 | 2.67845705232809e-78 |
| FCGR2A | 7.57798868500981e-83 | -0.476559855893915 | 0.555 | 0.651 | 3.01778243403146e-78 |
| STX3 | 1.21298891850896e-82 | 0.263336376088035 | 0.428 | 0.267 | 4.83048577017821e-78 |
| RPS8 | 2.24257880475064e-82 | -0.391310583139419 | 0.957 | 0.963 | 8.93062157415849e-78 |
| EIF3H | 3.27570010684162e-82 | -0.291633449505029 | 0.27 | 0.445 | 1.30448205354754e-77 |
| ITGAV | 4.7451229053087e-82 | 0.268311175493432 | 0.47 | 0.303 | 1.88965029458108e-77 |
| RPS7 | 5.76919737211689e-82 | -0.354905050829774 | 0.956 | 0.963 | 2.29746746949811e-77 |
| BZW1 | 6.92403637656165e-82 | 0.259583391238554 | 0.736 | 0.573 | 2.75735900623815e-77 |
| STAB1 | 9.12008151510844e-82 | -0.370057176629153 | 0.048 | 0.182 | 3.63189006176163e-77 |
| ATP5D | 4.61271237948668e-81 | -0.294149140357101 | 0.262 | 0.434 | 1.83692045088298e-76 |
| NUDT4 | 2.18276997790653e-80 | 0.25512110555292 | 0.344 | 0.196 | 8.69244488301718e-76 |
| NMB | 9.44075957823546e-80 | 0.292816818755955 | 0.279 | 0.144 | 3.75959368684071e-75 |
| LST1 | 3.57674072146082e-79 | -0.469599898868061 | 0.348 | 0.483 | 1.42436545750734e-74 |
| RAB20 | 2.26197807531843e-78 | -0.282827026051803 | 0.158 | 0.318 | 9.0078752893406e-74 |
| TMEM123 | 8.70657190859048e-78 | 0.290937040299528 | 0.569 | 0.408 | 3.46721813115799e-73 |
| EIF1AX | 9.72312892251292e-78 | 0.257170055992679 | 0.503 | 0.336 | 3.87204163081232e-73 |
| EIF3F | 1.78328161202427e-77 | -0.283876178105399 | 0.215 | 0.38 | 7.10156236356424e-73 |
| NINJ1 | 2.38627849122539e-77 | 0.283673360207434 | 0.936 | 0.837 | 9.50287683560687e-73 |
| SPP1 | 5.21671699376137e-77 | 0.681238346353189 | 0.734 | 0.614 | 2.07745320842559e-72 |
| EIF3K | 1.00825237313999e-76 | -0.336090495406535 | 0.362 | 0.51 | 4.01516342555537e-72 |
| RPS15A | 2.69447193773524e-76 | -0.352049484183283 | 0.984 | 0.983 | 1.07301955976431e-71 |
| ATP5O | 5.62465720260215e-76 | -0.290408108142103 | 0.277 | 0.442 | 2.23990723779225e-71 |
| NDUFS5 | 6.35161062812541e-76 | -0.288146269412533 | 0.413 | 0.584 | 2.52940190043838e-71 |
| OAZ1 | 6.74969090961558e-76 | 0.258845035427099 | 0.997 | 0.985 | 2.68792941093621e-71 |
| EEF1D | 2.23951464528634e-75 | -0.306626811994149 | 0.774 | 0.838 | 8.9184191719238e-71 |
| SEC61G | 4.57822246733874e-75 | 0.264174653308464 | 0.852 | 0.707 | 1.82318553316831e-70 |
| FUS | 7.31062560877503e-75 | -0.375310041068465 | 0.465 | 0.596 | 2.91131043618248e-70 |
| CXCL5 | 9.38883609824447e-75 | 0.768146396624779 | 0.319 | 0.182 | 3.7389161994039e-70 |
| ALDH2 | 1.77157136639117e-74 | -0.306413668308309 | 0.168 | 0.323 | 7.05492865237957e-70 |
| TPT1 | 5.40106804952732e-74 | -0.356601585128699 | 1 | 1 | 2.15086732936327e-69 |
| SPAG9 | 1.06059651308263e-73 | 0.275488881650465 | 0.621 | 0.467 | 4.22361349404896e-69 |
| PSME1 | 1.53481838406859e-73 | -0.315981763362063 | 0.33 | 0.483 | 6.11210725087636e-69 |
| IFITM2 | 1.73884940254972e-72 | -0.362368102568872 | 0.173 | 0.32 | 6.92461997577374e-68 |
| PYCARD | 3.31621771755555e-72 | -0.293866153795042 | 0.135 | 0.282 | 1.32061738166215e-67 |
| AIF1 | 3.92593483110591e-72 | -0.847656089346293 | 0.634 | 0.647 | 1.56342502779131e-67 |
| RCAN1 | 6.95888964226123e-72 | 0.253405777537946 | 0.216 | 0.104 | 2.77123862223769e-67 |
| MCEMP1 | 7.56438930980997e-72 | 0.270199599459992 | 0.164 | 0.07 | 3.01236675484562e-67 |
| EVI2B | 9.65146221163941e-72 | -0.256278481234182 | 0.161 | 0.323 | 3.84350179654116e-67 |
| C14orf2 | 1.27924582665163e-71 | 0.260423036086493 | 0.762 | 0.621 | 5.09434065547478e-67 |
| LAIR1 | 1.8920373555525e-71 | -0.258075933217883 | 0.171 | 0.327 | 7.53466036101674e-67 |
| EMP1 | 2.38055714020061e-71 | 0.272422552790336 | 0.507 | 0.345 | 9.48009269942088e-67 |
| GNAS | 2.43285304277984e-71 | -0.355467267671694 | 0.506 | 0.631 | 9.68835067226216e-67 |
| UBALD2 | 4.11490163393599e-71 | -0.294046236531656 | 0.204 | 0.359 | 1.63867727768233e-66 |
| SRRM1 | 7.67717403967894e-71 | -0.286489845313623 | 0.313 | 0.471 | 3.05728101782134e-66 |
| RPS5 | 1.73719442461064e-70 | -0.385346292758374 | 0.877 | 0.889 | 6.91802935712696e-66 |
| UCP2 | 2.789721338596e-70 | -0.267841824350265 | 0.091 | 0.228 | 1.11095072866909e-65 |
| RPL18A | 8.84333663755145e-70 | -0.378971761248994 | 0.976 | 0.973 | 3.52168194917211e-65 |
| RPS23 | 1.74036052165721e-69 | -0.382796916976236 | 0.957 | 0.961 | 6.93063770539551e-65 |
| GRB2 | 4.52165567794392e-68 | -0.323894099038901 | 0.406 | 0.547 | 1.80065894062761e-63 |
| RPLP0 | 6.16635687931348e-68 | -0.34142041653046 | 0.836 | 0.876 | 2.45562830004901e-63 |
| RPL11 | 7.44134140067995e-68 | -0.338753083795035 | 0.99 | 0.99 | 2.96336538599277e-63 |
| RGS1 | 3.40057070605794e-67 | -0.984144179997628 | 0.365 | 0.493 | 1.35420927227345e-62 |
| SLC25A5 | 7.20181464975542e-67 | -0.327396730129803 | 0.306 | 0.45 | 2.8679786479721e-62 |
| VAMP5 | 1.56751404840834e-66 | -0.335895996797907 | 0.117 | 0.251 | 6.24231119497655e-62 |
| RPS13 | 3.6695078109093e-66 | -0.329676760616649 | 0.982 | 0.975 | 1.46130809553841e-61 |
| MT-CO1 | 5.05798815730167e-66 | -0.316582775108954 | 0.911 | 0.976 | 2.01424262388224e-61 |
| FN1 | 5.45476881739257e-66 | -0.987844219851517 | 0.206 | 0.344 | 2.17225258615024e-61 |
| COX4I1 | 6.03890001918157e-66 | -0.257411875519104 | 0.894 | 0.92 | 2.40487115463868e-61 |
| ZNF331 | 7.13879715670876e-66 | -0.263489255996268 | 0.063 | 0.185 | 2.84288319171613e-61 |
| CYBB | 7.34435004716417e-66 | -0.389480728479646 | 0.186 | 0.331 | 2.92474051928219e-61 |
| LAPTM4A | 9.55185605810657e-66 | -0.303743072850091 | 0.494 | 0.624 | 3.80383563801978e-61 |
| SLC5A3 | 1.07041143001044e-65 | 0.297099330693812 | 0.281 | 0.158 | 4.26269943773057e-61 |
| LRRC75A-AS1 | 1.21183525972263e-65 | -0.363549330072252 | 0.564 | 0.66 | 4.82589155479341e-61 |
| PHLDA1 | 1.8024575144067e-65 | 0.475314903669876 | 0.63 | 0.497 | 7.17792655962181e-61 |
| P4HB | 3.99921019650529e-65 | -0.335293072244966 | 0.417 | 0.556 | 1.5926054765543e-60 |
| CFL1 | 6.24246629331616e-65 | -0.302836652668868 | 0.958 | 0.946 | 2.48593735198729e-60 |
| LMNA | 6.50107069636222e-65 | -0.55940662744866 | 0.55 | 0.633 | 2.58892138341233e-60 |
| HMGB2 | 1.05162769875149e-64 | -0.295949307363197 | 0.149 | 0.293 | 4.18789698473804e-60 |
| IGSF6 | 2.23682458915696e-64 | -0.294297054737687 | 0.091 | 0.22 | 8.90770656139977e-60 |
| CCL4L2 | 2.29019600518073e-64 | -1.46800885670496 | 0.322 | 0.438 | 9.12024755143121e-60 |
| C1QB | 2.93825775049928e-64 | -0.776064316589717 | 0.647 | 0.739 | 1.17010238398133e-59 |
| SRSF5 | 4.83366790003612e-64 | -0.33255809018529 | 0.593 | 0.692 | 1.92491156783138e-59 |
| HCLS1 | 9.39340519011096e-64 | -0.257274425427968 | 0.159 | 0.299 | 3.74073574885789e-59 |
| HNRNPDL | 1.33703039740165e-63 | -0.281828613726539 | 0.479 | 0.612 | 5.32445615157259e-59 |
| PGK1 | 2.36717051086563e-63 | -0.355648086770215 | 0.396 | 0.531 | 9.42678312542022e-59 |
| UBE2D1 | 3.01242380365835e-63 | 0.287925143192739 | 0.745 | 0.607 | 1.19963753133087e-58 |
| COX7C | 3.67824552908677e-62 | -0.285975751361005 | 0.729 | 0.8 | 1.46478771704823e-57 |
| RBM39 | 4.49715226721625e-62 | -0.319619629038463 | 0.532 | 0.647 | 1.79090094737353e-57 |
| RPS19 | 5.82707110307126e-62 | -0.290758790345381 | 0.995 | 0.994 | 2.32051452537607e-57 |
| RPL37 | 2.40397453187149e-61 | -0.311558202092394 | 0.982 | 0.979 | 9.57334777827183e-57 |
| VIM | 3.59986244685479e-61 | -0.403924924868672 | 0.996 | 0.993 | 1.43357322221098e-56 |
| HNRNPA1 | 1.60618457855929e-60 | -0.337131792616036 | 0.661 | 0.741 | 6.39630884719665e-56 |
| MZT2B | 1.57755964329704e-59 | -0.304904210905704 | 0.309 | 0.447 | 6.28231576750178e-55 |
| ATP5H | 2.37734844158904e-59 | -0.266276193394208 | 0.292 | 0.432 | 9.46731469894003e-55 |
| RPS6 | 1.80078025937363e-58 | -0.36005457868276 | 0.982 | 0.979 | 7.17124722690359e-54 |
| FOS | 2.9956876349208e-58 | -0.656116148593822 | 0.251 | 0.406 | 1.19297268685451e-53 |
| PABPC4 | 3.00355652959591e-58 | -0.338777897912594 | 0.42 | 0.535 | 1.19610631678098e-53 |
| RPL8 | 6.56702803846642e-57 | -0.313480954951002 | 0.983 | 0.981 | 2.61518757575848e-52 |
| HLA-DRA | 7.11477327285809e-56 | -0.503885792234235 | 0.861 | 0.927 | 2.83331616045028e-51 |
| IRF1 | 1.8026419619939e-55 | -0.356881882551637 | 0.298 | 0.428 | 7.17866108524832e-51 |
| IRF8 | 2.71446314049848e-55 | -0.267407049102469 | 0.123 | 0.244 | 1.08098065644071e-50 |
| RPS12 | 3.41737267246024e-55 | -0.350698979785242 | 0.971 | 0.978 | 1.36090031935384e-50 |
| TYMP | 3.53792789367495e-55 | -0.384851476386831 | 0.692 | 0.748 | 1.40890902509817e-50 |
| RND3 | 3.5584058109979e-55 | 0.297501296519893 | 0.183 | 0.093 | 1.41706394611369e-50 |
| VCAN | 7.01932282902009e-55 | 0.441850987368859 | 0.341 | 0.22 | 2.79530493020067e-50 |
| TPI1 | 2.9336622104821e-54 | -0.354156411204632 | 0.643 | 0.722 | 1.16827230208029e-49 |
| RPS27A | 1.579766360518e-53 | -0.263220548208237 | 0.995 | 0.992 | 6.29110357749085e-49 |
| HLA-DMA | 1.11642064965374e-52 | -0.458056650391783 | 0.542 | 0.609 | 4.4459219531161e-48 |
| NR4A2 | 1.62586476110335e-52 | -0.27517684341147 | 0.151 | 0.275 | 6.47468123814187e-48 |
| GNA13 | 2.2485918821958e-52 | -0.280346064777814 | 0.261 | 0.384 | 8.95456745246835e-48 |
| TUBA1A | 5.92817699083214e-52 | -0.254016456242952 | 0.113 | 0.231 | 2.36077792305908e-47 |
| CSRNP1 | 1.28939983715675e-51 | -0.270060269034599 | 0.205 | 0.33 | 5.13477697150932e-47 |
| HMGN1 | 1.65626038908345e-51 | -0.260032013959459 | 0.342 | 0.47 | 6.59572574744703e-47 |
| MT-TE | 3.26794379809017e-50 | -0.331751692357445 | 0.101 | 0.213 | 1.30139325871345e-45 |
| HLA-DRB1 | 4.28015532480066e-50 | -0.567548918479968 | 0.838 | 0.897 | 1.70448625499537e-45 |
| EEF2 | 1.85912039505563e-49 | -0.263405937999848 | 0.607 | 0.696 | 7.40357514923003e-45 |
| GAS5 | 2.09215224268794e-49 | -0.267346988111187 | 0.339 | 0.466 | 8.33157787605618e-45 |
| INSIG1 | 4.68670908120504e-48 | 0.385141965948348 | 0.497 | 0.377 | 1.86638815740828e-43 |
| C5AR1 | 4.19116054813592e-46 | -0.265026457685577 | 0.361 | 0.491 | 1.66904586508417e-41 |
| RPS21 | 1.52391661041678e-45 | -0.290412732027328 | 0.875 | 0.88 | 6.06869311766273e-41 |
| RRAD | 1.83683143577887e-45 | 0.318943985094821 | 0.19 | 0.103 | 7.31481382670221e-41 |
| EIF1B | 2.06212847921403e-45 | 0.311505587941189 | 0.587 | 0.475 | 8.21201424277403e-41 |
| CDKN1A | 2.28243638275751e-45 | 0.259790829876779 | 0.685 | 0.582 | 9.08934640705525e-41 |
| KLF6 | 4.66299316415077e-45 | -0.378999092337446 | 0.869 | 0.903 | 1.85694376775976e-40 |
| C1QC | 5.86359864754241e-45 | -0.745877015567496 | 0.708 | 0.74 | 2.33506088941082e-40 |
| HLA-E | 1.34466685522566e-44 | -0.264208183345234 | 0.887 | 0.893 | 5.35486681756516e-40 |
| SEPP1 | 3.17274084190703e-43 | -0.611932019986787 | 0.169 | 0.27 | 1.26348058547264e-38 |
| HLA-DMB | 8.09653700166722e-43 | -0.346440968406503 | 0.254 | 0.359 | 3.22428393017394e-38 |
| MARCKSL1 | 1.90303333667547e-42 | 0.27299502870902 | 0.363 | 0.254 | 7.5784496566427e-38 |
| DUSP2 | 2.26418089192758e-42 | -0.469558348723411 | 0.167 | 0.272 | 9.01664756592322e-38 |
| OTUD1 | 5.86112153787682e-42 | -0.266853955683689 | 0.201 | 0.311 | 2.33407443002869e-37 |
| A2M | 9.84886874906359e-42 | -0.350740820948318 | 0.281 | 0.384 | 3.9221150019396e-37 |
| VAMP8 | 6.45686715062119e-41 | -0.346518275681509 | 0.37 | 0.47 | 2.57131820539188e-36 |
| ENO1 | 2.29273116710572e-40 | -0.368054434285848 | 0.699 | 0.73 | 9.13034332676513e-36 |
| PTGES | 3.05587278781875e-40 | 0.259604207931778 | 0.168 | 0.09 | 1.21694022029306e-35 |
| RPL12 | 3.45454603165554e-40 | -0.251520288361611 | 0.977 | 0.98 | 1.37570386618618e-35 |
| TSPO | 1.71316365548796e-39 | -0.300491416196224 | 0.508 | 0.584 | 6.82233162524969e-35 |
| C1QA | 2.95146049796455e-39 | -0.627712548550736 | 0.718 | 0.768 | 1.17536011410442e-34 |
| CXCL16 | 3.64322234580867e-39 | -0.258164462999302 | 0.484 | 0.565 | 1.45084043477139e-34 |
| CTSS | 9.98725926616579e-39 | 0.257450768945805 | 0.831 | 0.842 | 3.9772262575652e-34 |
| HLA-DQB1 | 1.03553130916883e-38 | -0.502429948856785 | 0.709 | 0.754 | 4.12379633250301e-34 |
| MS4A6A | 1.30073903222173e-38 | -0.47767379190858 | 0.304 | 0.388 | 5.17993304801661e-34 |
| SAMSN1 | 3.87355076546233e-38 | -0.258768782751596 | 0.449 | 0.562 | 1.54256412133006e-33 |
| MT1E | 1.01649252135357e-37 | -0.397596531474502 | 0.143 | 0.246 | 4.04797816778632e-33 |
| RPL7 | 1.12067188883857e-37 | -0.301631099670336 | 0.949 | 0.928 | 4.46285166292185e-33 |
| EIF4A3 | 1.97883775955094e-37 | -0.273017485726584 | 0.168 | 0.271 | 7.88032560985972e-33 |
| ALDOA | 4.2737789893585e-37 | -0.264317566834099 | 0.85 | 0.862 | 1.70194700693224e-32 |
| FCER1G | 5.26441022897379e-37 | -0.333546034687006 | 0.981 | 0.962 | 2.09644608548423e-32 |
| MT1G | 6.06724282264588e-37 | -0.496120487764893 | 0.092 | 0.183 | 2.41615810926227e-32 |
| IFI27 | 6.36446693069221e-37 | 0.411509025924719 | 0.555 | 0.425 | 2.53452166580956e-32 |
| RBM3 | 3.1924492888803e-36 | -0.250502795499948 | 0.533 | 0.606 | 1.2713290803108e-31 |
| CD14 | 6.10491828690589e-35 | -0.28591006294071 | 0.453 | 0.584 | 2.43116160939453e-30 |
| CLEC2B | 1.04915361365223e-34 | -0.254527902054989 | 0.268 | 0.369 | 4.17804443564727e-30 |
| MT-ND3 | 2.75848623400086e-34 | -0.323563734313396 | 0.895 | 0.961 | 1.09851197296616e-29 |
| BNIP3L | 4.32305381732777e-33 | -0.250887108090291 | 0.382 | 0.48 | 1.72156972167444e-28 |
| IFIT2 | 5.33952903458857e-32 | -0.437329000679186 | 0.051 | 0.12 | 2.12636064744421e-27 |
| BST2 | 8.16207861682668e-31 | -0.26173028897092 | 0.375 | 0.46 | 3.25038456757889e-26 |
| IFITM3 | 3.39350990686751e-30 | -0.431154962259905 | 0.541 | 0.59 | 1.35139745021185e-25 |
| CST3 | 2.32473216067505e-28 | -0.417117383085 | 0.92 | 0.92 | 9.25778088345627e-24 |
| HAMP | 2.89842719078547e-27 | -0.411944281947338 | 0.045 | 0.104 | 1.1542406601865e-22 |
| CTSC | 1.08905408578342e-26 | -0.318936608873844 | 0.305 | 0.381 | 4.33694008581532e-22 |
| TMSB4X | 7.00392565071975e-26 | -0.275666040382254 | 0.999 | 0.995 | 2.78917331188613e-21 |
| KCNMA1 | 6.16448121381825e-25 | 0.318526191136907 | 0.364 | 0.284 | 2.45488135377884e-20 |
| PSME2 | 1.02221926172604e-23 | -0.302866106412832 | 0.415 | 0.474 | 4.07078376597162e-19 |
| ARL4C | 2.24382202949684e-23 | -0.252203449906947 | 0.41 | 0.486 | 8.93557246806527e-19 |
| EGR1 | 5.08136784652401e-23 | -0.261649733952071 | 0.106 | 0.175 | 2.02355311752126e-18 |
| CALM1 | 1.05164390926865e-22 | -0.251206762645533 | 0.853 | 0.859 | 4.18796153988055e-18 |
| ITM2B | 3.07232316124266e-22 | -0.290515356695976 | 0.772 | 0.781 | 1.22349125250167e-17 |
| CTSH | 4.02995492515295e-22 | -0.262348044246808 | 0.487 | 0.535 | 1.60484894984366e-17 |
| TREM2 | 5.80399911690777e-21 | -0.373801508891415 | 0.275 | 0.329 | 2.31132656832618e-16 |
| PLTP | 8.15086992096223e-21 | -0.533145645643864 | 0.435 | 0.477 | 3.24592092862479e-16 |
| TUBA1B | 9.16999381886478e-19 | -0.401292810478764 | 0.676 | 0.697 | 3.65176663848652e-14 |
| VSIG4 | 8.38234967110564e-18 | -0.482349468487101 | 0.265 | 0.317 | 3.3381031095244e-13 |
| FOLR2 | 3.3854795996381e-17 | -0.300479503294985 | 0.196 | 0.252 | 1.34819954096388e-12 |
| CD9 | 6.97466025255707e-16 | -0.269738101547432 | 0.329 | 0.38 | 2.7775189523758e-11 |
| DNAJB1 | 2.17095265549797e-14 | -0.34979239994682 | 0.193 | 0.251 | 8.64538475998955e-10 |
| AREG | 4.49402251059876e-13 | -0.30938568647151 | 0.104 | 0.149 | 1.78965458439574e-08 |
| IL2RA | 1.0087778014128e-11 | -0.307693985174031 | 0.176 | 0.22 | 4.01725583856618e-07 |
| HSPA6 | 1.1166215067007e-09 | -0.318936057924175 | 0.068 | 0.101 | 4.4467218261342e-05 |
| DUSP1 | 3.89381568231911e-08 | -0.392899924188165 | 0.594 | 0.608 | 0.00155063421916994 |
| FCGR3A | 4.09870441373364e-08 | -0.337444361121311 | 0.365 | 0.385 | 0.00163222705868115 |
| HLA-DQA1 | 6.05870578305925e-08 | -0.333596425652635 | 0.698 | 0.747 | 0.00241275840398769 |
| ITGB2 | 1.99802443091675e-06 | -0.277220046598053 | 0.555 | 0.535 | 0.0795673269123978 |
| FCN1 | 2.10445938681274e-05 | -0.268376329229792 | 0.137 | 0.161 | 0.838058861610437 |
| CD163 | 0.00723194061466727 | -0.363714231821672 | 0.493 | 0.475 | 1 |
| ALOX5AP | 0.0911846177556649 | 0.32138295732548 | 0.541 | 0.536 | 1 |
| MARCO | 0.0949847373435471 | -0.655313310076983 | 0.352 | 0.303 | 1 |
| HSPA1A | 0.0978569734903417 | -0.475462989185813 | 0.311 | 0.308 | 1 |
| ISG15 | 0.300879334751564 | -0.522939297760506 | 0.578 | 0.544 | 1 |

Supplementary table 2: Samples collected for multiomic TME characterization

| Patient  code | Sample type | Treatment.strategy | OvaHRDscar | BRCAmut status | PFI category (12 months) | Age at diagnosis | Stage FIGO2014 | tcycif | scRNAseq | bulkRNAseq | WGS (SBS3) | GeoMx | GeoMxRoi nr (CD8-IBA1-/CD8-IBA1+/CD8+IBA1-/CD8+IBA1+) |
| --- | --- | --- | --- | --- | --- | --- | --- | --- | --- | --- | --- | --- | --- |
| S001 | IDS | NACT | HRP | BRCAwt | short | 60 | IIIC | ovary | ovary | ovary | HRP | NA |  |
| S002 | chemo-naive | NACT | HRD | BRCAmut | long | 74 | IIIC | ovary | omentum | ovary | HRD | NA |  |
| S003 | chemo-naive | PDS | HRP | BRCAwt | short | 68 | IVB | peritoneum | peritoneum | peritoneum | HRP | NA |  |
| S004 | chemo-naive | PDS | HRP | BRCAwt | long | 75 | IIIC | ovary | ovary | ovary | HRP | NA |  |
| S005 | chemo-naive | PDS | HRP | BRCAwt | short | 78 | IIIC | omentum | omentum | omentum | HRD | NA |  |
| S006 | chemo-naive | PDS | HRD | BRCAmut | long | 77 | IIIC | mesentery | mesentery | mesentery | HRD | NA |  |
| S007 | chemo-naive | PDS | HRD | NA | long | 78 | IIIC | ovary | ovary | ovary | NA | NA |  |
| S008 | IDS | NACT | HRD | BRCAwt | long | 81 | IIIC | ovary | ovary | omentum+ovary | HRP | NA |  |
| S008 | chemo-naive | NACT | HRD | NA | long | 81 | IIIC | omentum | omentum | omentum | NA | NA |  |
| S009 | IDS | NACT | HRD | BRCAwt | short | 54 | IVA | omentum | omentum | omentum | HRD | NA |  |
| S009 | chemo-naive | NACT | HRD | BRCAwt | short | 54 | IVA | omentum | omentum | omentum | HRD | NA |  |
| S010 | chemo-naive | NACT | HRD | BRCAwt | short | 62 | IIIC | peritoneum | omentum | peritoneum | HRD | NA |  |
| S011 | IDS | NACT | HRP | BRCAwt | short | 67 | IVA | omentum | omentum | omentum | HRP | NA |  |
| S011 | chemo-naive | NACT | HRP | BRCAwt | short | 67 | IVA | peritoneum | peritoneum | omentum+peritoneum | HRP | NA |  |
| S012 | IDS | NACT | HRP | BRCAwt | short | 62 | IIIC | omentum | omentum | omentum | HRP | NA |  |
| S012 | chemo-naive | NACT | HRP | BRCAwt | short | 62 | IIIC | peritoneum | peritoneum | omentum+peritoneum | HRP | NA |  |
| S013 | chemo-naive | NACT | HRP | BRCAwt | short | 72 | IVB | peritoneum | peritoneum | peritoneum | HRP | NA |  |
| S014 | IDS | NACT | HRP | NA | short | 73 | IVA | other | omentum | omentum | NA | NA |  |
| S014 | chemo-naive | NACT | HRP | BRCAwt | short | 73 | IVA | peritoneum | peritoneum | omentum+peritoneum | HRP | NA |  |
| S015 | IDS | NACT | HRD | BRCAwt | long | 78 | IVA | omentum | omentum | omentum | HRD | NA |  |
| S015 | chemo-naive | NACT | HRD | BRCAwt | long | 78 | IVA | omentum | omentum | omentum | HRD | NA |  |
| S016 | chemo-naive | NACT | HRP | BRCAwt | short | 74 | IIIC | peritoneum | peritoneum | NA | HRP | NA |  |
| S017 | IDS | NACT | NA | NA | short | 72 | IIIC | NA | omentum | omentum | NA | NA |  |
| S017 | chemo-naive | NACT | HRD | BRCAwt | short | 72 | IIIC | NA | peritoneum | omentum | HRD | NA |  |
| S018 | IDS | NACT | HRP | BRCAwt | short | 71 | IIIC | NA | omentum | omentum | HRP | NA |  |
| S018 | chemo-naive | NACT | HRP | BRCAwt | short | 71 | IIIC | NA | omentum | omentum | HRP | NA |  |
| S019 | IDS | NACT | NA | NA | short | 75 | IIIC | NA | omentum | omentum | NA | NA |  |
| S019 | chemo-naive | NACT | HRD | BRCAwt | short | 75 | IIIC | NA | omentum | omentum | HRP | NA |  |
| S020 | IDS | NACT | HRP | BRCAwt | short | 74 | IVA | NA | omentum | omentum | HRP | NA |  |
| S020 | chemo-naive | NACT | HRP | BRCAwt | short | 74 | IVA | NA | omentum | omentum | HRP | NA |  |
| S021 | IDS | NACT | NA | NA | short | 77 | IIIC | NA | omentum | omentum | NA | NA |  |
| S021 | chemo-naive | NACT | het | BRCAwt | short | 77 | IIIC | NA | omentum | omentum | HRP | NA |  |
| S022 | IDS | NACT | NA | NA | long | 77 | IVB | NA | mesentery | NA | NA | NA |  |
| S022 | chemo-naive | NACT | HRP | BRCAwt | long | 77 | IVB | NA | omentum | NA | HRP | NA |  |
| S001 | chemo-naive | NACT | HRP | BRCAwt | short | 60 | IIIC | NA | peritoneum | NA | HRP | NA |  |
| S002 | IDS | NACT | NA | BRCAwt | long | 74 | IIIC | NA | omentum | NA | HRD | NA |  |
| S025 | IDS | NACT | HRD | BRCAwt | long | 68 | IIIC | NA | peritoneum | NA | HRD | NA |  |
| S025 | chemo-naive | NACT | HRD | BRCAwt | long | 68 | IIIC | NA | peritoneum | NA | HRD | NA |  |
| S010 | IDS | NACT | HRD | gBRCAmut | short | 62 | IIIC | NA | omentum | NA | HRD | NA |  |
| S027 | IDS | NACT | NA | NA | short | 62 | IIIC | NA | omentum | NA | NA | NA |  |
| S027 | chemo-naive | NACT | HRP | BRCAwt | short | 62 | IIIC | NA | omentum | NA | HRD | NA |  |
| S028 | IDS | NACT | HRD | BRCAwt | long | 64 | IVA | NA | omentum | NA | HRD | NA |  |
| S028 | chemo-naive | NACT | HRD | BRCAwt | long | 64 | IVA | NA | mesentery | NA | HRD | NA |  |
| S029 | IDS | NACT | HRP | BRCAwt | short | 72 | IVA | NA | omentum | NA | HRD | NA |  |
| S029 | chemo-naive | NACT | HRD | BRCAwt | short | 72 | IVA | NA | peritoneum | NA | HRD | NA |  |
| S030 | IDS | NACT | NA | NA | short | 67 | IVB | NA | omentum | NA | NA | NA |  |
| S030 | chemo-naive | NACT | NA | NA | short | 67 | IVB | NA | peritoneum | NA | NA | NA |  |
| S031 | IDS | NACT | HRP | gBRCAmut | long | 78 | IIIC | NA | omentum | NA | HRD | NA |  |
| S031 | chemo-naive | NACT | HRD | gBRCAmut | long | 78 | IIIC | NA | peritoneum | NA | HRD | NA |  |
| S032 | IDS | NACT | NA | NA | short | 68 | IIIC | NA | ascites | NA | NA | NA |  |
| S032 | chemo-naive | NACT | HRP | BRCAwt | short | 68 | IIIC | NA | ascites | NA | HRP | NA |  |
| S033 | IDS | NACT | HRP | BRCAwt | short | 69 | IVA | NA | NA | omentum | HRP | NA |  |
| S033 | chemo-naive | NACT | HRP | BRCAwt | short | 69 | IVA | NA | NA | omentum | HRP | NA |  |
| S034 | IDS | NACT | HRP | BRCAwt | short | 55 | IIIC | NA | NA | omentum | HRP | NA |  |
| S034 | chemo-naive | NACT | HRP | BRCAwt | short | 55 | IIIC | NA | NA | omentum | HRP | NA |  |
| S035 | IDS | NACT | HRP | BRCAwt | short | 79 | IIIC | NA | NA | omentum | HRP | NA |  |
| S035 | chemo-naive | NACT | HRP | BRCAwt | short | 79 | IIIC | NA | NA | omentum | HRP | NA |  |
| S036 | IDS | NACT | HRP | BRCAwt | short | 76 | IIIC | NA | NA | omentum | HRP | NA |  |
| S036 | chemo-naive | NACT | HRD | BRCAwt | short | 76 | IIIC | NA | NA | omentum | HRP | NA |  |
| S037 | IDS | NACT | HRP | BRCAwt | short | 72 | IIIC | NA | NA | omentum | HRP | NA |  |
| S037 | chemo-naive | NACT | HRP | BRCAwt | short | 72 | IIIC | NA | NA | omentum | HRP | NA |  |
| S038 | IDS | NACT | HRD | BRCAwt | long | 71 | IIIC | NA | NA | omentum | HRD | NA |  |
| S038 | chemo-naive | NACT | HRD | BRCAwt | long | 71 | IIIC | NA | NA | omentum | HRD | NA |  |
| S039 | IDS | NACT | HRP | BRCAwt | short | 68 | IIIC | NA | NA | omentum | HRP | NA |  |
| S039 | chemo-naive | NACT | HRP | BRCAwt | short | 68 | IIIC | NA | NA | omentum | HRP | NA |  |
| S040 | IDS | NACT | HRD | BRCAwt | long | 64 | IVB | NA | NA | omentum | HRD | NA |  |
| S040 | chemo-naive | NACT | HRD | BRCAwt | long | 64 | IVB | NA | NA | omentum | HRD | NA |  |
| S041 | IDS | NACT | HRP | BRCAwt | short | 70 | IVA | NA | NA | omentum | HRP | NA |  |
| S041 | chemo-naive | NACT | HRP | BRCAwt | short | 70 | IVA | NA | NA | omentum | HRP | NA |  |
| S042 | IDS | NACT | HRP | BRCAwt | short | 68 | IIIC | NA | NA | omentum | HRP | NA |  |
| S042 | chemo-naive | NACT | HRP | BRCAwt | short | 68 | IIIC | NA | NA | omentum | HRP | NA |  |
| S043 | IDS | NACT | HRP | BRCAwt | short | 73 | IIIC | NA | NA | omentum | HRP | NA |  |
| S043 | chemo-naive | NACT | HRP | BRCAwt | short | 73 | IIIC | NA | NA | omentum | HRP | NA |  |
| S044 | IDS | NACT | HRD | BRCAwt | short | 75 | IIIC | NA | NA | omentum | HRD | NA |  |
| S044 | chemo-naive | NACT | het | BRCAwt | short | 75 | IIIC | NA | NA | omentum | HRD | NA |  |
| S045 | IDS | NACT | HRP | BRCAwt | short | 70 | IVA | NA | NA | omentum | HRP | NA |  |
| S045 | chemo-naive | NACT | HRP | BRCAwt | short | 70 | IVA | NA | NA | omentum | HRP | NA |  |
| S046 | IDS | NACT | NA | NA | short | 57 | IIIC | NA | NA | omentum | NA | NA |  |
| S046 | chemo-naive | NACT | NA | NA | short | 57 | IIIC | NA | NA | omentum | NA | NA |  |
| S047 | IDS | NACT | NA | NA | short | 63 | IIIC | NA | NA | omentum | NA | NA |  |
| S047 | chemo-naive | NACT | NA | NA | short | 63 | IIIC | NA | NA | omentum | NA | NA |  |
| C063 | IDS | NACT | HRP | BRCAwt | long | 66 | IIIC | omentum | NA | omentum | NA | omentum | 1/1/1/7 |
| C063 | chemo-naive | NACT | HRP | BRCAwt | long | 66 | IIIC | peritoneum | NA | NA | NA | peritoneum | 1/2/1/5 |
| C033 | IDS | NACT | HRP | BRCAwt | long | 63 | IVB | omentum | NA | omentum | NA | omentum | 1/1/1/7 |
| C033 | chemo-naive | NACT | HRP | BRCAwt | long | 63 | IVB | omentum | NA | NA | NA | omentum | 1/1/1/8 |
| C686 | IDS | NACT | HRD | BRCAwt | long | 58 | IVB | ovary | NA | ovary | NA | ovary | 1/1/1/7 |
| C686 | chemo-naive | NACT | HRD | BRCAwt | long | 58 | IVB | omentum | NA | NA | NA | omentum | 1/1/1/7 |
| C879 | IDS | NACT | HRP | BRCAwt | short | 74 | IIIC | omentum | NA | NA | NA | omentum | 0/1/2/8 |
| C538 | IDS | NACT | HRP | BRCAwt | long | 55 | IVB | omentum | NA | omentum | NA | omentum | 1/1/1/7 |
| C799 | IDS | NACT | HRP | BRCAwt | long | 67 | IIIC | omentum | NA | omentum | NA | omentum | 1/1/1/7 |
| C917 | IDS | NACT | HRP | BRCAwt | short | 73 | IVB | omentum | NA | omentum | NA | omentum | 1/1/1/7 |
| C790 | IDS | NACT | HRP | BRCAwt | short | 69 | IVB | omentum | NA | omentum | NA | omentum | 1/1/1/7 |
| C423 | IDS | NACT | HRP | BRCAwt | long | 63 | IVB | omentum | NA | NA | NA | omentum | 1/1/1/7 |
| C129 | IDS | NACT | HRP | BRCAwt | short | 60 | IVB | omentum | NA | omentum | NA | omentum | 1/1/1/6 |
| C129 | chemo-naive | NACT | HRP | BRCAwt | short | 60 | IVB | adnex | NA | NA | NA | adnex | 1/1/1/7 |
| C535 | IDS | NACT | HRP | BRCAwt | long | 64 | IIIC | omentum | NA | omentum | NA | omentum | 1/1/1/9 |
| C535 | chemo-naive | NACT | HRP | BRCAwt | long | 64 | IIIC | peritoneum | NA | NA | NA | peritoneum | 1/1/1/5 |

Supplementary table 3. Pathways used in GeoMx analysis

| HALLMARK_APOPTOSIS | ADD1 AIFM3 ANKH ANXA1 APP ATF3 AVPR1A BAX BCAP31 BCL10 BCL2L1 BCL2L10 BCL2L11 BCL2L2 BGN BID BIK BIRC3 BMF BMP2 BNIP3L BRCA1 BTG2 BTG3 CASP1 CASP2 CASP3 CASP4 CASP6 CASP7 CASP8 CASP9 CAV1 CCNA1 CCND1 CCND2 CD14 CD2 CD38 CD44 CD69 CDC25B CDK2 CDKN1A CDKN1B CFLAR CLU CREBBP CTH CTNNB1 CYLD DAP DAP3 DCN DDIT3 DFFA DIABLO DNAJA1 DNAJC3 DNM1L DPYD EBP EGR3 EMP1 ENO2 ERBB2 ERBB3 EREG ETF1 F2 F2R FAS FASLG FDXR FEZ1 GADD45A GADD45B GCH1 GNA15 GPX1 GPX3 GPX4 GSN GSR GSTM1 GUCY2D H1-0 HGF HMGB2 HMOX1 HSPB1 IER3 IFITM3 IFNB1 IFNGR1 IGF2R IGFBP6 IL18 IL1A IL1B IL6 IRF1 ISG20 JUN KRT18 LEF1 LGALS3 LMNA PLPPR4 LUM MADD MCL1 MGMT MMP2 NEDD9 NEFH PAK1 PDCD4 PDGFRB PEA15 PLAT PLCB2 PMAIP1 PPP2R5B PPP3R1 PPT1 PRF1 PSEN1 PSEN2 PTK2 RARA RELA RETSAT RHOB RHOT2 RNASEL ROCK1 SAT1 SATB1 SC5D SLC20A1 SMAD7 SOD1 SOD2 SPTAN1 SQSTM1 TAP1 TGFB2 TGFBR3 TIMP1 TIMP2 TIMP3 TNF TNFRSF12A TNFSF10 TOP2A TSPO TXNIP VDAC2 WEE1 XIAP |
| --- | --- |
| HALLMARK_HYPOXIA | ADM ADORA2B AK4 AKAP12 ALDOA ALDOB ALDOC AMPD3 ANGPTL4 ANKZF1 ANXA2 ATF3 ATP7A B3GALT6 B4GALNT2 BCAN BCL2 BGN BHLHE40 BNIP3L BRS3 BTG1 CA12 CASP6 CAV1 CCNG2 NOCT CDKN1A CDKN1B CDKN1C CHST2 CHST3 CITED2 COL5A1 CP CSRP2 CCN2 CXCR4 ACKR3 CCN1 DCN DDIT3 DDIT4 DPYSL4 DTNA DUSP1 EDN2 EFNA1 EFNA3 EGFR ENO1 ENO2 ENO3 ERO1A ERRFI1 ETS1 EXT1 F3 FAM162A FBP1 FOS FOSL2 FOXO3 GAA GALK1 GAPDH GAPDHS GBE1 GCK GCNT2 GLRX GPC1 GPC3 GPC4 GPI GRHPR GYS1 HAS1 HDLBP HEXA HK1 HK2 HMOX1 HOXB9 HS3ST1 HSPA5 IDS IER3 IGFBP1 IGFBP3 IL6 ILVBL INHA IRS2 ISG20 JMJD6 JUN KDELR3 KDM3A KIF5A KLF6 KLF7 KLHL24 LALBA LARGE1 LDHA LDHC LOX LXN MAFF MAP3K1 MIF MT1E MT2A MXI1 MYH9 NAGK NCAN NDRG1 NDST1 NDST2 NEDD4L NFIL3 NR3C1 P4HA1 P4HA2 PAM PCK1 PDGFB PDK1 PDK3 PFKFB3 PFKL PFKP PGAM2 PGF PGK1 PGM1 PGM2 PHKG1 PIM1 PKLR PKP1 PLAC8 PLAUR PLIN2 PNRC1 PPARGC1A PPFIA4 PPP1R15A PPP1R3C PRDX5 PRKCA CAVIN3 CAVIN1 PYGM RBPJ RORA RRAGD S100A4 SAP30 SCARB1 SDC2 SDC3 SDC4 SELENBP1 SERPINE1 SIAH2 SLC25A1 SLC2A1 SLC2A3 SLC2A5 SLC37A4 SLC6A6 SRPX STBD1 STC1 STC2 SULT2B1 TES TGFB3 TGFBI TGM2 TIPARP TKTL1 TMEM45A TNFAIP3 TPBG TPD52 TPI1 TPST2 UGP2 VEGFA VHL VLDLR CCN5 WSB1 XPNPEP1 ZFP36 ZNF292 |
| HALLMARK_IL2_STAT5_SIGNALING | ABCB1 ADAM19 AGER AHCY AHNAK AHR ALCAM AMACR ANXA4 APLP1 ARL4A BATF BATF3 BCL2 BCL2L1 BHLHE40 BMP2 BMPR2 CA2 CAPG CAPN3 CASP3 DRC1 CCND2 CCND3 CCNE1 CCR4 CD44 CD48 CD79B CD81 CD83 CD86 CDC42SE2 CDC6 CDCP1 CDKN1C CISH CKAP4 COCH COL6A1 CSF1 CSF2 CST7 CTLA4 CTSZ CXCL10 CYFIP1 DCPS DENND5A DHRS3 ECM1 EMP1 ENO3 ENPP1 EOMES ETV4 F2RL2 FAH HYCC2 FGL2 FLT3LG FURIN GABARAPL1 GADD45B GALM GATA1 GBP4 GLIPR2 GPR65 GPR83 GPX4 GSTO1 GUCY1B1 HIPK2 HK2 HOPX HUWE1 ICOS IFITM3 IFNGR1 IGF1R IGF2R IKZF2 IKZF4 IL10 IL10RA IL13 IL18R1 IL1R2 IL1RL1 IL2RA IL2RB IL3RA IL4R IRF4 IRF6 IRF8 ITGA6 ITGAE ITGAV ITIH5 KLF6 LCLAT1 LIF LRIG1 LRRC8C LTB MAFF MAP3K8 MAP6 MAPKAPK2 ETFBKMT MUC1 MXD1 MYC MYO1C MYO1E EEF1AKMT1 NCOA3 NCS1 NDRG1 NFIL3 NFKBIZ NOP2 NRP1 NT5E ODC1 P2RX4 P4HA1 PDCD2L PENK PHLDA1 PHTF2 PIM1 PLAGL1 PLEC PLIN2 PLSCR1 PNP POU2F1 PLPP1 PRAF2 PRKCH PRNP PTCH1 PTGER2 PTH1R PTRH2 PUS1 RABGAP1L RGS16 RHOB RHOH RNH1 RORA RRAGD S100A1 SCN9A SELL SELP SERPINB6 SERPINC1 SH3BGRL2 SHE SLC1A5 SLC29A2 SLC2A3 SLC39A8 SMPDL3A SNX14 SNX9 SOCS1 SOCS2 SPP1 SPRED2 SPRY4 ST3GAL4 SWAP70 SYNGR2 SYT11 TGM2 TIAM1 TLR7 TNFRSF18 TNFRSF1B TNFRSF21 TNFRSF4 TNFRSF8 TNFRSF9 TNFSF10 TNFSF11 TRAF1 TTC39B TWSG1 UCK2 UMPS WLS XBP1 |
| HALLMARK_INFLAMMATORY_RESPONSE | ABCA1 ABI1 ACVR1B ACVR2A ADM ADORA2B ADRM1 AHR APLNR AQP9 ATP2A2 ATP2B1 ATP2C1 AXL BDKRB1 BEST1 BST2 BTG2 C3AR1 C5AR1 CALCRL CCL17 CCL2 CCL20 CCL22 CCL24 CCL5 CCL7 CCR7 CCRL2 CD14 CD40 CD48 CD55 CD69 CD70 CD82 CDKN1A CHST2 CLEC5A CMKLR1 CSF1 CSF3 CSF3R CX3CL1 CXCL10 CXCL11 CXCL6 CXCL9 CXCR6 CYBB DCBLD2 EBI3 EDN1 EIF2AK2 EMP3 ADGRE1 EREG F3 FFAR2 FPR1 FZD5 GABBR1 GCH1 GNA15 GNAI3 GP1BA GPC3 GPR132 GPR183 HAS2 HBEGF HIF1A HPN HRH1 ICAM1 ICAM4 ICOSLG IFITM1 IFNAR1 IFNGR2 IL10 IL10RA IL12B IL15 IL15RA IL18 IL18R1 IL18RAP IL1A IL1B IL1R1 IL2RB IL4R IL6 IL7R CXCL8 INHBA IRAK2 IRF1 IRF7 ITGA5 ITGB3 ITGB8 KCNA3 KCNJ2 KCNMB2 KIF1B KLF6 LAMP3 LCK LCP2 LDLR LIF LPAR1 LTA LY6E LYN MARCO MEFV MEP1A MET MMP14 MSR1 MXD1 MYC NAMPT NDP NFKB1 NFKBIA NLRP3 NMI NMUR1 NOD2 NPFFR2 OLR1 OPRK1 OSM OSMR P2RX4 P2RX7 P2RY2 PCDH7 PDE4B PDPN PIK3R5 PLAUR PROK2 PSEN1 PTAFR PTGER2 PTGER4 PTGIR PTPRE PVR RAF1 RASGRP1 RELA RGS1 RGS16 RHOG RIPK2 RNF144B ROS1 RTP4 SCARF1 SCN1B SELE SELL SELENOS SEMA4D SERPINE1 SGMS2 SLAMF1 SLC11A2 SLC1A2 SLC28A2 SLC31A1 SLC31A2 SLC4A4 SLC7A1 SLC7A2 SPHK1 SRI STAB1 TACR1 TACR3 TAPBP TIMP1 TLR1 TLR2 TLR3 TNFAIP6 TNFRSF1B TNFRSF9 TNFSF10 TNFSF15 TNFSF9 TPBG VIP |
| HALLMARK_TNFA_SIGNALING_VIA_NFKB | ABCA1 AREG ATF3 ATP2B1 B4GALT1 B4GALT5 BCL2A1 BCL3 BCL6 BHLHE40 BIRC2 BIRC3 BMP2 BTG1 BTG2 BTG3 CCL2 CCL20 CCL4 CCL5 CCND1 CCNL1 CCRL2 CD44 CD69 CD80 CD83 CDKN1A CEBPB CEBPD CFLAR CLCF1 CSF1 CSF2 CXCL1 CXCL10 CXCL11 CXCL2 CXCL3 CXCL6 ACKR3 CCN1 RIGI DENND5A DNAJB4 DRAM1 DUSP1 DUSP2 DUSP4 DUSP5 EDN1 EFNA1 EGR1 EGR2 EGR3 EHD1 EIF1 ETS2 F2RL1 F3 FJX1 FOS FOSB FOSL1 FOSL2 FUT4 G0S2 GADD45A GADD45B GCH1 GEM GFPT2 GPR183 HBEGF HES1 ICAM1 ICOSLG ID2 IER2 IER3 IER5 IFIH1 IFIT2 IFNGR2 IL12B IL15RA IL18 IL1A IL1B IL23A IL6 IL6ST IL7R INHBA IRF1 IRS2 JAG1 JUN JUNB KDM6B KLF10 KLF2 KLF4 KLF6 KLF9 KYNU LAMB3 LDLR LIF LITAF MAFF MAP2K3 MAP3K8 MARCKS MCL1 MSC MXD1 MYC NAMPT NFAT5 NFE2L2 NFIL3 NFKB1 NFKB2 NFKBIA NFKBIE NINJ1 NR4A1 NR4A2 NR4A3 OLR1 PANX1 PDE4B PDLIM5 PER1 PFKFB3 PHLDA1 PHLDA2 PLAU PLAUR PLEK PLK2 PMEPA1 PNRC1 PLPP3 PPP1R15A PTGER4 PTGS2 PTPRE PTX3 RCAN1 REL RELA RELB RHOB RIPK2 RNF19B SAT1 SDC4 SERPINB2 SERPINB8 SERPINE1 SGK1 SIK1 SLC16A6 SLC2A3 SLC2A6 SMAD3 SNN SOCS3 SOD2 SPHK1 SPSB1 SQSTM1 STAT5A TANK TAP1 TGIF1 TIPARP TLR2 TNC TNF TNFAIP2 TNFAIP3 TNFAIP6 TNFAIP8 TNFRSF9 TNFSF9 TNIP1 TNIP2 TRAF1 TRIB1 TRIP10 TSC22D1 TUBB2A VEGFA YRDC ZBTB10 ZC3H12A ZFP36 |
| BIOCARTA_CTLA4_PATHWAY | PIK3R1 PIK3CA CD3G IL2 CD3D CD3E CD247 CTLA4 LCK CD86 CD28 HLA-DRB1 GRB2 HLA-DRB5 PTPN11 CD80 ITK ICOS ICOSLG HLA-DRA HLA-DRB4 HLA-DRB3 |
| REACTOME_CTLA4_INHIBITORY_SIGNALING | FYN PPP2R5A PPP2R5B PPP2R5C PPP2CB AKT2 PPP2R1A PPP2R5D PPP2CA CD86 AKT3 CD80 PPP2R1B AKT1 PPP2R5E CTLA4 YES1 PTPN11 LCK SRC LYN |
| GOBP_REGULATION_OF_T_CELL_APOPTOTIC_PROCESS | ADA BCL2L11 ADAM8 RIPK3 LGALS16 GIMAP8 PTCRA TSC22D3 EFNA1 KIFAP3 PRKD2 PRELID1 CD274 HIF1A IL7R IDO1 JAK3 ARG2 LGALS3 LGALS9 P2RX7 PDCD1 PIP PRKCQ GPAM SLC46A2 RAG1 BCL2 BCL3 CCL5 ST3GAL1 BCL11B BMP4 TP53 WNT5A DOCK8 ZC3H8 FADD CD27 |
| KEGG_JAK_STAT_SIGNALING_PATHWAY | AKT3 SPRY3 SPRY1 SPRY2 STAM2 IRF9 PIAS3 IL24 CISH IL22RA2 SOCS4 CNTF CNTFR CREBBP CSF2 CSF2RA CSF2RB CSF3 CSF3R CSH1 CTF1 IL23R SPRED1 IFNLR1 SPRED2 EP300 EPO EPOR AKT1 AKT2 CLCF1 PIK3R5 CBLC GH1 GH2 GHR IFNL2 IFNL3 IFNL1 GRB2 IL19 SOCS7 IFNE IFNA1 IFNA2 IFNA4 IFNA5 IFNA6 IFNA7 IFNA8 IFNA10 IFNA13 IFNA14 IFNA16 IFNA17 IFNA21 IFNAR1 IFNAR2 IFNB1 IFNG IFNGR1 IFNGR2 IFNW1 IL2 IL2RA IL2RB IL2RG IL3 IL3RA IL4 IL4R IL5 IL5RA IL6 IL6R IL6ST IL7 IL7R IL9 IL9R IL10 IL10RA IL10RB IL11 IL11RA IL12A IL12B IL12RB1 IL12RB2 IL13 IL13RA1 IL13RA2 IL15 IL15RA JAK1 JAK2 JAK3 LEP LEPR LIF LIFR MPL MYC OSM IL20 IL21R IL22 IL23A PIAS4 PIK3CA PIK3CB PIM1 PIK3CD PIK3CG PIK3R1 PIK3R2 IL20RA IL20RB IL26 PRL PRLR IFNK PTPN6 PTPN11 IL22RA1 IL21 CCND1 BCL2L1 CRLF2 SOS1 SOS2 STAT1 STAT2 STAT3 STAT4 STAT5A STAT5B STAT6 TPO TYK2 STAM SPRY4 PIK3R3 TSLP PIAS1 SOCS1 CBL CBLB SOCS2 CCND2 CCND3 SOCS3 PIAS2 OSMR SOCS5 |
| KEGG_MTOR_SIGNALING_PATHWAY | AKT3 EIF4B EIF4E EIF4EBP1 AKT1 AKT2 VEGFD PIK3R5 MTOR RICTOR EIF4E1B ULK3 RPS6KA6 HIF1A IGF1 INS PDPK1 CAB39 PGF PIK3CA PIK3CB PIK3CD PIK3CG PIK3R1 PIK3R2 DDIT4 PRKAA1 PRKAA2 MAPK1 MAPK3 RPTOR RHEB RPS6 RPS6KA1 RPS6KA2 RPS6KA3 RPS6KB1 RPS6KB2 MLST8 BRAF STK11 TSC1 TSC2 VEGFA VEGFB VEGFC CAB39L ULK1 PIK3R3 STRADA EIF4E2 ULK2 |
| BIOCARTA_IL2_PATHWAY | SOS1 IL2RG JAK3 IL2RA IL2 IL2RB MAPK3 LCK ELK1 SHC1 HRAS SYK CSNK2A1 GRB2 JAK1 JUN MAPK8 MAP2K1 RAF1 STAT5A FOS STAT5B |
| WP_MACROPHAGE_MARKERS | F3 LYZ RAC2 CD14 CD83 CD163 CD86 CD68 CD74 |
| COATES_MACROPHAGE_M1_VS_M2_DN | ADAM8 AK4 ALDOC ANG ARG1 BNIP3 BST1 C1QB C1QC CBLB CD5L CTSK F7 F10 HLA-A HLA-B HSPA1A HSPA1B IGF2R ITGB3 LBP MFGE8 MGST2 MYO1F PTGER2 SCD SLC2A1 SOAT1 PRDX2 THBS1 TLR1 UCHL1 VEGFA XDH HS6ST1 TJP2 CRIPT GDF15 CYTIP IL18BP PROCR SH2B2 CD300C RRAS2 PDXDC1 RIMBP2 PADI4 FLRT2 CHIA RGCC HILPDA ERO1A GASK1B GPRC5B FBLIM1 QRICH1 MCOLN3 LIN7C TULP4 RPGRIP1 COL20A1 TMEM267 PINK1 EFHD2 LRRC27 AKR1E2 ATG4D ZNF616 PKDCC EGLN3 AHNAK2 FAM241A FAM199X ARSK ATP6V0D2 FAM177A1 ANKRD37 C5orf34 C1QTNF12 CD300LD CD24 |
| COATES_MACROPHAGE_M1_VS_M2_UP | ACYP2 ADA AIF1 ALAD ARSB AXL BCKDHB DST CAPN5 CD59 RCBTB2 COMT ENO2 GATM GJA1 GLO1 CFH HLA-DQA1 HLA-DQA2 KCNJ10 LAMA3 MAF MSR1 NRCAM PPP2R5C PTGER3 RAPSN RCN1 RFX3 DYNLT1 TGFBI TTC3 ZNF85 CXCR4 MARCO RPL14 SLC7A7 HMGN3 RASAL2 CXCL14 GDA SFI1 ABCA9 CAP1 SLC35A1 TXNIP RAB10 PLAAT3 FILIP1L SLCO2B1 EMC1 CBX6 CADM1 CNRIP1 BLOC1S6 CRIM1 TM7SF3 P2RY13 HAUS2 ARFGEF3 FAM135A GNB4 GALNT11 TOR3A MARCKSL1 OGFRL1 OCEL1 SRD5A3 PLBD1 COLEC12 ACSS1 ABHD1 PLXDC2 ZNF616 PIK3AP1 CLBA1 LYPLAL1 ATP6V0E2 PIANP FAM124A FGD2 LCORL NAT8L TUBB2B HDDC3 TIFAB |
| GSE5099_CLASSICAL_M1_VS_ALTERNATIVE_M2_MACROPHAGE_DN | AEBP1 ALDH2 STS ADGRB2 DST CD1D CD79B CDC25A CDK6 CDSN LYST CHRNA3 CHRNA4 COL6A3 COL7A1 CYP2F1 DBN1 DDX5 DLAT TRDMT1 ECT2 ELAVL4 CTTN EPB41L2 ETV5 GALNT2 GLI4 GNA15 GNAQ GPR4 H3-3B HMGB3 HNRNPA2B1 HNRNPK HOXA7 HOXA9 HOXA10 HTR2A IL4 INHBC INPP5B ITIH3 ANOS1 KIR2DL4 LMO2 SH2D1A MDM4 MELTF KMT2A NEK2 SLC22A18AS PCMT1 POU3F4 PPEF1 PKN2 PRKY PTGS1 RPS4Y1 SORT1 XCL1 SOX4 SP4 ITPRID2 TCF7 TFF3 TFPI SEM1 CUBN ANP32A KDM5D USP9Y FZD1 PPM1D GAS7 AKR1C3 DDX3Y EIF1AY KIF23 ADAMTS1 KIF20B CCDC144A LAPTM4A SPCS2 EPM2AIP1 GJC1 CORIN PDE10A SLC22A7 YWHAQ PTGDR2 CD300A CHSY1 R3HDM2 PLCH1 PALLD DNAJC9 PHLPP1 TSPYL4 CBX5 SLC7A11 GGA1 OR2F1 LINC01558 SIGLEC9 TNFRSF21 IFT81 ZBTB44 RAPGEFL1 POMP UBE2J1 GALNT7 CCNJ FAM118A NEIL3 BATF3 SMPD3 RUFY2 KIAA1217 CMC2 RGMA RTN4 PELI2 LRRN1 OPRPN HPSE2 HHIP BCL11B MARCKSL1 CEP97 QSER1 RUBCNL CHD9 PCGF5 SLC7A3 RNFT2 UBASH3B CFAP74 DEPDC7 SYCE1 ZNF257 MBD6 GPRIN1 OSBPL5 MSI2 DNAI3 EVC2 CAPSL GOT1L1 WFDC3 LDLRAD3 LDC1P GARIN3 LRRN4 ZNF169 SPRED2 CENPV SLC44A5 CCDC83 ZBTB12 PHACTR1 TTTY10 TXLNGY CALHM5 PDZD9 GLIPR1L1 LINC01686 SEC14L4 H2BW2 FBLL1 LILRA5 ANKRD20A11P KRTAP5-2 TSPAN11 LINC00205 ANKRD36BP2 GXYLT2 ZNF736 ZSCAN16-AS1 TIMM23B CRNDE |
| BIOCARTA_MHC_PATHWAY | ACAA1 ADD3 BIN1 ANK3 ANXA6 APRT BTG1 CAPN2 CBLB CD27 CD48 CCR7 COL1A2 COX7C CTSL CTSZ CD55 E4F1 S1PR1 EEF1G EEF2 EPB41 FOXO1 FLNA FXN FUCA1 FYB1 GGT1 GYPC HLA-B HLA-G IL4R IL12RB1 LFNG LIPA LTA MAF MFNG MGAT1 MLLT3 NELL2 NFKBIA PNP NPAS2 P4HA1 PDE4B CFP PFDN5 PSME2 RGS1 RGS16 RLN1 RPL6 RPL7A RPL9 RPL12 RPL13 RPL17 RPL30 RPL29 RPL31 RPL32 RPLP1 RPLP2 RPS6KA3 RPS10 RPS15A RPS18 RPS19 S100A6 S100B TPP2 TUBA4A TXK UPP1 XIST ZNF16 ZNF229 EIF3F INPP4B NOG TSIX IL27RA EIF4E2 LITAF BAG3 LPIN2 PRORP KIAA0513 RASSF2 BMS1 ELMO1 AREL1 TLK1 GPA33 RGS19 SEMA3A BTN3A3 CEPT1 SPON1 CIB1 ARID5A HCP5 AP3M2 ADAMTS6 RASA3 LPIN1 ESYT1 DOCK9 COTL1 RPL13A ARHGAP45 SAMHD1 LRIG1 AUTS2 LDLRAP1 TES GIMAP2 GNL3 PABPC1 LAMP3 FOXP1 EIF3K NKIRAS1 CNIH4 C11orf21 HCFC2 GMPPA CERS2 NOP53 HEBP1 PLEKHO1 PLAC8 DPH5 ZFAND6 LRRN3 DEF8 SEMA4C TRMT10C NMRK1 FBXO34 LRRC8D PANK4 IMPACT CNOT11 POLR3E PAG1 ZNF302 PARP11 ZNFX1 ATP10A ATP8B2 STIM2 RNF213 SFMBT2 SQOR PLEKHA2 IFIH1 ERAP2 IKZF4 SMURF2 YTHDC2 PGAP1 PUS1 APOL3 KDM7A TSPAN14 SLA2 LRRC8C KISS1R TMEM128 RSAD2 EPSTI1 ERMAP GBP4 C17orf49 FBLN7 CD200R1 TRIM69 RIMS4 MPP7 PYHIN1 GPR155 GIMAP8 ZNF883 SPIN3 MPZL3 SUSD3 SPATA13 UBALD2 SMIM20 TOMM5 |
| REACTOME_CLASS_I_MHC_MEDIATED_ANTIGEN_PROCESSING_PRESENTATION | KLHL13 CDC27 FBXL3 ASB4 MYLIP PSMB1 UBE3C BTK MRC2 PSMC4 RNF14 UFL1 UBR2 UBA6 RNF19A CUL3 PSMA4 CUL7 HSPA5 VAMP3 NEDD4L CYBA ANAPC4 CUL1 SEC61A1 UNKL RNF4 BTBD1 SEC61A2 ASB1 PSME4 NEDD4 RNF126 LNX1 UBE2D1 FBXW11 KLHL20 UBE2A UBE2K PIK3C3 ITCH UBE2D4 KEAP1 UBA5 ITGB5 HACE1 HUWE1 PSMC5 KLHL42 ANAPC5 PSME1 HECTD1 SNAP23 CDC23 PSMD5 PSMD8 FBXL19 CDC34 KLHL22 FBXO7 NCF4 RBX1 TRIM9 PSMC6 PSMA3 ASB2 PSMC1 PSMB5 PSMA6 PSME2 SEC23A PSMA7 PSMD10 ASB9 MGRN1 PSMD7 FBXO31 STUB1 ELOB STX4 HERC1 UBE2W IKBKB FZR1 PSMA2 SEC61B UBE2R2 FBXW4 FBXL15 CUL2 UBE2S FBXL20 PSMD3 TRIM37 BLMH PSMD11 SMURF2 WSB1 UBE2D3 KLHL2 FBXW7 KLHL5 ZBTB16 UBE4A PSMD9 SPSB2 FBXO9 FBXL4 TAPBP RNF130 LNPEP SKP1 SEC24A UBE3A RNF7 CBLB PSMD14 ASB3 CD207 RNF19B FBXO2 FBXO6 NCF2 CDC20 UBE3D FBXO30 FBXL5 VAMP8 UBE2B TRIM32 FBXW2 AREL1 KBTBD7 RBBP6 WWP1 RNF114 PSMF1 RBCK1 PSMB HECTD3 BECN1 ATG14 CANX FBXL12 UBR4 FBXL16 RNF6 SEM1 HERC2 ANAPC13 PSMA1 CDC16 UBAC1 UBE2M UBA1 RLIM PSME3 UBE2D2 TRAF7 FBXW9 TRIM21 LRRC41 UBE2G1 SEC61G FBXO44 MKRN1 VHL RNF138 TPP2 CTSL FBXO21 CD36 FBXL8 LMO7 TLR4 PSMB7 CTSV RNF144B HLA-F TLR2 FBXO11 HECW2 ITGAV HERC3 HERC6 HERC5 SEC31A SEC24B UBE2Q2 DET1 NPEPPS ANAPC11 FBXO15 PSMB6 PSMA5 S100A8 RPS27A UBA3 DCAF1 SKP2 KLHL3 TRIM41 RNF217 TRIM50 ASB15 TRIM4 ASB10 FBXO10 ASB6 LRSAM1 HERC4 FCGR1A TIRAP SEC24D UBC UBE3B FBXO4 TRIM36 SAR1B ANAPC1 TRIP12 ASB17 LONRF1 TRIM11 SH3RF1 ELOC LY96 PSMA8 FBXL18 UBE2L6 FBXO32 SEC13 RNF111 TRIM63 NCF1 FBXW5 UBE2Z PSMD4 PSMB4 UBR1 UBE2J2 S100A1 UBE2Q1 FBXL13 PSMC2 FBXO27 ASB16 CCNF KLHL21 FBXO41 CTSS S100A9 KBTBD8 RNF25 PSMD6 RCHY1 FBXO40 DTX3L FBXW12 RNF123 ASB5 ANAPC10 ERAP1 ERAP2 CYBB ASB11 HECTD2 LRR1 KBTBD6 PSMC3 BTRC CUL5 B2M PDIA3 FBXO22 KCTD6 TAP1 UBE2V2 UBE2E3 UBE2E1 UBB CD14 RNF34 FGG FGA FGB SPSB1 FBXL14 FBXW10 THOP1 MYD88 PSMD1 RNF213 TLR1 TLR6 GLMN FBXW8 UBE2C SPSB4 PSMD2 CBLL2 UBE2O ANAPC2 CDC26 MEX3C SEC24C UBE2U ARIH2 UBE2N ASB8 KLHL11 CALR ZNRF2 RNF182 PJA1 SIAH2 RNF41 ASB18 UBA7 UBE2E2 ASB7 FBXL7 KLHL25 TRAIP UBE2F SOCS3 UBE2G2 BTBD6 UBOX5 SOCS1 PRKN PSMD13 UBE2L3 BCAP31 TRIM69 ZNRF1 UBE2H RNF220 HMGB1 ASB13 PIK3R4 SIAH1 ANAPC7 PSMD12 FBXL22 MIB2 ATG7 FCGR1BP KLHL9 SMURF1 UBE2J1 LTN1 ASB12 DZIP3 PJA2 PSMB8 TAP2 HLA-C HLA-E TRIM39 HLA-G PSMB10 HLA-B HLA-A TRIM71 CHUK UBA52 PSMB11 KBTBD13 ASB14 KLHL41 PSMB9 KCTD7 UBE2V1 MRC1 GAN RNF115 SEC22B FBXO17 IKBKG PSMB3 |
| REACTOME_MHC_CLASS_II_ANTIGEN_PRESENTATION | AP2B1 CD74 AP2S1 CTSA KIF26A KIF2A AP1M1 RAB7A TUBA3D KIFAP3 DYNC1I2 CAPZB KIF22 DNM2 KIF3C DYNLL1 LAG3 KIF4A AP1B1 LGMN SEC23A TUBB1 KIF3B CTSH DCTN6 TUBB4A KLC3 AP1S1 DNM1 SH3GL2 CTSC KIF20A SEC24A ACTR1B CAPZA1 CTSD KIF18A CLTA TUBA1B KLC1 CANX TUBA4A AP1M2 KIF3A ACTR10 DCTN4 CTSL DYNC1LI2 CTSV DCTN3 KLC4 TUBB2A TUBB2B KIF23 ACTR1A KIF11 SEC31A CENPE SEC24B KIF2B CLTC OSBPL1A KIF2C CTSK ARF1 DYNC1LI1 SEC24D AP1S3 TUBA3E SAR1B KIF5A SEC13 DYNC1I1 AP2M1 RACGAP1 CTSS KIF15 CTSB AP1G1 DCTN5 TUBA1A TUBA1C RILP HLA-DPA1 KIF5B TUBB8B SPTBN2 CTSF KLC2 DCTN2 TUBB6 SEC24C CAPZA3 TUBAL3 AP1S2 AP2A2 TUBA8 TUBB4B HLA-DRB3 HLA-DRB1 CTSE HLA-DQB2 HLA-DQA1 HLA-DQA2 AP2A1 DYNC1H1 DNM3 TUBA3C HLA-DRB5 CAPZA2 HLA-DOA HLA-DMA HLA-DRA DCTN1 HLA-DQA2 HLA-DPB1 IFI30 HLA-DQB1 HLA-DMB KIF4B HLA-DRB4 HLA-DOB TUBA4B CTSO TUBB3 TUBB8 DYNLL2 |
| REACTOME_PD_1_SIGNALING | CD4 CSK PTPN6 CD274 CD3G CD3D PTPN11 LCK PDCD1 PDCD1LG2 HLA-DRB5 CD247 CD3E HLA-DRB4 HLA-DRB3 HLA-DQA1 HLA-DQB2 HLA-DQA2 HLA-DQB1 HLA-DRA HLA-DPA1 HLA-DPB1 HLA-DQA2 HLA-DPB1 HLA-DQA2 PDCD1 HLA-DRA |
| TCELL_EXHAUSTION_ZHANG | PDCD1 TOX CXCL13 TIGIT CTLA4 TNFRSF9 HAVCR2 LAG3 CST7 GZMK GZMA NKG7 IFNG PRF1 GZMB GNLY CD28 EOMES CCR5 CCL4 CCL5 CD200R1 TMEM155 CD27 TOX2 TSHZ2 BATF GEM CD200 SNX9 ENTPD1 LAYN ETV1 PRDM1 CSF1 MYO7A MYO1E GNG4 BTLA FOXP3 TNFRSF4 TCF7 EEF1A1 SELL CCR7 IL6R IGFBP4 IGFL2 |
| HALLMARK_EPITHELIAL_MESENCHYMAL_TRANSITION | ABI3BP ACTA2 ADAM12 ANPEP APLP1 AREG BASP1 BDNF BGN BMP1 CADM1 CALD1 CALU CAP2 CAPG CCN1 CCN2 CD44 CD59 CDH11 CDH2 CDH6 COL11A1 COL12A1 COL16A1 COL1A1 COL1A2 COL3A1 COL4A1 COL4A2 COL5A1 COL5A2 COL5A3 COL6A2 COL6A3 COL7A1 COL8A2 COLGALT1 COMP COPA CRLF1 CTHRC1 CXCL1 CXCL12 CXCL6 CXCL8 DAB2 DCN DKK1 DPYSL3 DST ECM1 ECM2 EDIL3 EFEMP2 ELN EMP3 ENO2 FAP FAS FBLN1 FBLN2 FBLN5 FBN1 FBN2 FERMT2 FGF2 FLNA FMOD FN1 FOXC2 FSTL1 FSTL3 FUCA1 FZD8 GADD45A GADD45B GAS1 GEM GJA1 GLIPR1 GPC1 GPX7 GREM1 HTRA1 ID2 IGFBP2 IGFBP3 IGFBP4 IL15 IL32 IL6 INHBA ITGA2 ITGA5 ITGAV ITGB1 ITGB3 ITGB5 JUN LAMA1 LAMA2 LAMA3 LAMC1 LAMC2 LGALS1 LOX LOXL1 LOXL2 LRP1 LRRC15 LUM MAGEE1 MATN2 MATN3 MCM7 MEST MFAP5 MGP MMP1 MMP14 MMP2 MMP3 MSX1 MXRA5 MYL9 MYLK NID2 NNMT NOTCH2 NT5E NTM OXTR P3H1 PCOLCE PCOLCE2 PDGFRB PDLIM4 PFN2 PLAUR PLOD1 PLOD2 PLOD3 PMEPA1 PMP22 POSTN PPIB PRRX1 PRSS2 PTHLH PTX3 PVR QSOX1 RGS4 RHOB SAT1 SCG2 SDC1 SDC4 SERPINE1 SERPINE2 SERPINH1 SFRP1 SFRP4 SGCB SGCD SGCG SLC6A8 SLIT2 SLIT3 SNAI2 SNTB1 SPARC SPOCK1 SPP1 TAGLN TFPI2 TGFB1 TGFBI TGFBR3 TGM2 THBS1 THBS2 THY1 TIMP1 TIMP3 TNC TNFAIP3 TNFRSF11B TNFRSF12A TPM1 TPM2 TPM4 VCAM1 VCAN VEGFA VEGFC VIM WIPF1 WNT5A |
| HALLMARK_INTERFERON_GAMMA_RESPONSE | ADAR APOL6 ARID5B ARL4A AUTS2 B2M BANK1 BATF2 BPGM BST2 BTG1 C1R C1S CASP1 CASP3 CASP4 CASP7 CASP8 CCL2 CCL5 CCL7 CD274 CD38 CD40 CD69 CD74 CD86 CDKN1A CFB CFH CIITA CMKLR1 CMPK2 CMTR1 CSF2RB CXCL10 CXCL11 CXCL9 DDX60 DHX58 EIF2AK2 EIF4E3 EPSTI1 FAS FCGR1A FGL2 FPR1 GBP4 GBP6 GCH1 GPR18 GZMA HELZ2 HERC6 HIF1A HLA-A HLA-B HLA-DMA HLA-DQA1 HLA-DRB1 HLA-G ICAM1 IDO1 IFI27 IFI30 IFI35 IFI44 IFI44L IFIH1 IFIT1 IFIT2 IFIT3 IFITM2 IFITM3 IFNAR2 IL10RA IL15 IL15RA IL18BP IL2RB IL4R IL6 IL7 IRF1 IRF2 IRF4 IRF5 IRF7 IRF8 IRF9 ISG15 ISG20 ISOC1 ITGB7 JAK2 KLRK1 LAP3 LATS2 LCP2 LGALS3BP LY6E LYSMD2 MARCHF1 METTL7B MT2A MTHFD2 MVP MX1 MX2 MYD88 NAMPT NCOA3 NFKB1 NFKBIA NLRC5 NMI NOD1 NUP93 OAS2 OAS3 OASL OGFR P2RY14 PARP12 PARP14 PDE4B PELI1 PFKP PIM1 PLA2G4A PLSCR1 PML PNP PNPT1 PSMA2 PSMA3 PSMB10 PSMB2 PSMB8 PSMB9 PSME1 PSME2 PTGS2 PTPN1 PTPN2 PTPN6 RAPGEF6 RBCK1 RIGI RIPK1 RIPK2 RNF213 RNF31 RSAD2 RTP4 SAMD9L SAMHD1 SECTM1 SELP SERPING1 SLAMF7 SLC25A28 SOCS1 SOCS3 SOD2 SP110 SPPL2A SRI SSPN ST3GAL5 ST8SIA4 STAT1 STAT2 STAT3 STAT4 TAP1 TAPBP TDRD7 TNFAIP2 TNFAIP3 TNFAIP6 TNFSF10 TOR1B TRAFD1 TRIM14 TRIM21 TRIM25 TRIM26 TXNIP UBE2L6 UPP1 USP18 VAMP5 VAMP8 VCAM1 WARS1 XAF1 XCL1 ZBP1 ZNFX1 |

Supplementary table 4: Pathway grouping for GeoMx correlation analyses

| Pathway | Pathway group |
| --- | --- |
| HALLMARK_APOPTOSIS | other |
| HALLMARK_HYPOXIA | other |
| HALLMARK_IL2_STAT5_SIGNALING | il2_jak_stat |
| HALLMARK_INFLAMMATORY_RESPONSE | other |
| HALLMARK_TNFA_SIGNALING_VIA_NFKB | tnf_nfkb |
| BIOCARTA_CTLA4_PATHWAY | texh |
| REACTOME_CTLA4_INHIBITORY_SIGNALING | texh |
| GOBP_REGULATION_OF_T_CELL_APOPTOTIC_PROCESS | other |
| KEGG_JAK_STAT_SIGNALING_PATHWAY | il2_jak_stat |
| KEGG_MTOR_SIGNALING_PATHWAY | mtor |
| BIOCARTA_IL2_PATHWAY | il2_jak_stat |
| WP_MACROPHAGE_MARKERS | other |
| COATES_MACROPHAGE_M1_VS_M2_DN | m2 |
| COATES_MACROPHAGE_M1_VS_M2_UP | m1 |
| GSE5099_CLASSICAL_M1_VS_ALTERNATIVE_M2_MACROPHAGE_DN | m2 |
| GSE5099_CLASSICAL_M1_VS_ALTERNATIVE_M2_MACROPHAGE_UP | m1 |
| BIOCARTA_MHC_PATHWAY | mhc |
| REACTOME_CLASS_I_MHC_MEDIATED_ANTIGEN_PROCESSING_PRESENTATION | mhc |
| REACTOME_MHC_CLASS_II_ANTIGEN_PRESENTATION | mhc |
| REACTOME_PD_1_SIGNALING | texh |
| TCELL_EXHAUSTION_ZHANG | texh |
| HALLMARK_EPITHELIAL_MESENCHYMAL_TRANSITION | emt |
| HALLMARK_INTERFERON_GAMMA_RESPONSE | ifng |
| PROGENY_Androgen | other |
| PROGENY_EGFR | other |
| PROGENY_Estrogen | other |
| PROGENY_Hypoxia | other |
| PROGENY_JAK.STAT | il2_jak_stat |
| PROGENY_MAPK | mapk |
| PROGENY_NFkB | tnf_nfkb |
| PROGENY_p53 | other |
| PROGENY_PI3K | mtor |
| PROGENY_TGFb | tgfb_wnt |
| PROGENY_TNFa | tnf_nfkb |
| PROGENY_Trail | other |
| PROGENY_VEGF | other |
| PROGENY_WNT | tgfb_wnt |

Supplementary table 5: Log2FC relative to non-treated control

|  | **P5** | **P10** | **T5** | **T10** | **P5T5** | **P5T10** | **P10T10** |
| --- | --- | --- | --- | --- | --- | --- | --- |
| **C659_pOme** | -0,18057 | -0,27 | 0,13 | -0,34 |  | 0,121562 | 0,106915 |
| **C973_pAdn** |  | 0,12 |  | 0,21 |  | -0,33407 | -0,16189 |
| **C817_pOme** |  | 0,53 |  | 0,196589 |  | 0,759949 | 0,545824 |
| **C181_iOme** | 0,250375 | 0,72 | 0,68 | 0,56 | 0,742771 | 0,200622 | 0,274622 |

Supplementary table 6: scRNAseq data published previously

| sample | scRNAseq_Previous_Publication_DOI |
| --- | --- |
| S001_IDS | NA |
| S002_chemo-naive | NA |
| S003_chemo-naive | NA |
| S004_chemo-naive | 10.1371/journal.pcbi.1009290 |
| S005_chemo-naive | NA |
| S006_chemo-naive | NA |
| S007_chemo-naive | NA |
| S008_IDS | NA |
| S008_chemo-naive | NA |
| S009_IDS | 10.1126/sciadv.abm1831 |
| S009_chemo-naive | 10.1126/sciadv.abm1831;10.1093/bioinformatics/btab178 |
| S010_chemo-naive | 10.1371/journal.pcbi.1009290 |
| S011_IDS | 10.1126/sciadv.abm1831 |
| S011_chemo-naive | 10.1126/sciadv.abm1831 |
| S012_IDS | 10.1126/sciadv.abm1831 |
| S012_chemo-naive | 10.1126/sciadv.abm1831 |
| S013_chemo-naive | NA |
| S014_IDS | 10.1126/sciadv.abm1831 |
| S014_chemo-naive | 10.1126/sciadv.abm1831 |
| S015_IDS | 10.1126/sciadv.abm1831 |
| S015_chemo-naive | 10.1126/sciadv.abm1831 |
| S016_chemo-naive | 10.1093/bioinformatics/btab178 |
| S017_IDS | NA |
| S017_chemo-naive | NA |
| S018_IDS | NA |
| S018_chemo-naive | NA |
| S019_IDS | NA |
| S019_chemo-naive | NA |
| S020_IDS | 10.1126/sciadv.abm1831 |
| S020_chemo-naive | 10.1126/sciadv.abm1831 |
| S021_IDS | NA |
| S021_chemo-naive | NA |
| S022_IDS | 10.1093/bioinformatics/btab178 |
| S022_chemo-naive | NA |
| S001_chemo-naive | NA |
| S024_IDS | NA |
| S025_IDS | 10.1126/sciadv.abm1831 |
| S025_chemo-naive | 10.1126/sciadv.abm1831 |
| S026_IDS | NA |
| S027_IDS | 10.1126/sciadv.abm1831 |
| S027_chemo-naive | 10.1126/sciadv.abm1831; 10.1016/j.devcel.2023.04.012 |
| S028_IDS | 10.1126/sciadv.abm1831 |
| S028_chemo-naive | 10.1126/sciadv.abm1831 |
| S029_IDS | 10.1126/sciadv.abm1831; 10.1016/j.devcel.2023.04.012 |
| S029_chemo-naive | 10.1126/sciadv.abm1831; 10.1016/j.devcel.2023.04.012 |
| S030_IDS | 10.1126/sciadv.abm1831 |
| S030_chemo-naive | 10.1126/sciadv.abm1831 |
| S031_IDS | NA |
| S031_chemo-naive | NA |
| S032_IDS | NA |
| S032_chemo-naive | NA |

Supplementary table 7: T-cell exhaustion signature used in scRNAseq data analysis

exhaustion_zhang

CXCL13

TNFRSF9

LAYN

ENTPD1

HAVCR2

CTLA4

KRT86

TNFRSF18

GEM

TIGIT

DUSP4

PHLDA1

VCAM1

SNX9

MYO7A

MYO1E

KIR2DL4

AFAP1L2

RBPJ

TNS3

ETV1

PDLIM4

PDCD1

HMOX1

CD82

AKAP5

TOX

TNFSF4

NDFIP2

LAG3

FAM3C

ZBED2

GOLIM4

MIR155HG

MS4A6A

SIRPG

ACP5

RGS1

RGS2

CD200

CD27

PDE7B

LMCD1

FKBP1A

GZMB

AHI1

TBC1D4

SNAP47

SRGAP3

Supplementary table 8: Antibodies used in the t-CycIF protocol

| Cycle | Antibody | Fluorochrome | Company | Cat number | Clone | Concentration | Role |
| --- | --- | --- | --- | --- | --- | --- | --- |
| 0 | Rabbit | Rabbit |  |  |  | 2000 |  |
| 0 | Goat | Goat |  |  |  | 2000 |  |
| 0 | Mouse | Mouse |  |  |  | 2000 |  |
| 1 | TAZ | 488 | CST | 8418 |  | 200 | DEG |
| 1 | CD207 | 555 | R&D | AF2088 |  | 400 | APC |
| 1 | SNAT1 | 647 | Millipore | N104-37 |  | 100 | glutamine metabolism |
| 2 | CD163 | 488 | Abcam | ab218293 |  | 500 | M2 Macrophges |
| 2 | CD20 | 647 | eBioscience | 50-0202-80 |  | 500 | B-cells |
| 3 | Annexin | 488 | BDBiosciences | 611838 |  | 200 | DEG |
| 3 | pSTAT1 | 555 | CST | 8183S |  | 100 | Interferon activation |
| 4 | CD4 | 488 | R&D | fab8165g |  | 150 | T-Cells |
| 4 | CD8a | eFluor 660 | eBioscience | 50-0008-80 |  | 200 | T-Cells |
| 5 | CD45RO | 488 | BioLegend | 304212 |  | 400 | Memory T-cells |
| 5 | FOXP3 | 555 | EbioSciences | 41-4777-82 |  | 200 | T-Regs |
| 5 | CD3d | 647 | Abcam | ab208514 |  | 200 | T-cells |
| 6 | TIM3 | 488 | CST | 54669S |  | 200 | ICP |
| 6 | Desmin | 647 | Abcam | ab195177 |  | 200 | DEG |
| 7 | FOXOA3 | 647 | Bioss | BS-3140R-AF647 | Ziming; 200 | 100 | DEG |
| 8 | HE4 (Rb-Z) | 488 | Abcam | ab24480 |  | Zenon Rb488 | DEG |
| 8 | CD11c | 555 | CST |  |  | 150 | Antigen presenting cells |
| 8 | yH2AX | 647 | Biolegend | 613407 |  | 200 | DNA damage |
| 9 | Ki67 | 488 | CST | 11882 |  | 300 | Proliferation |
| 9 | Vimentin | 555 | CST | 9855 |  | 100 | Stroma |
| 9 | MHCII | 647 | Abcam | ab201347 |  | 1000 | Antigen presentation |
| 10 | Lamin B1 | 488 | abcam | ab194106 |  | 300 | Segmentation |
| 10 | CK7 | 555 | Abcam | ab209601 |  | 400 | Tumor cells |
| 10 | MHCI | 647 | Abcam | 199837 |  | 2000 | Antigen presentation |
| 11 | E-cadherin | 488 | CST | 3199 |  | 400 | Tumor cells |
| 11 | SMA | 555 | abcam | ab202509 |  | 800 | Stroma |
| 11 | CD31 | 647 | Abcam | 218582 | GR3219255-2 | 500 | Blood vessels |
| 12 | IBA1 | 488 | abcam | ab195031 |  | 450 | Macrophages |

Supplementary table 9: Cell type calling logic for TRIBUS

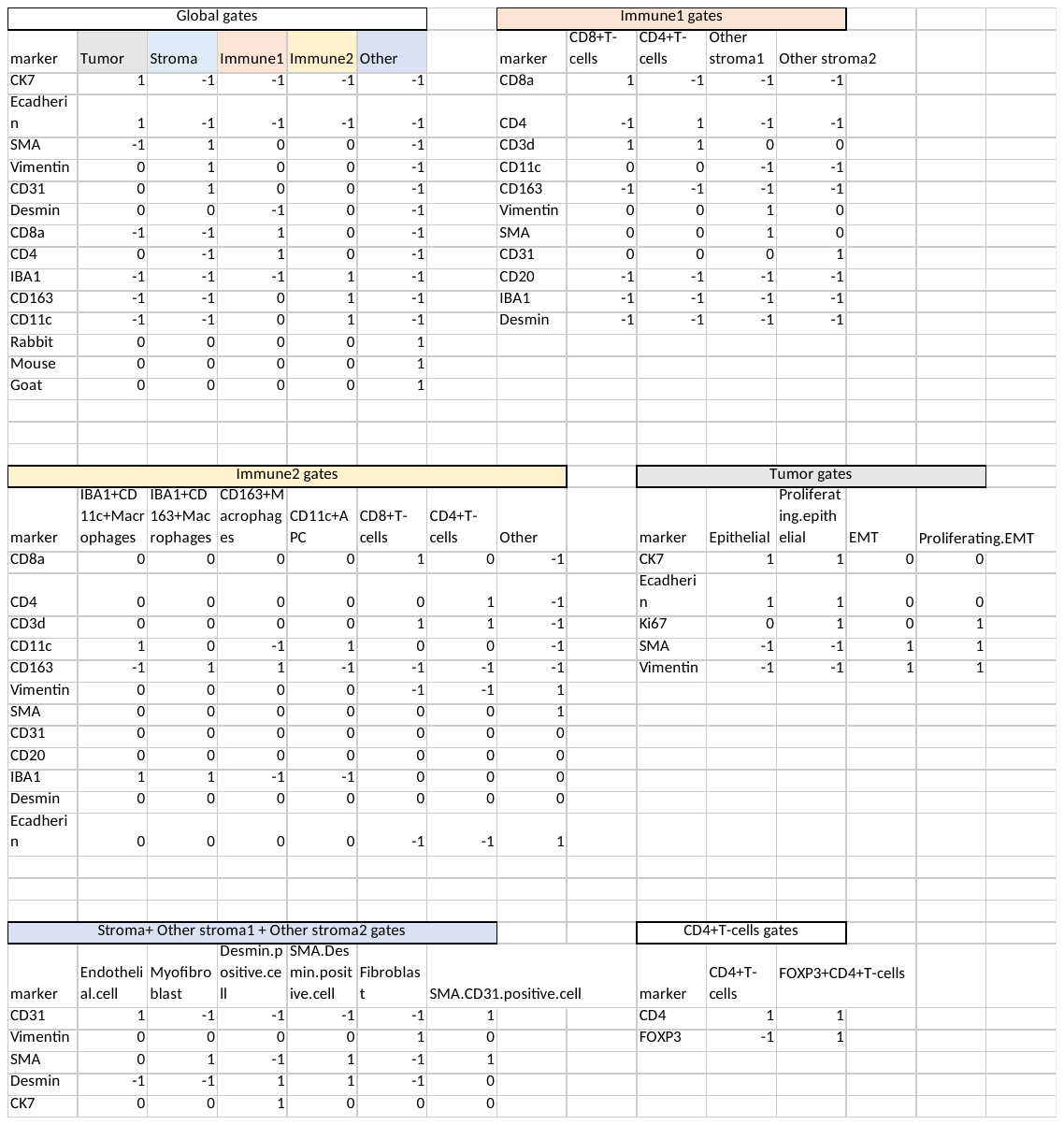

Supplementary table 10: Antibodies used in the lowplex t-CycIF experiment

| Cycle | Antibody | Fluorochrome | Company | Cat number | Clone | Concentration |
| --- | --- | --- | --- | --- | --- | --- |
| 0 | Hoechst_1 |  |  |  |  |  |
| 0 |  |  |  |  |  |  |
| 0 | NaK ATPase | 555 | Abcam | ab283345 | EP1845Y | 50 |
| 0 |  |  |  |  |  |  |
| 1 | Hoechst_2 |  |  |  |  |  |
| 1 |  |  |  |  |  |  |
| 1 | PanCK | 555 | eBioscience/Invitrogen | 41-9003-82 | AE1/AE3 | 200 |
| 1 | CD45 | 647 | Biolegend | 304018 | HI30 | 100 |
| 2 | Hoechst_3 |  |  |  |  |  |
| 2 | IBA1 | 488 | Abcam | ab195031 | EPR136(2) | 100 |
| 2 | Vimentin | 555 | CST | 98555 | D21H3 | 100 |
| 2 | CD8 | 647 | eBioscience/Invitrogen | 50-0008-80 | AMC908 | 50 |

Supplementary table 11: Antibody details for multiparametric flowcytometry

| Antibody | Host species | Clone | Reference |
| --- | --- | --- | --- |
| LIVE/DEAD Fixable Aqua Dead Cell Stain | - | - | Invitrogen #L34965 |
| FITC/CD56 | Mouse | MEM-188 | ImmunoTools, #21270563 |
| PE/CD14 | Mouse | 18D11 | ImmunoTools, #21620144 |
| PE-CF594/CD4 | Mouse | RPA-T4 | BD Biosciences, #562281 |
| PerCP-CY5.5/HLA-DR | Mouse | G46-6 | BD Pharmingen™, #552764 |
| PE-CY7/Ki-67 | Mouse | B56 | BD Biosciences, #561283 |
| APC/Granzyme B | Mouse | GB11 | Invitrogen, #GRB05 |
| AF700/IFNg | Mouse | B27 | BD Biosciences, #557995 |
| APC-Fire 750/EpCAM | Mouse | 9C4 | BioLegend, #324233 |
| BV421/CD45 | Mouse | HI30 | BD Horizon™, #563880 |
| BV605/CD8 | Mouse | SK1 | BD Biosciences, #564116 |
| BV711/CD11c | Mouse | B-ly6 | BD Horizon™, #563130 |
| BV786/CD3 | Mouse | SK7 | BD Horizon™, #563800 |
